# A nucleoside signal generated by a fungal endophyte regulates host cell death and promotes root colonization

**DOI:** 10.1101/2022.03.11.483938

**Authors:** Nick Dunken, Heidi Widmer, Gerd U. Balcke, Henryk Straube, Gregor Langen, Nyasha M. Charura, Pia Saake, Concetta De Quattro, Jonas Schön, Hanna Rövenich, Stephan Wawra, Armin Djamei, Matias D. Zurbriggen, Alain Tissier, Claus-Peter Witte, Alga Zuccaro

## Abstract

The intracellular colonization of plant roots by the beneficial fungal endophyte *Serendipita indica* follows a biphasic strategy. After an early biotrophic phase, the interaction transitions to a host cell death phase restricted to the epidermal and cortex layers of the root. Host cell death contributes to the successful accommodation of the fungus during the beneficial interaction in *Arabidopsis thaliana*. How host cell death is initiated and controlled is largely unknown. Here we show that two fungal enzymes, the ecto-5’-nucleotidase *Si*E5NT and the nuclease *Si*NucA, act synergistically in the plant apoplast at the onset of cell death to produce deoxyadenosine (dAdo), a potent cell death inducer in animal systems. The uptake of extracellular dAdo, but not the structurally related adenosine (Ado), activates a previously undescribed cell death mechanism in *A. thaliana*. Mutation of the equilibrative nucleoside transporter ENT3 in *A. thaliana* results in resistance to cell death triggered by extracellular dAdo and reduced fungal-mediated cell death during root colonization. A library screen of *A. thaliana* T-DNA insertion lines identified a toll/interleukin-1 receptor nucleotide-binding leucine-rich repeat (TIR-NLR) protein as an additional intracellular component in dAdo-triggered cell death. Mutation of this previously uncharacterised TIR-NLR, which we have named ISI (<u>i</u>nduced by *<u>S</u>. <u>i</u>ndica*), affects host cell death, fungal colonization and growth promotion, suggesting a key role in the regulation of root cell death and plant-microbe interaction. Our data show that the combined activity of two fungal apoplastic enzymes leads to the production of a metabolite that, upon uptake, triggers TIR-NLR-modulated plant cell death, providing a link to immunometabolism in plants.

**Short summary:** Efficient intraradical colonization by the beneficial fungal endophyte *Serendipita indica* requires restricted host cell death. How this symbiotic host cell death is initiated and controlled is largely unknown. Here we show that two fungal enzymes, the ecto-5’-nucleotidase *Si*E5NT and the nuclease *Si*NucA, act synergistically in the apoplast at the onset of cell death to produce deoxyadenosine (dAdo), a potent cell death inducer in animal systems. Uptake of extracellular dAdo activates a previously undescribed cell death mechanism in plants. Mutation of the *A. thaliana* equilibrative nucleoside transporter *ENT3* leads to resistance to cell death triggered by uptake of extracellular dAdo and to reduced fungal-mediated cell death during colonization. A library screen of *A. thaliana* T-DNA insertion lines identified a TIR-NLR protein as an additional intracellular component in dAdo-triggered cell death, providing a link to immunometabolism in plants.

**In a nutshell:** Regulated host cell death is part of the plant defense strategy against pathogens, but it is also involved in the accommodation of certain beneficial microbes in the roots. We have identified extracellular metabolites and intracellular metabolic signals that contribute to colonization by beneficial root fungal endophytes and uncovered a conserved cell death mechanism likely co-opted for establishing plant-endophyte symbiosis.

## Introduction

Regulated cell death (RCD) occurs in plants as part of normal growth and development and in response to abiotic and biotic stimuli. In plant-microbe interactions, host cell death programs can mediate either resistance or successful infection. Depending on the type of microbial lifestyle (biotrophic or necrotrophic), host cell death can benefit the plant by stopping the growth of biotrophs, or the microbe by promoting the growth of necrotrophs. Thus, control of plant host cell death is critical to the outcome of an interaction. Host cell death also plays a critical role in certain beneficial interactions, challenging the paradigm that cell death in plant-microbe interactions implies pathogenesis or host-microbe incompatibility. Both symbiosis with beneficial microbes and infection by pathogens require sophisticated control of host defenses and nutrient fluxes. Certain features of the interaction of beneficial microbes with plants, such as affecting host immunity, metabolism, and host cell death, are reminiscent of pathogen infections. In Rhizobium-legume symbioses, root nodules are formed to provide a niche for bacterial nitrogen fixation. The formation of infection pockets in some of these symbioses is associated with host cell death and the production of hydrogen peroxide (D’Haeze et al., 2003). Host cell death is also observed in ectomycorrhizal symbioses (Mucha et al., 2014; Ragnelli et al., 2014) and is a requisite for the establishment of symbiotic interactions with the widely distributed beneficial fungi of the order Sebacinales (Deshmukh et al., 2006; Lahrmann et al., 2013; Qiang et al., 2012). Molecular environmental studies have shown that some of the most abundant taxa of this order have little, if any, host specificity and interact with a wide variety of plant species. Sebacinales isolates, including *S. indica* and *S. vermifera*, exhibit beneficial effects such as growth promotion, increased seed production, and protection from pathogens, and thus play an important role in natural and managed ecosystems (Oberwinkler et al., 2013; Tedersoo et al., 2014; Weiss et al., 2016). The requirement of restricted host cell death for the establishment of certain beneficial microorganisms leads to the hypothesis that the activation of cell death mechanisms in roots has a more important ecological function than previously thought.

In the hosts *Hordeum vulgare* (hereafter barley) and *A. thaliana* (hereafter Arabidopsis), fungi of the order Sebacinales initially colonize living cells that die during the progression of colonization (Deshmukh *et al*., 2006; Lahrmann *et al*., 2013; Lahrmann et al., 2015; Lahrmann and Zuccaro, 2012; Qiang *et al*., 2012; Zuccaro et al., 2011). This symbiotic cell death is thought to contribute to niche differentiation during microbial competition for space and nutrients in the root and appears to be restricted to colonized cells in the epidermis and outer cortex. The host pathways that control the induction and execution of plant cell death and the fungal elicitors/effectors that initiate this process in roots are still largely unknown (Lahrmann *et al*., 2013; Nizam et al., 2019; Qiang *et al*., 2012; Schneider and Lynch, 2018). Pathogenic and beneficial fungi have a large repertoire of secreted effectors that can affect host cell physiology and suppress plant defenses, promoting fungal colonization. In fungi, effectors have been described mainly in biotrophic and hemibiotrophic foliar pathogens (Lo Presti et al., 2015). In contrast, only a few effectors of root symbiotic fungi have been functionally characterized (Kloppholz et al., 2011; Nizam *et al*., 2019; Nostadt et al., 2020; Plett et al., 2014; Voß et al., 2018; Wawra et al., 2016). Therefore, the modes of action of effectors of mutualistic fungi remain poorly understood.

Using genomics, transcriptomics, and proteomics, we identified proteins secreted by *S. indica* in the root apoplast, the space outside the plasma membrane, including cell walls, where material can freely diffuse (Nizam *et al*., 2019). One secreted fungal protein consistently found at various stages of symbiosis is the ecto-5’-nucleotidase *Si*E5NT (PIIN_01005) (Nizam *et al*., 2019). Expression of *SiE5NT is* induced during colonization of barley and Arabidopsis roots, but not in axenic fungal culture (Nizam *et al*., 2019). Animal ecto-5’-nucleotidases play a key role in the conversion of AMP to adenosine, counteracting the immunogenic effects of extracellular adenosine 5’-triphosphate (eATP) released from stimulated host cells (Antonioli et al., 2013). Extracellular ATP is an important signaling molecule in plants that controls development and response to biotic and abiotic stresses. In Arabidopsis, eATP mediates various cellular processes through its binding to the purinergic membrane-associated receptor proteins DORN1/P2K1 and the recently described P2K2 (Pham et al., 2020). In the apoplast, ATP accumulation increases cytoplasmic calcium and triggers a defense response against invading microbes. The perception of extracellular nucleotides, such as eATP, plays an important role in plant-fungal interactions – we recently demonstrated this by colonization experiments with the knockout (KO) mutant *dorn1* of Arabidopsis, which is better colonized by *S. indica. Si*E5NT is able to hydrolyze ATP, ADP, and AMP to adenosine and phosphate, which alters the eATP content in the apoplast and the plant response to fungal colonization (Nizam et al., 2019). Secretion of *Si*E5NT in Arabidopsis leads to enhanced colonization by *S. indica*, confirming its role as an apoplastic effector protein. Considering the important role *Si*E5NT plays in fungal accommodation at early symbiotic stages, we proposed that modulation of extracellular nucleotide levels and their perception play a key role in compatibility during early plant-fungal interactions in roots (Nizam *et al*., 2019). Secreted *Si*E5NT homologs are also present in fungal pathogens such as *Colletotrichium incanum* and *Fusarium oxysporum* and other Arabidopsis endophytes such as *Colletotrichum tofieldiae*, where their expression is induced during colonization, suggesting that purine-based extracellular biomolecules also play a role in other plant-fungal interactions.

At the onset of cell death, a small fungal endonuclease, which we named *Si*NucA (PIIN_02121), is secreted simultaneously with *Si*E5NT during colonization of barley and Arabidopsis by *S. indica* (Nizam *et al*., 2019; Thürich et al., 2018). In plants, the mechanisms linking immune recognition of DNA danger signals in the extracellular environment to innate signaling pathways in the cytosol are poorly understood, as is the role of (deoxy)nucleotide metabolism in root colonization and cell death. Here we show that the synergistic activity of *Si*NucA and *Si*E5NT leads to the production of deoxyadenosine (dAdo) and, in smaller amounts, other deoxynucleosides from extracellular DNA (eDNA). eDNA can be released from roots during microbial colonization in a process called root extracellular trap (RET) formation, or from dying cells and from microorganisms during biofilm development on the root surface (Chambard et al., 2021; Driouich et al., 2019; Tran et al., 2016; Wen et al., 2009). The production of dAdo by fungal extracellular enzymes is similar to the processes involved in dAdo-mediated immune cell death of *Staphylococcus aureus* in animals, which appears to ensure bacterial survival in host tissues (Thammavongsa et al., 2013; Winstel et al., 2018). Staphylococcal nuclease and adenosine synthase A (AdsA, a homolog of *Si*E5NT) are both required to release dAdo from neutrophil extracellular traps (NETs), which has a potent cytotoxic effect on macrophages and other immune cells (Thammavongsa *et al*., 2013).

Here we show that extracellular dAdo, but not the structurally similar Ado, activates a previously unidentified cell death mechanism in plants. Expression of either extracellular *Si*NucA or *Si*E5NT *in planta* results in enhanced colonization by *S. indica* and host cell death (Nizam *et al*., 2019). We found that a mutation in the equilibrative nucleoside transporter 3 (ENT3) of Arabidopsis leads to a strong resistance phenotype to dAdo-induced cell death but not to methyl jasmonate, a cell death inducer of senescence. Accordingly, compared with the WT line colonized by *S. indica*, the *ent3* KO line shows less fungal-induced cell death in the root and accumulates less of the extracellular signaling metabolite methylerythritol cyclodiphosphate (MEcPP) in response to fungal colonization or dAdo treatment. Finally, in a mutant screen of Arabidopsis T-DNA insertion lines, we found that a previously uncharacterized TIR-NLR protein is involved in modulating cell death triggered by dAdo as well as fungal colonization and growth promotion. Most characterized TIR-NLRs (TNLs) act as sensors of effectors secreted by pathogens. The involvement of a TNL protein in the modulation of cell death by a symbiotically produced metabolite establishes a connection to immunometabolism in plants. Therefore, we propose that hydrolysis of extracellular metabolites by the fungal-derived *Si*NucA and *Si*E5NT provides a direct link between purine metabolism, immunity, and cell death programs in roots (Figure 1).

**Figure 1.**
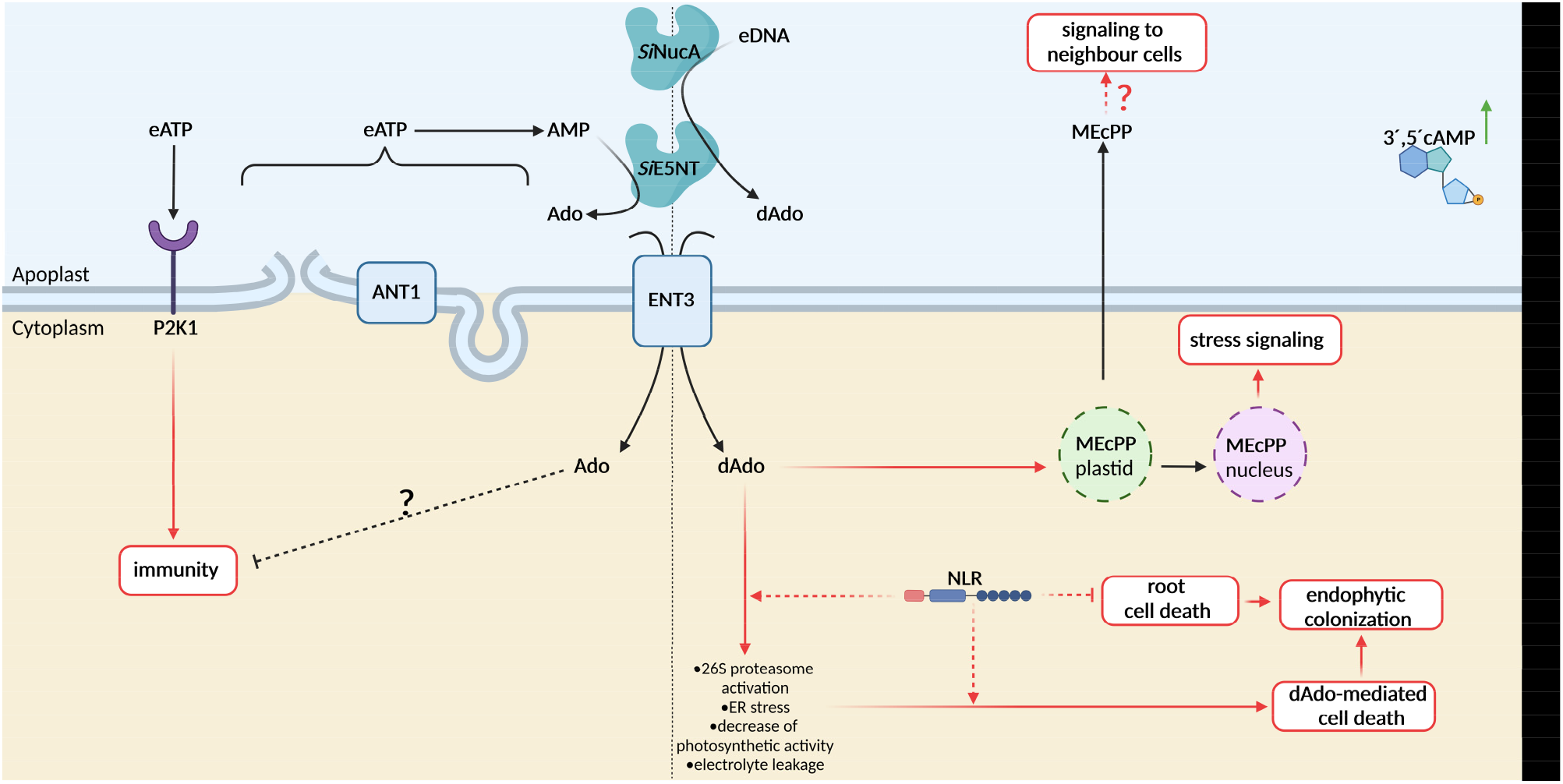
Current model for dAdo-triggered cell death during colonization of *A. thaliana* roots by *S. indica*. The beneficial endophyte *S. indica* secretes two enzymes into the apoplast, the nuclease *Si*NucA and the ecto-nucleotidase *Si*E5NT. *Si*E5NT is initially involved in manipulating eATP signaling (left panel), as described in Nizam et al. (2019). *SiNucA* accumulates at the onset of cell death. The combined activity of *SiNucA* and *SiE5NT* releases deoxynucleosides from DNA, with a strong preference for dAdo, a potent cell death inducer in animal systems. dAdo is transported to the cytoplasm *via AtENT3*, where it triggers a cell death process and contributes to successful fungal colonization. dAdo induces the production of the retrograde stress signal MEcPP from plastids, activating stress signaling and possibly intercellular communication. Cell death triggered by dAdo is modulated by an uncharacterized TIR-type NLR protein (ISI, <u>i</u>nduced by *<u>S</u>. <u>i</u>ndica*), providing a link to immunometabolism.

## Results

### Fading of host nuclei and vacuolar collapse are hallmarks of symbiotic cell death during *S. indica* root colonization

*S. indica* induces restricted cell death in colonized root cells of Arabidopsis and barley, resulting in characteristic cytological features at later stages of colonization. In these two hosts, the timing and extent of cell death differs (Lahrmann *et al*., 2013; Qiang *et al*., 2012). In Arabidopsis, the cell death phenotype is less pronounced than in barley, but cytological analyses showed fading of host nuclei, vacuolar collapse (Figure 2A-M), and swelling of the endoplasmic reticulum (ER) in colonized cells (Qiang *et al*., 2012) indicating ER stress and host cell death during root colonization. To characterize the timing of fungal-induced cell death, the presence and shape of plant nuclei during colonization by *S. indica* were monitored by confocal laser scanning microscopy using either the nucleic acid dye DAPI (4′,6-diamidino-2′-phenylindole dihydrochloride) and the cell death dye SYTOX Orange (Figure 2C-I) or an Arabidopsis line expressing the nuclear marker H2B:mCherry (Figure 2A, B). Nuclei stained with SYTOX Orange were visible 7 to 8 days post inoculation (dpi), indicating that at this time the plasma and nuclear membranes of Arabidopsis were permeable to the dye, which is a hallmark of cell death (Figure 2F, G). In the Arabidopsis H2B:mCherry line, plant nuclei in the epidermal layer were often elongated, faded, and eventually disappeared by 8 to 10 dpi in heavily colonized areas of the root (Figure 2A, B). In dying cells, *S. indica* hyphae embedded in plant nuclei were frequently observed, indicating that the fungus can digest and feed on host nuclear DNA (Figure 2C-E, H, I). Cytological analyses show that Arabidopsis root cell death begins around 7 dpi and by 10 dpi most host nuclei have faded or disappeared in cells colonized by *S. indica* under the growth condition tested.

**Figure 2.**
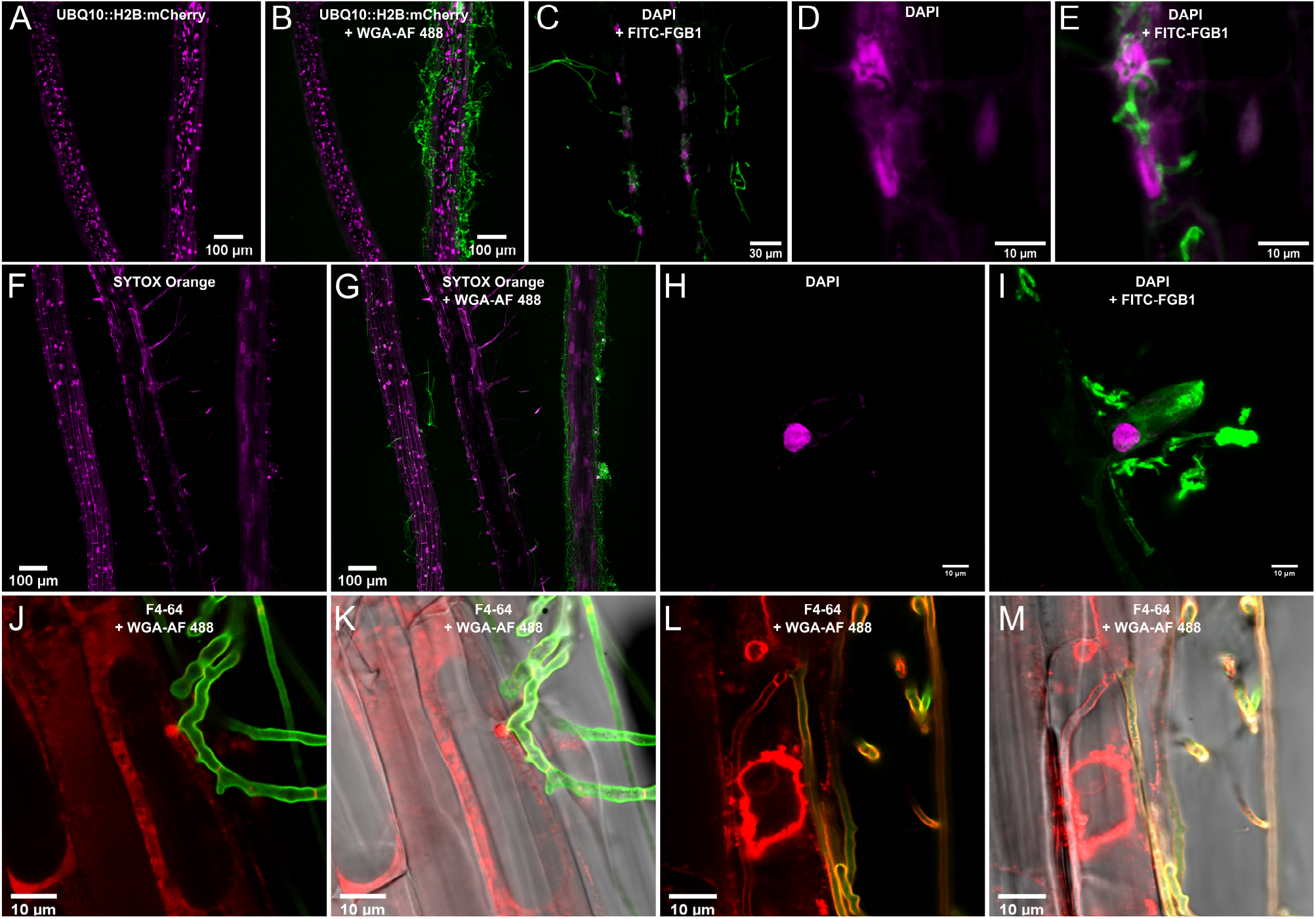
Host cell death during *S. indica* colonization of roots. (A and B) Arabidopsis roots expressing the fluorescent nuclear marker UBQ10::H2B:mCherry (magenta) stained with the fungal cell wall marker Wheat Germ Agglutinin Alexa Fluor 488 Conjugate WGA-AF 488 (green) at 10 dpi. (C) Plant nuclei stained with DAPI (magenta) and fungal cell wall and matrix stained with the β-glucan binding lectin FGB1-FITC 488 (green) at 6 dpi. As colonization *by S. indica* progresses, host nuclei often become elongated and fade. (D and E) Close-up of *S. indica* hyphae embedded in a host nucleus from (C). (F and G) Staining of nucleic acids of roots colonized with *S. indica* with the dead cell indicator (membrane integrity marker) SYTOX Orange (magenta) and fungal hyphae with WGA-AF 488 (green) at 10 dpi. (H and I) *S. indica* hyphae fluorescently labeled with the β-glucan-binding lectin FGB1-FITC 488 (green) embedded in a DAPI-stained host nucleus (magenta) at 6dpi. (J-M) Progressive vacuolar collapse of colonized root cells. Fungal hyphae are stained with WGA-AF 488 (green), while membranes are stained with FM4-64 (red). (J and K) Initial biotrophic colonization. (L and M) Vacuolar collapse in a dying host cell. CLSM was repeated at least five times with 3 to 4 plants colonized by *S. indica*. Fading of nuclei at the onset of cell death during fungal colonization was regularly observed.

### *Si*NucA and *Si*E5NT act synergistically in the production of deoxynucleosides

Although the interaction between *S. indica* and plant roots has been extensively studied, comparatively little is known about the contribution of apoplastic effectors to fungal accommodation or the mechanism of cell death in this system (Nizam *et al*., 2019; Nostadt *et al*., 2020; Rafiqi et al., 2013; Wawra *et al*., 2016). We previously analyzed soluble apoplastic proteins in barley at different stages of *S. indica* colonization. We found that *Si*E5NT was one of the predominant fungal proteins consistently present in this extracellular compartment (Nizam *et al*., 2019). We demonstrated that *Si*E5NT functions as a membrane-bound nucleotidase that is released into the apoplast during host colonization. *Si*E5NT is capable of releasing phosphate and adenosine from ATP, ADP, and AMP (Nizam *et al*., 2019). In addition, the small secreted protein *Si*NucA with predicted endonuclease activity (Figure S1) was found in the apoplastic fluid of colonized barley roots at the onset of cell death (5 dpi) (Nizam *et al*., 2019). *Si*NucA is also secreted during root colonization in Arabidopsis (Thürich *et al*., 2018). *Si*NucA expression is transiently induced during cell death in barley and Arabidopsis compared with axenic fungal growth, as shown by transcriptomic data (Lahrmann *et al*., 2013) and quantitative PCR analyses (Figure 3A). This prompted us to further investigate the involvement of this small secreted protein in fungal colonization and host cell death. The secretion of *Si*NucA and its enzymatic activity were investigated by overexpressing a *SiNucA*:HA:His construct in *S. indica* (Figure 3B, S2). Supernatants from fungal OE strains, but not from empty vector control strains, and affinity-purified *Si*NucA were able to degrade single- and double-stranded DNA and RNA from fungal and plant material, indicating nonspecific nuclease activity (Figure 3C). Addition of magnesium and calcium increased *Si*NucA activity in *in vitro* assays, whereas EDTA inhibited it (Figure S3).

**Figure 3.**
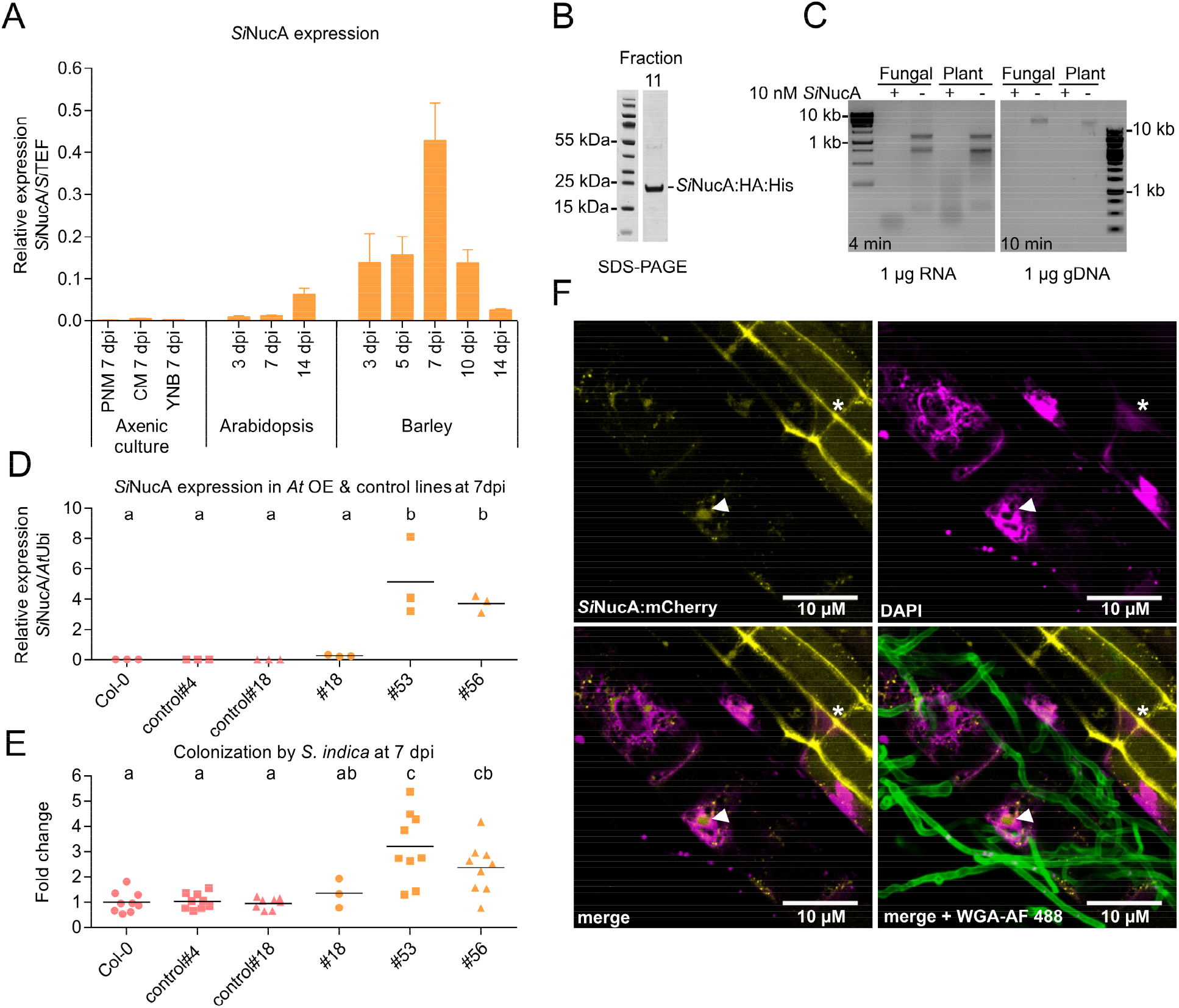
*Si*NucA is a small secreted nuclease involved in fungal accommodation. (A) *Si*NucA expression in Arabidopsis and barley roots colonized by *S. indica* and axenic cultures measured by the 2^−ΔCT^ method. Error bars: SD, n = 3 (biological replicates). (B) *SiNucA*:HA:His protein enrichment from culture filtrate precipitated with 80% ammonium sulfate and separated by size exclusion chromatography. The resulting fraction was separated by SDS-PAGE and *SiNucA:*HA:His was stained with Coomassie Brilliant Blue. (C) The purified *SiNucA*:HA:His protein was incubated with fungal (*S. indica)* or plant (Arabidopsis) RNA or DNA in 5 mM Tris buffer (pH = 8) containing 1 mM MgCl_2_ and 1 mM CaCl_2_ for 4 (RNA) or 10 (DNA) min and visualized after gel electrophoresis. (D) Arabidopsis lines expressing *SiNucA* driven by the 35S promoter (lines 18, 53, and 56) compared with control lines (4 and 18: segregating from T2 generation and Col-0 WT). Roots of plants grown on ½ MS medium were inoculated with *S. indica* and analyzed after 7 dpi. The dots represent independent biological replicates, and the lines represent the mean. Different letters indicate significantly different groups as determined by one-way ANOVA with post-hoc Tukey HSD test (p<0.05). (E) Root colonization by *S. indica* in transgenic 35S::*Si*NucA-Arabidopsis lines at 7 dpi was assessed by RT-qPCR by comparing expression of the fungal housekeeping gene *SiTEF* and the plant gene *AtUbi* and the 2^−ΔΔCT^ method, normalized to colonization in the WT Col-0. The dots represent independent biological replicates, while the lines represent the mean. Different letters indicate significantly different groups as determined by one-way ANOVA with post-hoc Tukey HSD test (p<0.05). (F) CLSM live cell images of an Arabidopsis root expressing *Si*NucA:mCherry (yellow) and inoculated with *S. indica*. Fungal cell walls are stained with WGA-AF 488 (green) and nuclei with DAPI (magenta). *Si*NucA:mCherry fluorenscence signal accumulates in the apoplast/cell periphery in non-colonized root cells (asterisk) and (re)localizes in host nuclei in colonized cells (arrow).

To investigate the effects of *Si*NucA on colonization, we tested independent homozygous T3 lines heterologously expressing a native version of *SiNucA* in Arabidopsis under control of the 35S promoter. As controls, we used WT and segregating lines lacking the *SiNucA* gene. Expression of *SiNucA* in Arabidopsis resulted in higher fungal colonization at 7 dpi, which correlated with the *SiNucA* expression level (Figure 3D, E), demonstrating its importance in fungal accommodation. Overcolonization resulted in a reduction in plant biomass that was not observed in the mock-treated *SiNucA* expression lines (Figure S4). Localization studies by confocal microscopy of Arabidopsis roots expressing either full-length *Si*NucA or a version lacking the N-terminal signal peptide (SP) fused to mCherry confirmed secretion of the full-length *Si*NucA fusion protein into the apoplast and functionality of the SP. After plasmolysis with 1M NaCl or sorbitol, the mCherry fluorescence signal was visible on the cell walls and apparently on the membrane of the shrinking cells for the full-length *Si*NucA fusion protein, but not for the cytoplasmic version without SP (Figure S5). Remarkably, the presence of *S. indica* resulted in relocalization of the full-length mCherry-tagged *Si*NucA to host nuclei only in cells colonized by the fungus and not in bystander cells, where the mCherry signal remained at the cell periphery even after plasmolysis (Figure 3F and S5, S6). The presence of *Si*NucA in the apoplast (Nizam *et al*., 2019) and in the nucleus of colonized host cells (Figure 3F) suggests that this small secreted protein can be redirected to the nucleus during colonization and can act on both eDNA and nuclear DNA.

The co-occurrence of *Si*NucA and *Si*E5NT in the apoplast at 5 dpi led us to speculate that these two enzymes might cooperate in promoting fungal colonization of roots. Using *Si*E5NT affinity-purified from leaves of *Nicotiana benthamiana* that transiently expressed the enzyme as a secreted C-terminal Strep-tagged variant, we found activity with dAMP and AMP (Figure 4A), whereas no activity was detected for other deoxyribonucleotides (dGMP, dCMP, dTMP) or ribonucleotides (GMP, UMP, 3′,5′-cAMP) tested. Moreover, dATP and the general phosphatase substrate *para*-nitrophenyl pyrophosphate (*p*NPP) were not *Si*E5NT substrates. Interestingly, only dAMP was hydrolyzed at a constant rate, whereas the AMP hydrolysis rate gradually decreased under the selected reaction conditions (Figure S7A, B). The K_M_ for AMP (15.9 µM) was ∼20-fold lower than for dAMP (361.6 µM), but the k_cat_ for dAMP (11.9 s^−1^) exceeded that of AMP (1.1 s^−1^) by a factor of ∼10 (Figure 4A, B). These data suggest that *Si*E5NT hydrolyzes AMP slightly better at low substrate concentrations (below 20 µM), but is far more efficient for dAMP at higher substrate concentrations that might prevail in a nucleus undergoing degradation and in the case of eDNA degradation. To test the ability of *Si*NucA and *Si*E5NT to work together, we incubated DNA with *Si*E5NT and *Si*NucA alone and in combination. As expected, *Si*NucA alone degraded DNA (Figure 4C), but neither deoxyribonucleotides nor deoxynucleosides were detected as reaction products by LC-MS analysis, suggesting that *Si*NucA degrades DNA to oligonucleotides. *Si*E5NT alone also did not release deoxyribonucleotides from DNA, but produced small amounts of deoxyribonucleosides (Figure 4D). However, in combination, *Si*NucA and *Si*E5NT released deoxyribonucleosides from DNA with a strong preference for dAdo. This demonstrates the ability of *Si*E5NT to use oligonucleotides released by *Si*NucA as a substrate and act synergistically with *Si*NucA to preferentially release dAdo from DNA (Figure S7C).

**Figure 4.**
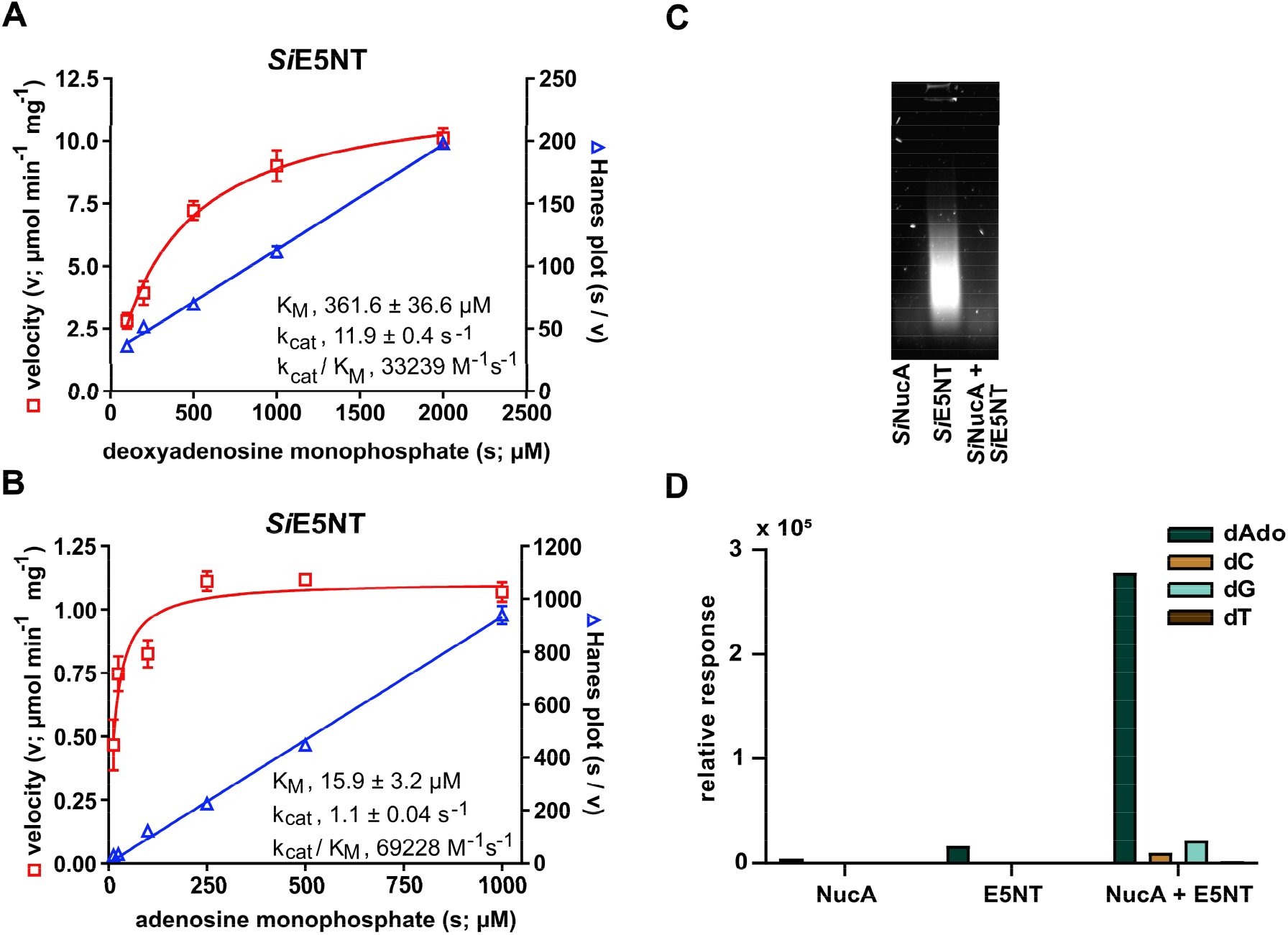
*Si*NucA and *Si*E5NT act synergistically in the production of deoxy nucleosides from DNA. (A) Enzymatic activity and kinetic constants of *Si*E5NT. Left axis, initial catalytic velocity (v) for phosphate production at different dAMP concentrations fitted with the Michaelis-Menten equation. Right axis, ratio of dAMP concentrations and velocities (s/v) plotted against dAMP concentrations (s) fitted by linear regression (Hanes plot). Error bars: SD, n = 3. (B) As in (A), but using AMP as substrate (n = 3). (C) Degradation of DNA by *Si*NucA and/or *Si*E5NT. 2% agarose gel loaded with the products of a one-hour incubation of 10 μg salmon sperm DNA at 25° C with different enzyme combinations. (D) Relative quantification of dAdo (deoxyadenosine), dC (deoxycytidine), dG (deoxyguanosine) and dT (deoxythymidine) released during incubation in (C) by HPLC-MS/MS. Deoxynucleotides could not be detected in any of the reactions.

### dAdo induces host cell death

The production of dAdo by the synergistic activity of the two secreted fungal enzymes resembles the processes involved in dAdo-mediated immune cell death of *Staphylococcus aureus* in animal cells, which ensures the exclusion of macrophages from the center of abscesses where the bacteria survive (Thammavongsa *et al*., 2013; Winstel *et al*., 2018). This motivated us to test the effect of dAdo in plants. Incubation of the H2B-mCherry line of Arabidopsis with extracellular dAdo, but not with Ado or buffer, resulted in the fading and disappearance of nuclei in roots within 48 hours (Figure 5A). This root cell death phenotype could be quantified using Evans blue azo dye (Figure 5B, C). In addition, extracellular dAdo-triggered hallmarks of cell death, such as increased electrolyte leakage, induction of cell death marker gene expression as well as activation of the 26S proteasome and decreased photosynthetic activity (F_V_/F_M_) (Figure 5D-H, S8). The effects were concentration-dependent, and removal of dAdo from the culture supernatant 24 hours after treatment resulted in recovery of Arabidopsis seedlings, indicating that activation of the cell death program in Arabidopsis is still reversible at this stage (Figure S9, S10).

**Figure 5.**
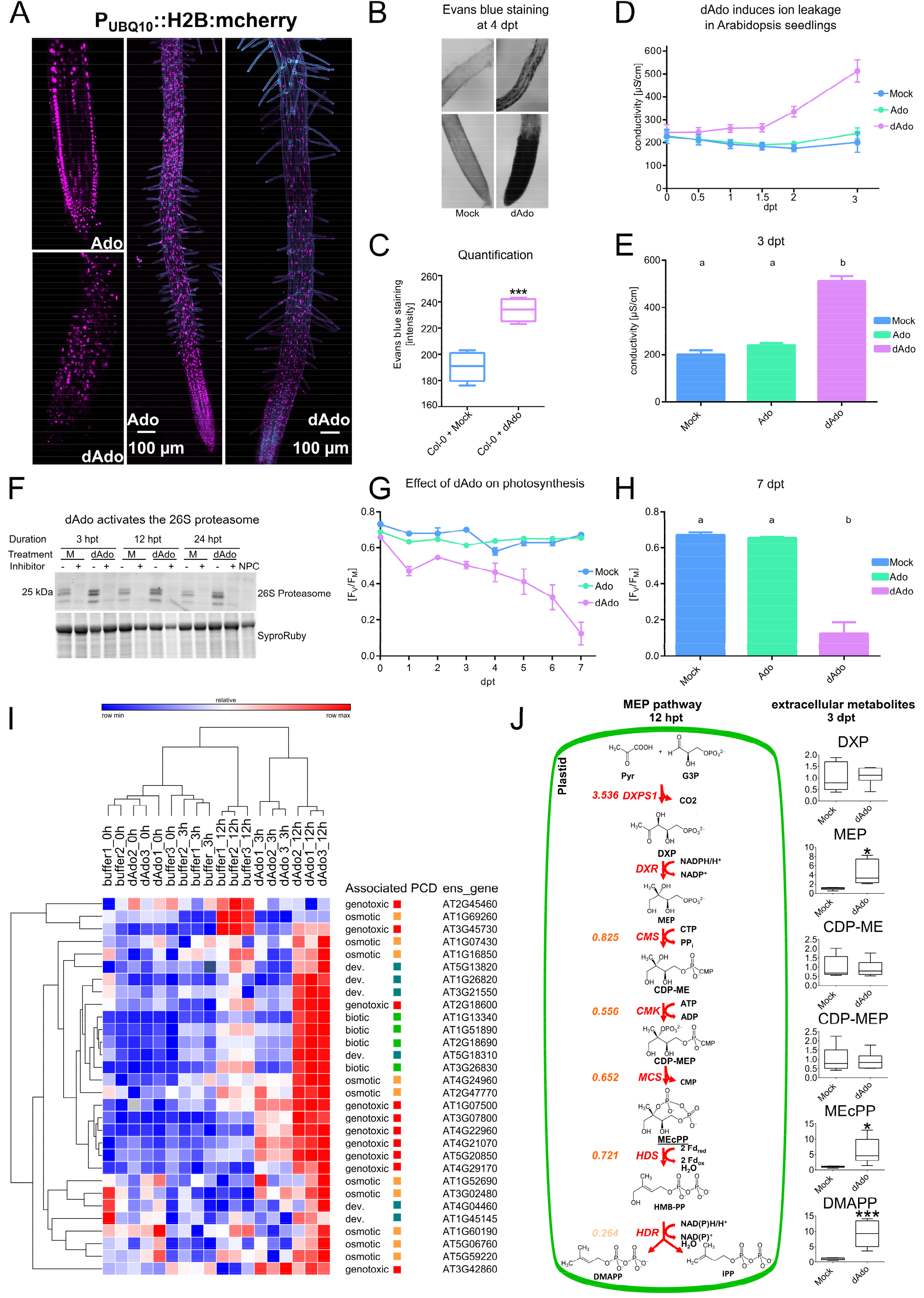
dAdo induces cell death *in planta*. (A) CLSM images of seven-day-old Arabidopsis roots expressing the nuclear marker UBQ10::H2B:mCherry. Incubation with dAdo, but not Ado (500 µM), results in disorganization of Arabidopsis cells in the root tip and disappearance or fading of nuclear material. (B) Bright field microscopy of the root tip and differentiation zone of Arabidopsis seedlings treated with mock/dAdo (500 µM) and stained with Evans blue cell death dye. (C) Quantification of root cell death in Col-0 root tips 4 days after dAdo treatment. Cell death was assessed by Evans blue staining. Boxplots show data from 5 biological replicates. Asterisks represent a significant difference from the mock-treated samples analyzed by Student’s t-test (p < 0.005 ***). The experiment was independently repeated 3 times with similar results. (D) Electrolyte leakage of Col-0 seedlings (9 days old) in MES buffer after mock, ado or dAdo treatment. Error bars show the standard error of the mean (SEM) from 6 biological replicates. The experiment was independently repeated at least 3 times with similar results. (E) Electrolyte leakage at 3 dpt of seedlings from (D). Error bars show the standard error of the mean (SEM) of 6 biological replicates. Different letters indicate significantly different groups as determined by one-way ANOVA with post-hoc Tukey HSD test (p<0.05). The experiment was independently repeated at least 3 times with similar results. (F) Activity of the 26S proteasome of Arabidopsis. Total protein extracts from 14-day-old seedlings treated with 500 µM dAdo or 2.5 mM MES buffer (M) were incubated with 1 µM of probe MVB072 (Kolodziejek et al. 2011). Prior to labeling, samples were incubated with 50 µM of the proteasome inhibitor epoximycin (+) or DMSO (-). Samples were then labeled for 2 hours and separated by SDS-PAGE. 26S proteasome activity was visualized by fluorescence scanning. SYPRO^™^ Ruby staining was performed to compare sample amounts. A non-probe control (NPC) consisting of a mixture of all samples incubated with DMSO was used as an additional control. The experiment was repeated twice with similar results. (G) Photosynthetic activity (F_V_ /F_M_) of 7-day-old Col-0 seedlings incubated with 500 µM Ado or dAdo. Error bars represent the standard error of the mean (SEM) obtained from twelve biological replicates. The experiment was repeated three times with similar results. (H) Photosynthetic activity (F_V_ /F_M_) of 7-day-old Col-0 seedlings incubated with 500 µM Ado or dAdo at 7 dpt from (G). Error bars represent the standard error of the mean (SEM) obtained from twelve technical replicates. Different letters indicate significantly different groups as determined by one-way ANOVA with post-hoc Tukey HSD test (p<0.05). (I) Heatmap of core regulated cell death (RCD) marker-genes expression (selected from Olvera-Carrillo et al., 2015) of 7-day-old Arabidopsis seedlings after dAdo treatment at 0, 3, and 12 hours. RNA-Seq data are presented as log_2_ (tpm) (Table S6). Heat maps and hierarchical clustering (one-minus Pearson correlation) were generated using Morpheus, https://software.broadinstitute.org/morpheus. (J) Graphical representation of the methyl erythritol 4-phosphate (MEP) pathway. The colored numbers on the left represent the log_2_ fold-change after 12 hours of dAdo treatment (Padj < 0.05) as measured by RNA-Seq. The boxplots on the right show the corresponding extracellular metabolites at 3 dpt. Asterisks represent significant differences from the mock-treated sample analyzed by Student’s t-test (p < 0.05 * ; p < 0.001 ***).

A cell death phenotype was also observed in young leaves of *N. benthamiana* during expression by *Agrobacterium tumefaciens* infiltration of a full length *Si*E5NT construct with the signal peptide for secretion into the apoplastic space (Figure S11, S12) and in seedlings incubated with dAdo (Figure S13). Cell death was not visible in older leaves or in leaves of *N. benthamiana* expressing the control vector p19 or *Si*NucA alone. The observed phenotype suggests that the presence of *Si*E5NT is sufficient to trigger cell death in this plant host upon wounding/damage by agroinfiltration, which could release DNA and DNAses into the apoplast or elicit a response to the presence of the bacterium or its proteins (Figure S11, S12). To test whether cell death triggered by dAdo is conserved in basal plant lineages, we additionally tested its activity in the liverwort *Marchantia polymorpha*. Incubation with dAdo also induced cell death in this plant species (Figure S14). Overall, these data suggest that the dAdo-mediated cell death mechanism is conserved across plant lineages.

### dAdo induces the MEP pathway and accumulation of the stress signaling metabolite MEcPP

To investigate the mechanism by which dAdo triggers cell death in plants and to determine whether it activates stress signaling pathways, we analyzed the transcriptional response of Arabidopsis to this metabolite at 0, 3, and 12 h compared to a buffer control using RNA-seq. Most marker genes for regulated cell death (Olvera-Carrillo et al., 2015) were upregulated at 12 hours (Figure 5I & Table S1-3). In addition, the plastidial 2-C-methyl-D-erythritol-4-phosphate (MEP) pathway was induced upon incubation with dAdo after 12 hours in Arabidopsis seedlings (Figure 5J & Table S4). Accordingly, MEcPP, a precursor of plastidial isoprenoids and a stress-specific retrograde signaling metabolite produced by the MEP pathway, accumulated extracellularly 3 days after treatment as measured by LC-MS/MS (Figure 5J). Abiotic stress and wounding increase levels of cytoplasmic MEcPP, which coordinates stress response pathways in plants (Xiao et al., 2012). The detection of this isoprenoid intermediate in the extracellular environment after dAdo treatment suggests that this metabolite may serve as a stress signal in bystander cells, a putative important function that needs further investigation. Incubation of Arabidopsis with extracellular MEcPP did not result in cell death, demonstrating that accumulation of this metabolite is not sufficient to trigger cell death and therefore is not the cause of the observed dAdo-mediated cell death (Figure S15).

### dAdo-triggered signaling and cell death are not mediated by canonical pattern-triggered immune responses

To determine whether cell death triggered by dAdo is mediated by signals generated by an immune receptor at the cell surface, we tested the ability of this metabolite to trigger a rapid response by monitoring calcium influx and ROS production. Both responses are part of PTI (pattern-triggered immunity), a process activated by recognition of MAMPs (microbe-associated molecular patterns) or DAMPs (damage-associated molecular patterns) by pattern-recognition receptors (PRRs) at the plasma membrane (Ngou et al., 2021; Yuan et al., 2021; Zipfel and Oldroyd, 2017).

Incubation of Arabidopsis seedlings with dAdo did not elicit calcium influx, whereas treatment with ATP or ADP, which have been previously described as DAMPs (Choi et al. 2014; Tanaka et al. 2014) but also with dATP or dADP triggered a rapid calcium influx that was dependent on the extracellular ATP receptor DORN1 (Figure 6 A-C). Treatment with dAdo or with any of the other purine derivatives did not induce a ROS burst (Figure S16). On the other hand, incubation with dAdo and to a lesser extent with ATP or dAMP, but not Ado, resulted in accumulation of the marker metabolite MEcPP and induction of the DAMP/MAMP- and fungus-responsive gene AT1G58420 at 3 dpt (Figure 6D, E & Table S5) (Choi et al., 2014; Kilian et al., 2007; Nizam *et al*., 2019). Overall, these data suggest that dAdo does not act as a typical extracellular DAMP or MAMP but induces a signaling pathway independent of calcium influx and ROS production.

**Figure 6.**
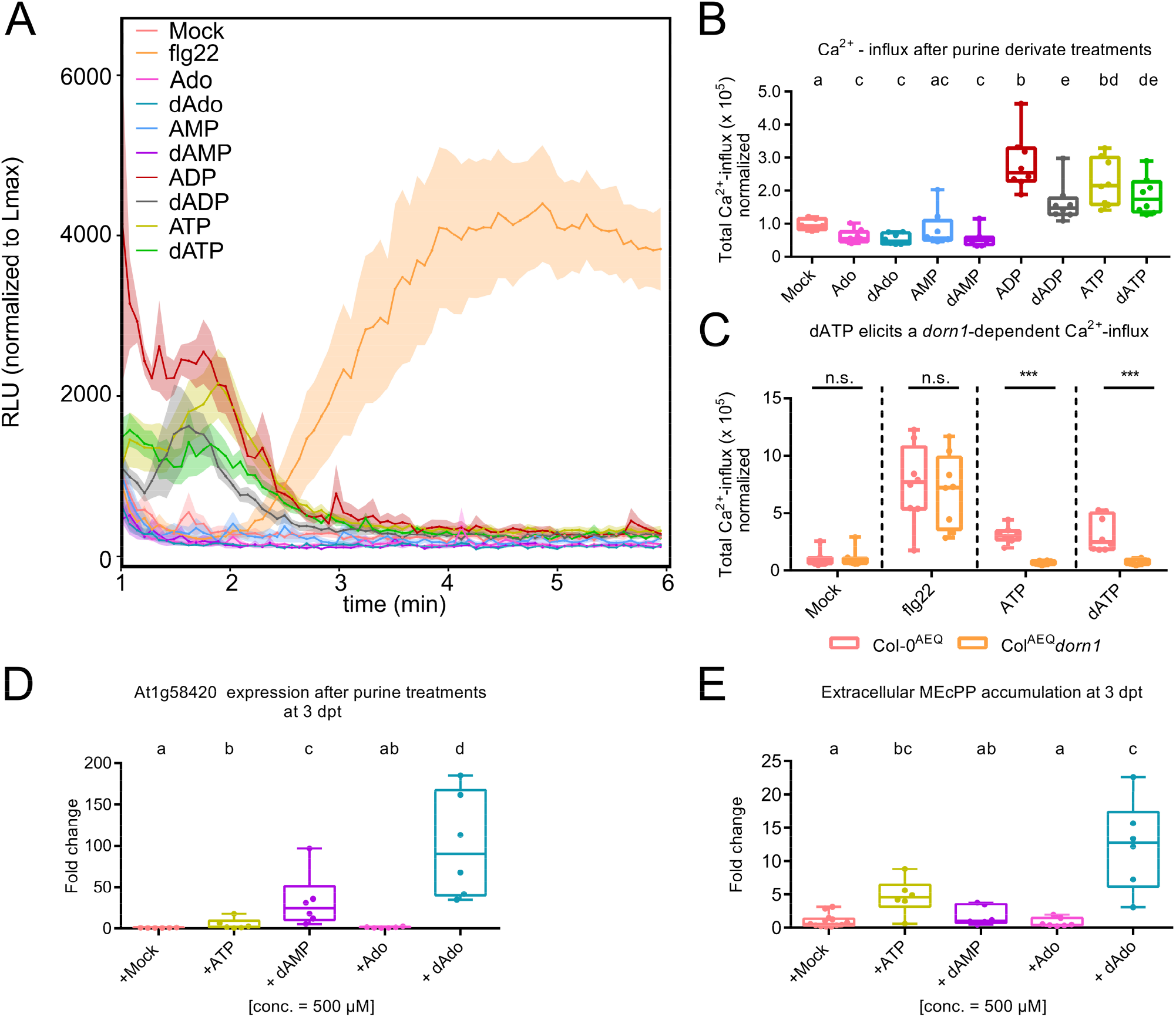
dAdo-triggered signaling and cell death are not mediated by canonical pattern-triggered immune responses. (A) Ca^2+^ influx after treatment of 7-day-old Arabidopsis Col-0^AEQ^ seedlings with 500 µM purine derivatives dissolved in 2.5 mM MES buffer (pH 5.7). Buffer and 33 µM flg22 were used as positive and negative controls, respectively. Ca^2+^ influx was monitored using a coelenterazine-based chemiluminescence assay and measured as relative light units (RLU). After discharge of the remaining aequorin by addition of CaCl_2_, the discharge kinetics were integrated and normalized to the maximum Ca^2+^ level. The discharge integral was then used to normalize the kinetics of Ca^2+^ in response to elicitor treatment. The curves represent eight biological replicates, and the experiments were additionally repeated three times independently with similar results. (B) Boxplots show the total Ca^2+^ influx over the measured time period. Calculations were performed as described in (A). Values are means ± SEM of eight biological replicates, each consisting of one seedling. Different letters indicate significant differences as determined by a Kruskal-Wallis test and Dunn’s post-hoc test (p < 0.05). The experiment was repeated three times with similar results. (C) Comparison of total Ca^2+^ influx in 7-day-old Arabidopsis Col-0^AEQ^ and Arabidopsis Col-0^AEQ^ *dorn1* seedlings after treatment with 500 µM ATP, dATP, 33 µM flg22, or 2.5 mM MES pH 5.7 (mock). Data represent values from eight biological replicates. Calculations were performed as described in (A). Asterisks indicate significant differences between the two genotypes analyzed by Student’s t-test (p < 0.005 ***). The experiment was repeated three times with similar results. (D) Expression of the marker gene AT1G58420, which responds to wounding and *S*. indica colonization, in treated 7-day-old Col-0 seedlings. Fold change expression was calculated in comparison to the housekeeping gene *AtUbi* and mock treatment using method 2^−ΔΔCt^. Data represent 6 independent biological replicates. Different letters indicate significant differences between groups as determined by Kruskal-Wallis test and post-hoc Wilcox BH adjustment (p < 0.05). (E) Metabolic analysis of supernatant from 7-day-old purine-treated Col-0 seedlings at 3 dpt. Points represent data from at least 6 biologically independent replicates normalized to the average of the corresponding mock samples. Different letters indicate significantly different groups as determined by a Kruskal-Wallis test and post-hoc Wilcox BH adjustment (p < 0.05).

### The Arabidopsis transporter ENT3 is required for dAdo-mediated signaling and cell death

The fact that we did not observe a canonical PTI response to dAdo led us to speculate that uptake of this metabolite is necessary to promote plant signaling and trigger cell death. In the animal system, treatment with extracellular dAdo leads to the accumulation of intracellular dATP, which appears to impair DNA synthesis and induces apoptosis via activation of caspase 3 (Winstel *et al*., 2018). Rapidly dividing cells are particularly susceptible to cell death triggered by dAdo and it has been shown that the toxic effect of dAdo in the animal system depends on dAdo uptake by the human equilibrative nucleoside transporter 1 (hENT1) (Winstel *et al*., 2018). Similiarly, we observed high sensitivity of dividing cells in root tips using Evans blue staining and young *N. benthamiana* plants (Figure 5 A,B; S11-13). On the contrary, *S. indica* was not sensitive to dAdo and showed normal growth ratio even at high concentrations (Figure S17). In Arabidopsis, eight potential ENT family members are annotated in the genome. Two of them are expressed in roots, namely ENT3, which is localized at the plasma membrane, and ENT1, which is localized at the tonoplast. The Arabidopsis KO line *ent3* showed a strong resistance phenotype to dAdo-induced cell death compared with WT and *ent1* KO but is unaffected in the response to methyl jasmonate-induced cell death (positive control) (Figure 7 A-C; S18). ENT3 has been shown to transport adenosine and uridine with high affinity and to be inhibited in competitive experiments by a number of different purine and pyrimidine nucleosides and 2’-deoxynucleosides, including dAdo (Li et al., 2003). This suggests that ENT3 has a broad substrate specificity and is a strong candidate for uptake of extracellular dAdo. In a competition assay, addition of extracellular Ado decreased the cell death phenotype induced by dAdo, suggesting that these two extracellular metabolites compete for the same transporter at the cell membrane (Figure 7D). Interestingly, *ENT3* is more highly expressed in the epidermis than in the rest of the root (Rich-Griffin et al., 2020), suggesting that dAdo-induced cell death may be cell type specific to some extent. This is consistent with the phenotype of root cell death observed after dAdo treatment with Evans blue staining (Figure 5B, C). Taken together, these data indicate that a functional ENT3 plays an important role in dAdo-triggered cell death in Arabidopsis, most likely by importing dAdo into the cytoplasm, where it activates signaling leading to cell death. Accordingly, the *ent3* KO line accumulates lower levels of extracellular signaling metabolites such as MEcPP, GSSG, and 3′,5′-cAMP in response to dAdo treatment at 3 dpt compared with the WT line (Figure 7E, Table S6).

**Figure 7.**
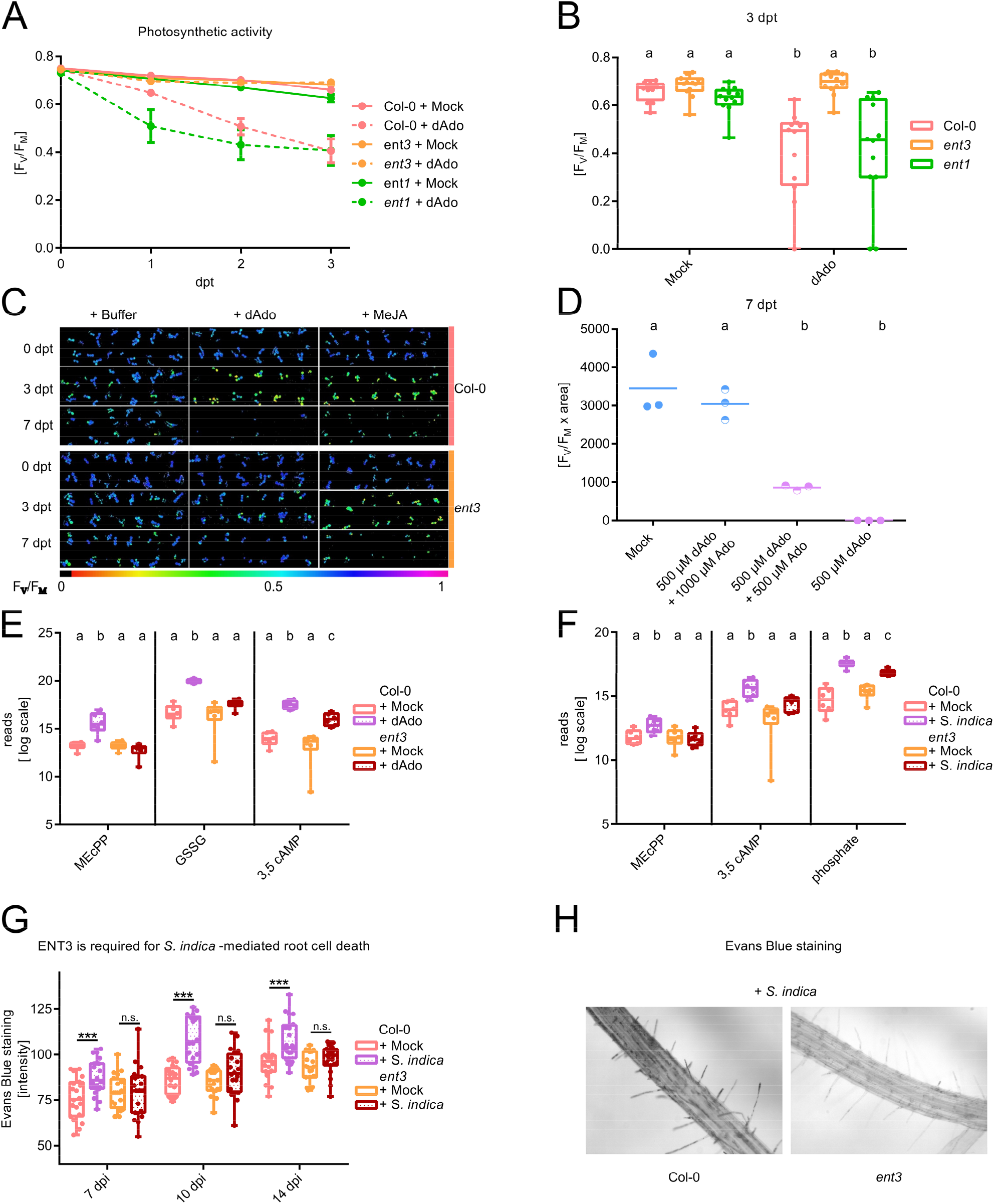
The equilibrative nucleoside transporter ENT3 of Arabidopsis is required for dAdo-mediated signal transduction and cell death. (A) Photosynthetic activity (F_V_ /F_M_) of 9-day-old mock- and dAdo-treated (500 µM) Col-0, *ent3*, and *ent1* seedlings. Measurements were taken 3 days after treatment (dpt) every 24 hours. Data show the mean and error bars show the standard error of the mean (SEM) obtained from 12 technical replicates with 3 seedlings each. The experiment was repeated more than three times independently with similar results. (B) F_V_ /F_M_ of seedlings from (A) at 3 dpt. Boxplots show data from 12 technical replicates with 3 seedlings each. Different letters indicate significant differences as determined by one-way ANOVA with post-hoc Tukey HSD test (p < 0.05). (C) Visualization of F_V_/F_M_ measured by PAM fluorometry. The FV/FM value is visualized by the color scale shown below. Shown are 12 wells with 9-day-old seedlings probed at 0, 3, and 7 dpt with either mock treatment (2.5 mM MES buffer pH 5.6), 500 µM dAdo, or 500 µM MeJA. The experiment was repeated more than three times independently with similar results. (D) Photosynthetic activity (F_V_/F_M_ x photosynthetically active area) of 9-day-old Col-0 seedlings incubated with different concentrations of Ado and dAdo at 7 dpt. Dots represent 3 biologically independent replicates consisting of 12 wells with 3 seedlings each. Different letters indicate significant differences as determined by one-way ANOVA with post-hoc Tukey HSD test (p < 0.05). (E) Metabolic analysis of supernatants from 7-day-old Col-0 and ent3 seedlings treated with dAdo at 3 dpt. Boxplots show data from 6 biologically independent replicates. Data are plotted on a log2 scale. Different letters indicate significant differences in measurements of a metabolite as determined by two-way ANOVA with post-hoc Tukey HSD test (p < 0.05). (F) Metabolic analysis of the supernatant of 7-day-old Col-0 and *ent3* seedlings *inoculated with S. indica* at different time points after treatment (3′,5′-cAMP: 3 dpi; MEcPP: 6 dpi; phosphate: 10 dpi). Boxplots show data from 6 biologically independent replicates. Data are plotted on a log2 scale. Different letters indicate significant differences in measurements of a metabolite using a two-way ANOVA with post-hoc Tukey HSD test (p < 0.05). (G) Quantification of root cell death in 7-day-old Col-0 and *ent3* roots *colonized with S. indica* at 7, 10, and 14 dpi. Cell death was assessed by Evans blue staining. Boxplots show data from 20 biological replicates. Asterisks represent significant differences analyzed by Student’s t-test (p < 0.05 *, p < 0.01 **, p < 0.005 ***). The experiment was repeated independently three times with similar results. (H) Evans blue staining of *S. indica-colonized* Col-0 and ent3 roots at 10 dpi. (I) Abundance of *S. indica* in 7-day-old Arabidopsis seedlings. Fungal (*SiTEF*) to plant (*AtUbi)* ratios were calculated using cDNA as template and the 2^− ΔCT^ method. Boxplots represent 6 independent biological replicates. Asterisks indicate a significant difference from Col-0 samples (Student’s t-test, p < 0.05 *).

### Mutation of *ENT3* impairs *S. indica*-mediated cell death

Next, we investigated whether ENT3 plays *a* role in fungal accommodation and cell death mediated by *S. indica* in roots. The *ent3* KO Arabidopsis line showed significantly less cell death upon colonization by *S. indica* compared with the WT line at 7, 10, and 14 dpi (Figure 7G, H). In addition, we observed a transient effect on fungal colonization at 8 dpi, where the *ent3* KO line was less colonized by *S. indica* compared with the WT control, as determined by qRT-PCR (Figure S19).

After colonization with *S. indica*, we also observed a transient accumulation of the extracellular signaling metabolites 3′,5′-cAMP at 3 dpi (early biotrophic phase) and MEcPP at 6 dpi (onset of cell death). In addition, a higher level of free phosphate was observed at 10 dpi (Figure 7F, Table S6). Consistent with the decreased cell death phenotype in the *ent3* KO line, the amount of these metabolites was lower in this line when colonized by *S. indica* compared with the WT line, suggesting that ENT3 is important for fungal-mediated signal transduction and cell death in Arabidopsis. Modification of extracellular metabolites levels by the activity of the fungal-derived enzymes *Si*NucA and *Si*E5NT and the host transporter ENT3 provides, for the first time, a direct link between purine metabolism, immunity, and cell death in roots. How the metabolic state of the host affects *S. indica*-induced cell death remains to be thoroughly elucidated.

### A TIR-NLR protein is involved in dAdo-mediated cell death

To identify downstream genetic determinants associated with dAdo-mediated cell death in plants, we performed a mutant screen of 6868 SALK-Arabidopsis T-DNA insertion lines (Figure S20). Sensitivity to dAdo was tested using Arabidopsis Col-0 WT as control and the SALK mutant lines grown for 14 days on solid media (1/2 MS) in 24-well plates with and without 500 µM dAdo. The dAdo-insensitive lines (survivors) from the screening were then analyzed by PAM fluorometry in three independent biological replicates together with the Col-0 WT line as control. Thirteen lines with varying degrees of dAdo resistance were identified (Table S7). One of the resistant SALK lines had a mutation at the AT5G45240 locus (SALK_034517C), which encodes a predicted TIR domain nucleotide-binding, leucine-rich repeat receptor (TIR-NLR). Mutation in this gene resulted in reproducible and significantly high resistance to dAdo-induced cell death, as evidenced by reduced electrolyte leakage, higher photosynthetic activity, and increased germination rate compared to wild-type Col-0 after incubation with dAdo (Figure 8A-F). Complementation by a cell-based transient expression system with the full-length TIR domain restored sensitivity to dAdo in a concentration-dependent manner, suggesting that the TIR domain is involved in mediating dAdo-induced cell death in Arabidopsis (Figure 8E). The TIR-NLR gene AT5G45240 is located in close proximity to *RPS4* and *RRS1* and is part of a larger locus containing multiple TNLs genes, most of which are functionally uncharacterized (Figure S21). Four of the five TNLs genes at this locus showed transiently increased expression during cell death associated with colonization by *S. indica*, with the AT5G45240 gene displaying the strongest relative induction (Figure S22). We therefore named this gene ISI, <u>i</u>nduced by *<u>S</u>. <u>i</u>ndica*. Induction could be detected especially during the onset of the cell death-associated phase (Figure 9A). While the *isi* KO line showed less cell death after dAdo treatment (Figure S23), there was increased cell death in the older parts of the mock-treated roots and during colonization with *S. indica* compared to the WT line (Figure 9B), which correlated with significantly greater fungal colonization (Figure 9C).. In addition, colonized *isi* KO seedlings did not show *S. indica*-mediated promotion of root growth, as observed in WT and *ent3* seedlings (Figure 9D and S24). These results could be explained by the activation of an alternative cell death pathway in the absence of a functional ISI TIR-NLR, leading to over-colonization by *S. indica* and loss of growth promotion (Figure 9E). Overall, our data suggest that the ISI TIR-NLR protein is involved in modulating different cell death programs in roots and mediating *S. indica* growth promotion. We further analysed the expression of both TIR- and CC-NLRs in the roots of Arabidopsis during *S. indica* colonization using RNA-seq data from different time points. We detected an induction of expression for *ISI* and also for previously functionally characterized NLRs such as *ZAR1* during beneficial colonization (Figure 9F and S25). Overall, the role of a TIR-NLR in dAdo-induced cell death suggests that this cell death is regulated by the plant immune response and is not a consequence of cytoplasmic toxicity, as assumed for animal cells.

**Figure 8.**
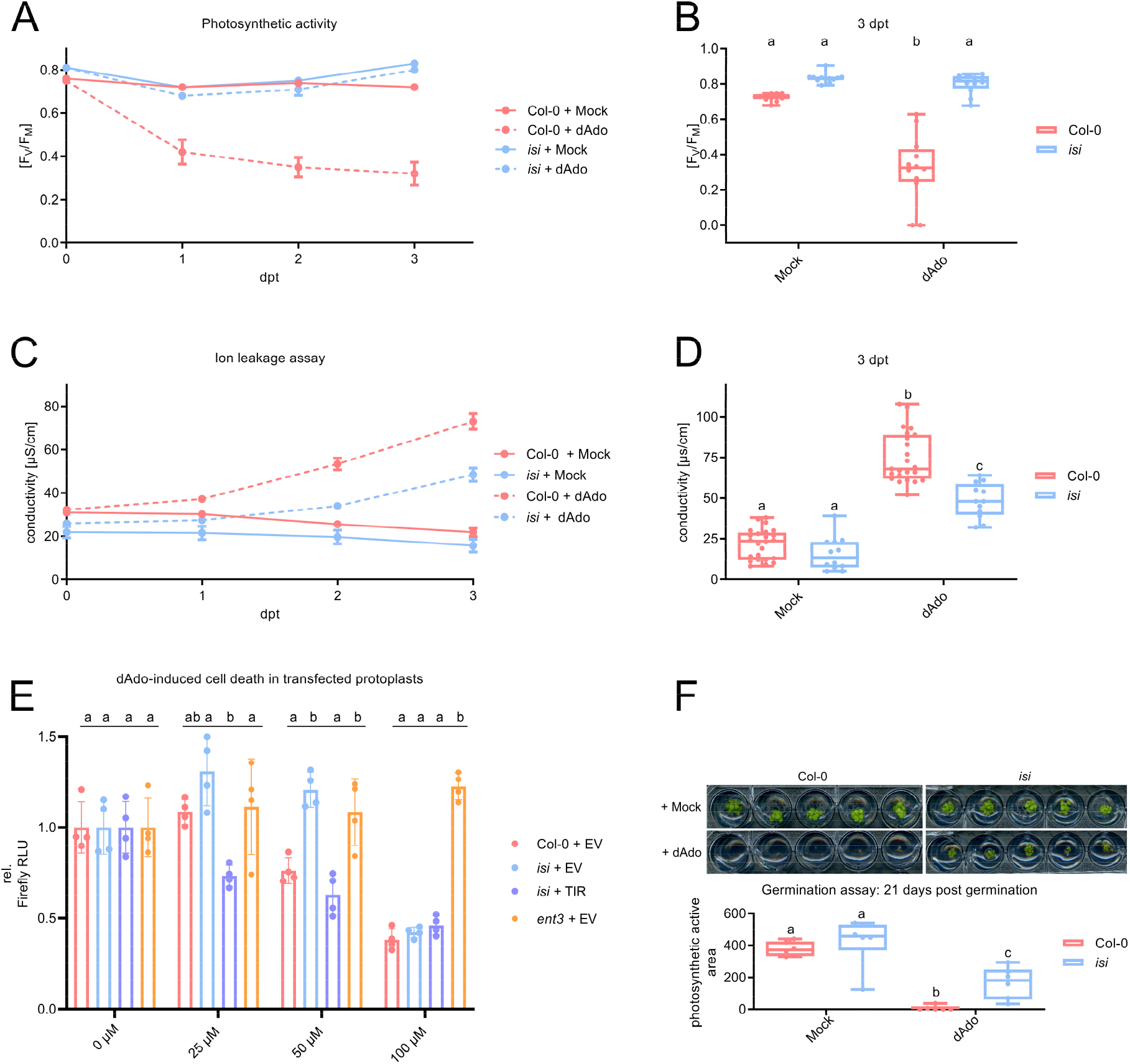
The *At*TIR-NLR AT5G45240 is involved in dAdo-mediated cell death. (A) F_V_ /F_M_ of 9-day-old mock- and dAdo-treated (500 µM) Col-0 and *isi* (AT5G45240 knockout mutant) seedlings. Measurements were performed over 3 days. Data show the mean and error bars show the standard error of the mean (SEM) obtained from 12 technical replicates with 3 seedlings each. The experiment was repeated three times independently with similar results. (B) Boxplots of the measurements of F_V_ /F_M_ from (A) at 3 dpt. Different letters indicate significant differences as determined by two-way ANOVA and post-hoc Tukey HSD test (p < 0.05). (C) Conductivity of water containing 9-day-old Col-0 or *isi* seedlings 0 - 3 days after mock or dAdo treatment (500 µM). Data points represent the mean, while error bars show the standard error of the mean (SEM) from 12 biological replicates. (D) Boxplots of conductivity measurements from (C) at 3 dpt. Different letters indicate significant differences as determined by two-way ANOVA with post-hoc Tukey HSD test (p < 0.05). (E) Relative Luciferase activity of *A. thaliana* protoplasts transfected with Luciferase and either a construct expressing the TIR domain of ISI or an empty vector. Values were normalized to mock treatment (0 µM dAdo). Different letters indicate significantly different groups as determined by one-way ANOVA with post-hoc Tukey HSD test (p<0.05) comparing all protoplast of treatment with one concentration. (F) Germination of Col-0 WT and *isi* seedlings on 1/10 PNM medium containing 500 µM dAdo or MES. The upper half shows photographs of the seedlings after 21 days. The lower half shows the quantification of F_V_ /F_M_ measured by PAM fluorometry. Boxplots show F_V_ /F_M_ measurements per well, containing six seedlings. Different letters indicate significant differences as determined by two-way ANOVA with post-hoc Tukey HSD test (p < 0.05). The experiment was repeated three times independently with similar results.

**Figure 9.**
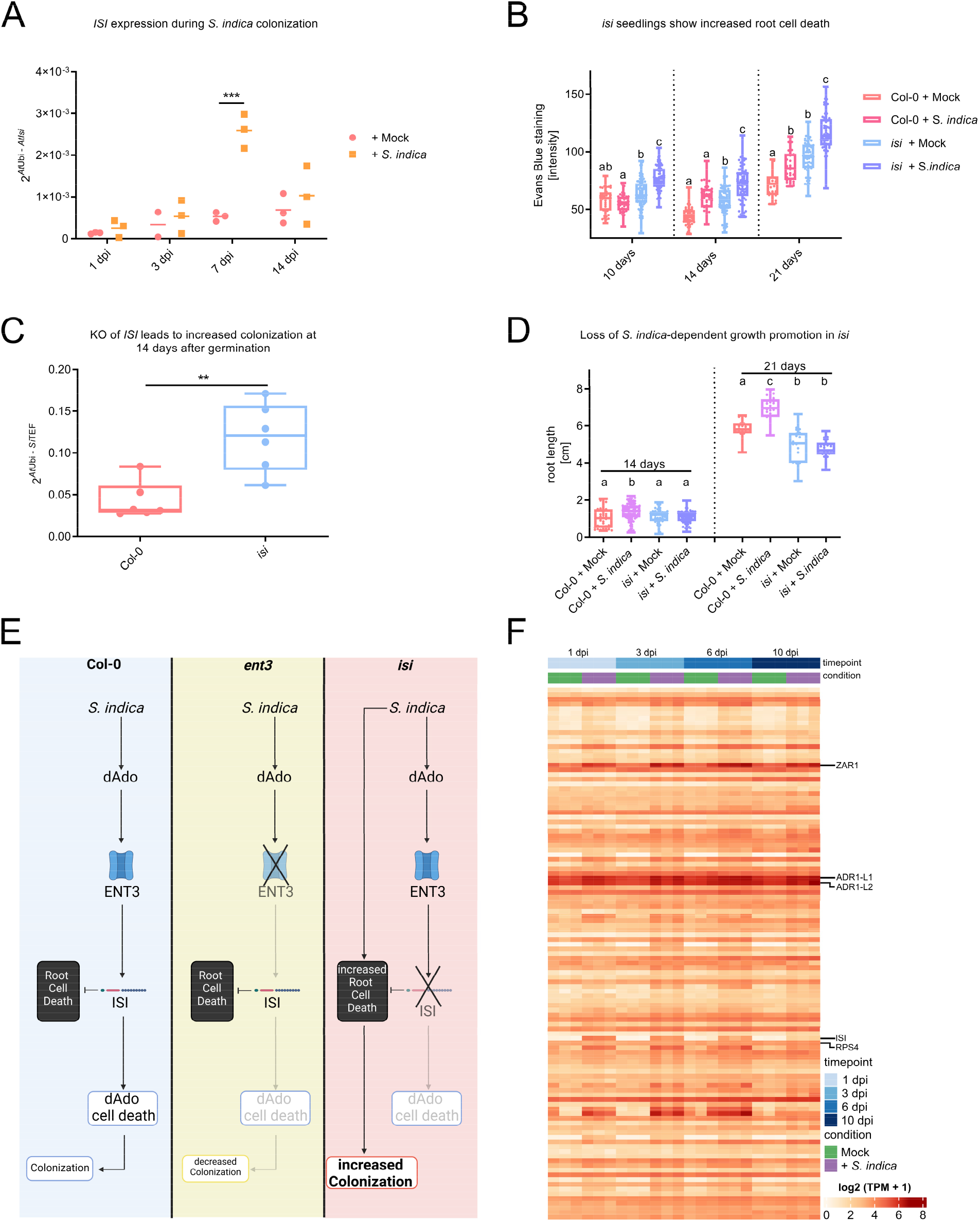
The *At*TIR-NLR AT5G45240 is involved in fungal-mediated growth promotion. (A) AT5G45240 *(AtISI)* expression in *A. thaliana* roots inoculated with *S. indica* or mock treated. Expression was measured via qRT-PCR and calcuated with the 2^−ΔCT^ method. Data points depict biological replicates and asterisks represent significant differences analyzed by Student’s t-test (p < 0.005 ***). (B) Quantification of root cell death in Col-0 WT and *isi* roots colonized by *S. indica* at 10, 14, and 21 days after germination. Cell death was assessed by Evans blue staining. Boxplots show data from 28-80 biological replicates. Different letters indicate significant differences between samples from one time point as assessed by two-way ANOVA with post-hoc Tukey HSD test (p < 0.05). (C) Abundance of *S. indica* in Arabidopsis seedlings 14 days after germination. Fungal (*SiTEF*) to plant (*AtUbi)* ratios were calculated using cDNA as template and method 2^−ΔCT^. Boxplots represent 6 independent biological replicates. Asterisks indicate a significant difference from Col-0 WT samples (Student’s t-test, p < 0.01 **). (D) Root length of Arabidopsis seedlings 14 and 21 days after germination in the presence of *S. indica* or mock treatment. Boxplots show data from 23-86 biological replicates. Different letters indicate significant differences in samples from one time point using two-way ANOVA with post-hoc Tukey HSD test (p < 0.05). The experiment was repeated 2 times independently with similar results. (E) Current model illustrating cell death pathways occurring during colonization of *S. indica* in Arabidopsis Col-0 (blue), *ent3* mutant (yellow), and *isi* mutant (red). (F) The heatmap shows the expression values of *A. thaliana* NLR genes with NB-ARC and LRR (NL) domains in *A. thaliana* root samples as log2 transformed TPM values. Samples were taken at 1, 3, 6 and 10 days post inoculation with *S. indica* or mock treatment. A more detailed version of the heatmap including an in depth description can be found in Figure S25.

## Discussion

### Symbiotic cell death: an evolutionarily conserved mechanism?

Root colonization by *S. indica* is associated with restricted host cell death. The mechanisms behind the induction and regulation of this symbiotic cell death are poorly understood. Previous work has shown that *S. indica* secretes *Si*E5NT, a ubiquitous apoplastic fungal enzyme that catalyzes the conversion of adenosine nucleotides released from immune-activated or damaged host tissues to adenosine (Nizam *et al*., 2019). The mechanism of the secreted ecto-5′-nucleotidase for host colonization appears to be similar to that evolved by *S. aureus* and other animal pathogens to evade the host immune response by secreting AdsA, which converts eATP to Ado, a suppressor of immunity in animal cells (Thammavongsa et al., 2009). In addition to the release of eATP by damaged animal cells, activated neutrophils release NETs, extracellular matrices composed of nuclear and mitochondrial DNA and equipped with granular proteins, cell-specific proteases, and antimicrobial peptides (Brinkmann et al., 2004). NETs rapidly immobilize and kill bacterial pathogens (Driouich *et al*., 2019). *S. aureus* escapes NETs by secreting proteases and the nuclease Nuc, which degrade antimicrobial peptides and DNA. Nuclease-mediated degradation of neutrophil NETs results in the formation of dAMP, which is converted to dAdo by *S. aureus* AdsA. Macrophages and other immune cells are highly sensitive to dAdo intoxication. This cellular intoxication is mediated by uptake via equilibrative nucleoside transporters (Thammavongsa *et al*., 2013; Winstel *et al*., 2018). The escape mechanism of *S. aureus* relies on the interaction of two extracellular microbial enzymes and allows staphylococci to block infiltration of abscess lesions by phagocytes and elimination of the bacteria. Similar to neutrophils, plant roots secrete RETs, extracellular root traps composed of extracellular DNA and a variety of antimicrobial compounds and polysaccharides. RETs and NETs share similar compositional and functional properties, but unlike NETs, RET production is not triggered only upon microbial infection; instead, RETs are continuously released during root development and form a large, mucus-rich network in the rhizosphere (Chambard *et al*., 2021; Driouich *et al*., 2019; Hawes et al., 2016; Tran *et al*., 2016). By cytological analyses we observed that *S. indica* is able to grow within RETs and along the root tip (Figure S26). The ability of *S. indica* to digest DNA *via* secretion of the nuclease NucA, which accumulates in the root apoplast at the onset of cell death (Nizam *et al*., 2019), suggests the possibility of RET digestion by *S. indica* during colonization. Moreover, localization of NucA in the nuclei of plant cells during S. indica colonization shows that this enzyme is able to digest intracellular and extracellular DNA. Here, we show that the combined enzymatic activity of *Si*E5NT and *Si*NucA, present simultaneously in the apoplast, results in the conversion of oligonucleotides with terminal deoxyadenosine obtained by nuclease digestion of DNA to free deoxyadenosine. Most importantly, we demonstrated that uptake of deoxyadenosine via ENT3 is required for host cell death in roots. Mutation of ENT3 results in reduced cell death during colonization by *S. indica*, demonstrating the importance of extracellular nucleoside uptake in regulating fungal induced cell death. The identification of a previously unknown immune-metabolic axis by which cells respond to extracellular purine nucleosides and trigger cell death in plants suggests some conservation or functional convergence between the immune avoidance and escape mechanisms developed by *S. aureus* and other bacterial pathogens in animals and the cell death triggered by dAdo during plant-fungal endophyte interactions in roots.

### A TIR-NLR regulates host cell death in roots

To fight off infections by microbial pathogens, plants have evolved immune receptors that are essential for a successful defense response. PRRs localized in the plasma membrane are capable of sensing conserved MAMPs/DAMPs and eliciting a relatively mild PTI immune response. Successful plant-associated microbes can provide a range of effectors to attenuate PTI to enable successful host colonization. In turn, plants have evolved polymorphic intracellular resistance proteins (R-proteins) to recognize the presence and/or activity of effectors, resulting in a robust defense response called effector-triggered immunity (ETI) that potentiates PTI (Ngou *et al*., 2021; Yuan *et al*., 2021). Most R proteins belong to the nucleotide-binding (NB) leucine-rich repeat (LRR) family (NLR), which are classified on the basis of their N-terminal domains as Toll-like interleukin 1 receptor (TIR) type NLRs (TNLs) or coiled-coil type NLRs (CNLs) (Wu et al., 2019). TNLs generally function as sensors for microbial effectors, whereas several CNLs are referred to as helper NLRs and are downstream of many sensor NLRs in Arabidopsis (Saur et al., 2021; Wu *et al*., 2019). The involvement of the TNL ISI in resistance to dAdo-mediated cell death in Arabidopsis suggests that this TIR-NLR might guard a protein targeted by dAdo. Alternatively, it is possible that dAdo is converted to a plausible substrate of TNLs associated with either nicotinamide adenine dinucleotide (NAD) or 2′,3′-cAMP, a noncanonical cyclic nucleotide monophosphate (cNMP) (Horsefield et al., 2019; Wan et al., 2019; Yu et al., 2021). Plant TIR domains of NLRs are enzymes that can degrade NAD in its oxidized form (NAD^+^). The NADase function of the TIR domain is necessary but not sufficient to activate plant immune responses. Recently, it was shown that plant TIR proteins are not only NADases but also act as 2′,3′-cAMP/cGMP synthetases by hydrolyzing RNA/DNA. Mutations that specifically disrupt synthetase activity abolish TIR-mediated cell death in *N. benthamiana*, demonstrating an important role for these cNMPs in TIR signaling. (Yu *et al*., 2021). The accumulation of extracellular 3′,5′-cAMP upon *S. indica* colonization and treatment with dAdo establishes a link to cyclic nucleotide monophosphates. Further analysis is needed to characterize the metabolism of dAdo in plant cells and to determine how intracellular dAdo leads to TNL activation in Arabidopsis. Since NLRs of the TIR domain type are also found in animals, our results open the possibility to further investigate the role of TNLs in cell death triggered by dAdo in plants and beyond.

In animal systems, it has been shown that following import of dAdo into macrophages, dAdo-mediated toxicity involves conversion of dAdo to dAMP by deoxycytidine kinase (DCK) and adenosine kinase (ADK) activity and signaling via subsequent conversion to the corresponding di- and tri-phosphates by nucleotide kinases and activation of caspase-3-induced apoptosis (Winstel *et al*., 2018). The absence of caspases in plants and the involvement of a TIR-NLR protein in dAdo-mediated cell death in Arabidopsis strongly suggest that this part of the signaling pathway is not conserved between plants and animals and relies on different regulatory and execution mechanisms that require further investigation. The role of the EDS1 family of immunity regulators, which are genetically required for pathogen resistance and execution of cell death by various TIR-NLRs (Lapin et al., 2020), in ISI-mediated cell death also remains to be investigated, as does the involvement of other ISI-like proteins, given that the dAdo-resistance phenotype of ISI is not complete.

In summary, we have uncovered a cellular signaling pathway that responds to extracellularly produced metabolites and links nucleoside transport by an equilibrative nucleoside transporter to cellular activation of the MEP pathway and cell death regulated by the TIR-NLR ISI. The fact that dAdo triggers cell death in several plant species, including a basal lineage, suggests that this pathway is likely conserved and represents an ancient cell death mechanism that is shared for the establishment of plant-endophyte symbiosis. This paves the way for a better understanding of immunometabolism in plant-microbe interactions.

## Material and methods

### Fungal strains and cultivation techniques

*S. indica* strain DSM11827 (German Collection of Microorganisms and Cell Cultures, Braunschweig, Germany) was cultured on complete medium (CM) containing 2% (w/v) glucose and 1.5% (w/v) agar as previously described in (Hilbert et al., 2012). Liquid (CM) cultures were incubated from spores at 28 °C and 120 rpm.

### Plant lines and Arabidopsis transformation

Seeds of *A. thaliana* ecotype Columbia 0 (Col-0), Col-0^AEQ^, and Col-0^AEQ^ *dorn1* (Choi et al. 2014), SALK lines SALK_034517C (*isi*) and SALK_204257C (*ent3*) were used in the experiments. In addition, Col-0 expressing histone H2B fused to mCherry (H2B:2xmCherry) under the UBQ10 promoter in root cells (Marquès-Bueno et al., 2016) were used in this study. Col-0 transformation was performed using the floral dip method as described in (Nizam *et al*., 2019) to generate the transgenic lines *35S::SiNucA* and *35S::SiNucA:mCherry* (with and without signal peptide).

For *Marchantia polymorpha*, the gemmae of the male Tak1 and female Tak2 gametophytes were used.

### Plant growth conditions and fungal inoculation

Surface sterilized seeds were germinated and grown on ½ MS medium (Murashige-Skoog medium, with vitamins, pH 5.7) containing 1% (w/v) sucrose and 0.4% (w/v) Gelrite under short-day conditions (8 h light, 16 h dark) with 130 μmol m^−2^ s^−1^ light and 22 °C/18 °C. Two methods were used for fungal inoculation: 1) 7-day-old seedlings were transferred to ½ MS without sucrose or 1/10 PNM (Plant Nutrion Medium, pH 5.7) plates (15-20 seedlings per plate). 1 ml of water containing 5×10^5^ chlamydospores of *S. indica* was pipetted onto the root and surrounding area. Control plants were inoculated with sterile water. 2) Sterile *A*. thaliana seeds were incubated in 1 ml of water containing 5×10^5^ *S. indica* chlamydospores for one hour and then pipetted onto 1/10 PNM plates.

For harvesting, individual roots were thoroughly washed with water, a 4 cm root section was cut 0.5 cm below the shoot, and immediately frozen in liquid nitrogen. Two to three plates of 20 seedlings each were pooled per replicate.

### Confocal microscopy

Colonized Arabidopsis roots were treated with 5 μg/ml Wheat Germ Agglutinin-Alexa Fluor 488 conjugate (WGA-AF 488, Life Technologies, Thermo Fisher Scientific, Schwerte, Germany) or 1 μM FGB1-FITC of (Wawra et al., 2019) To visualize the fungal cell wall. 250 ng/ml 4′,6-diamidino-2′-phenylindole dihydrochloride (DAPI) or 500 nM SYTOX Orange (Life Technologies, Thermo Fisher Scientific, Schwerte, Germany) were used as nucleic acid stain.

A TCS SP8 confocal microscope (Leica, Wetzlar, Germany) was used for confocal laser scanning microscopy on living cells. AF 488 and FITC were excited with an argon laser at 488 nm, and the emitted light was detected with a hybrid detector at 500-550 nm. mCherry and SYTOX orange were excited with a DPSS laser at 561 nm, and the signal was detected with a hybrid detector at 590-660 nm. DAPI was excited with a diode laser at 405 nm, and the emitted light was detected with a hybrid detector at 415-460 nm.

### DNA and RNA extraction

DNA was extracted from frozen root or fungal material as described in Wawra et al. 2016. Briefly, approximately 500 mg of ground frozen material was dissolved in 1 ml of CTAB extraction buffer (100 mM TrisHCl pH 7.5, 50 mM EDTA pH 8, 1.5 M NaCl, 2% (w/v) cetyltrimethylammonium bromide, 0.05% (v/v) ß-mercaptoethanol) and homogenized for 10 min. 500 µl chloroform:isoamyl alcohol mixture (24:1) was added and the tubes were mixed and centrifuged for 5 min. Ethanol and 1 volume of chloroform:isoamyl alcohol mixture (24:1) were added to the upper phase and centrifuged again. The DNA in the upper phase was precipitated with 1 volume of isopropanol at 4 °C for 1 h.

RNA was extracted with TRIzol (Invitrogen, Thermo Fisher Scientific, Schwerte, Germany) and DNA was digested with DNase I (Thermo Fisher Scientific, Schwerte, Germany) according to the manufacturer’s instructions. cDNA was synthesized using the Fermentas First Strand cDNA Synthesis Kit (Thermo Fisher Scientific, Schwerte, Germany).

### Quantitative RT-PCR analysis

For quantitative real-time PCR, the 2x GoTaq qPCR Master Mix (Promega, Mannheim, Germany) was used. 500 nM forward and reverse primers and 10-20 ng of cDNA or gDNA template were added to each. The reaction was performed in a CFX connect real time system (BioRad, Munich, Germany) with the following program: 95°C 3min, 95°C 15s, 59°C 20s, 72°C 30s, 40 cycles and melting curve analysis. Relative expression was calculated using the 2^−ΔΔCT^ method (Livak and Schmittgen 2001). All oligonucleotides used can be found in Table S8.

### *Serendipita* indica transformation

*S. indica* protoplasts were transformed using the PEG-mediated transformation system as described in (Wawra *et al*., 2019).

### Nuclease activity test

To assay the nuclease activity of the purified protein, 10 nM *Si*NucA was mixed with 100 ng linearized plasmid, 1 µg gDNA, or 1 µg RNA in buffer (5 mM Tris pH 8, 1 mM MgCl_2_, 1 mM CaCl_2_, 0.1% microelement solution (from CM medium)). The mixture was incubated at RT for 1 to 30 min, then loading dye was added and samples were run on a 1-2 % agarose gel.

To test nuclease activity in the filtrate of an *S. indica* culture, a 5-day-old CM culture was minced and incubated for an additional 3 days. The culture was filtered through Miracloth (Millipore Merck, Darmstadt, Germany), and 50 μl of culture filtrate was added to linearized plasmid DNA, gDNA, or RNA.

### *Si*NucA-HA-His purification

A 7-day-old liquid culture of *S. indica* grown in CM medium was filtered through Miracloth. The mycelium was washed with 0.9% NaCl and minced in fresh CM medium in a mixer (MicrotronR MB550 homogenizer (Kinematica, Lucerne, Switzerland)). The culture was regenerated for two days. Subsequently, the culture was filtered with Miracloth and through a 0.45 μm membrane filter. To this cell-free culture filtrate, 1 mM phenylmethanesulfonyl fluoride (PMSF) was added and the pH was adjusted to pH 7 with 1 M Tris pH 8. Proteins were precipitated with 80% ammonium sulfate. The protein pellet was resuspended in 20 mM Tris pH 8. Proteins were separated by size exclusion chromatography (Sephadex G 200 column, Hiload 6/600) using a 20 mM Tris pH 8 / 150 mM NaCl buffer. Fractions containing *Si*NucA-HA-His were desalted by dialysis and checked by SDS-PAGE and anti-HA Western blot. The protein was stored in 20 mM Tris pH 8. The identity of the protein was verified by LC-MS/MS.

### Purification of proteins and measurement of enzyme kinetics of *Si*E5NT

*Si*E5NT fused to a C-terminal hemaglutinin (HA) and Strep tag (pXCScpmv-HAStrep, V69) was transiently expressed in *Nicotiana benthaminana* and purified by affinity chromatography as described by (Myrach et al., 2017; Werner et al., 2008). Enzyme concentrations were determined using bovine serum albumin standards after SDS-PAGE and Coomassie Blue staining with an Odyssey Fc Dual Mode Imaging System (Li-cor Biosciences, Germany).

To determine substrate specificity, different substrates were tested. Briefly, 10 µl of purified enzyme (1.9 × 10^−4^ mg protein) was added to 70 µl of reaction buffer (10 mM MES buffer, pH 6.0; 1 mM CaCl_2_ and 1 mM MgCl_2_) and pre-incubated for 5 minutes. Substrates were added (20 µl of a 5 mM stock solution) to give a final reaction volume of 100 µl. The reaction was quenched with 400 µl of MeOH at the indicated times and centrifuged for 10 min at 21,000g at 4°C. The supernatant was evaporated in a vacuum concentrator and the dry pellet resuspended in 100 µl of HPLC mobile phase A.

Samples were analyzed using an Agilent 1200 SL HPLC system equipped with a diode array detector. 10 µl of the sample was injected onto a Supelcosil LC-18-T column (Sigma-Aldrich) at a flow rate of 0.8 ml min^−1^ and a column temperature of 25°C. The analytes were separated with the following gradient: 0 min, 100% A; 9 min, 100% A; 15 min, 75% A; 17.5 min, 10% A; 19 min, 0% A; 23 min, 0% A; 24 min, 100% A; 30 min, 100% A. Mobile phase A consisted of 100 mM KH_2_ PO_4_, pH 6.0, in deionized water and mobile phase B consisted of 90% 100 mM KH_2_ PO_4_, pH 6.0, in deionized water and 10% MeOH. Appropriate standard solutions were used for quantification. Data were analyzed using Agilent Chemstation software.

Kinetic constants for deoxyadenosine monophosphate and adenosine monophosphate were determined using the EnzCheck (Thermo Fisher Scientific, Waltham, USA) phosphatase assay kit according to specifications, using only one-tenth of the recommended reaction volume.

### Release of deoxynucleosides from salmon sperm DNA by *Si*NucA and *Si*E5NT

10 µg of salmon sperm DNA was incubated with 4 µl of *Si*NucA, 10 µl of *Si*E5NT, or a combination of both in a final volume of 40 µl containing 10 mM HEPES, pH 7.2; 100 mM MgCl_2_ ; and 1 mM DTT. The enzymes were incubated in the buffer mixture for 5 minutes prior to the addition of DNA. The reaction was incubated at 22 °C for 1 hour and inactivated at 95 °C for 5 minutes. Subsequently, centrifugation was performed at 40000 g and 4 °C for 20 minutes. 20 µl of the supernatant was analyzed on a 2% agarose gel. The remaining 20 µl of the supernatant was mixed with 50 µl of water and used for HPLC MS/MS analysis. Samples and standards were analyzed using an Agilent 1290 Infinity II HPLC system coupled to an Agilent 6470 triple quadrupole mass spectrometer. Analytes (10 µl samples) were separated on a 50 × 4.6 mm Polaris C18A column (Agilent) using the following gradient: 0 min, 96% A; 8 min, 35% A; 8.2 min, 0% A; 10 min, 0% A; 10.1 min, 96% A. Mobile phase A consisted of 10 mM ammonium acetate, pH 7.5, and mobile phase B was pure MeOH. The flow rate was 0.6 ml min^−1^ and the column temperature was 30°C. In-source parameters were set as previously described for deoxyribonucleoside analysis (Straube et al., 2021). All analytes were measured in positive mode. Transitions (precursor and production), collision energies, and fragmentor energies are listed in Table S9.

### PAM fluorometric measurements

PAM fluorometry measurements were performed by transferring 9-day-old Arabidopsis seedlings into 24 well plates containing 2 ml of 2.5 mM MES buffer (pH 5.6). 3 seedlings were pooled in one well. After 24 hours of regeneration, seedlings were treated with solutions of 5’ deoxyadenosine (dAdo), adenosine (Ado), or methyl jasmonate (all Sigma-Aldrich, Taufkirchen, Germany), adjusting to a final concentration of 500 µM. Treated seedlings were incubated in complete darkness for 20 min to reach a dark-adapted condition. The photosynthetic activity of the plants was measured using the M-Series PAM fluorometer (Heinz Walz GmbH, Effeltrich, Germany). Data were analyzed using ImagingWin software (v.2.41a; Walz, Germany). When further analysis of photosynthetic area development was required, it was evaluated using Fiji (ImageJ).

### Cell death staining with Evans blue

A modified protocol as described in (Vijayaraghavareddy et al., 2017)was used. To quantify cell death induced by *S. indica* in *Arabidopsis*, plants were used for cell death staining at three time points, 7, 10, and 14 days after fungal spore inoculation, using five plants per treatment. For cell death induced by dAdo (or chemically induced cell death), plants were microscoped after 4 days of cell death treatment. To remove external fungal growth or chemical treatment solutions, plants were washed three times in ddH_2_ O before cell death staining in a 2.5 mM Evans blue solution (Sigma-Aldrich, lot #MKCH7958) dissolved in 0.1 M CaCl_2_ pH 5.6 for 15 minutes. After extensive washing for one hour with ddH_2_ O, images were captured using a Leica M165 FC microscope.

### Activity-based protein profiling

After treatment with 500 µM dAdo, 500 µM Ado, or MES buffer, 9-day-old Arabidopsis seedlings were frozen in liquid nitrogen, ground, and dissolved in 50 mM Tris-HCl buffer (pH 7). After centrifugation, the supernatant was divided and treated with either the proteasome inhibitor MG132 (final concentration: 50 µM, company) or DMSO for 30 min. Samples were then incubated with 26S proteasome probe MVB072 (final concentration: 1 µM, company): 1 µM, (Kolodziejek et al., 2010) Samples were then denatured in SDS loading dye at 95 °C and separated on 12% SDS gels. The probe was visualized using the rhodamine settings (excitation: 532 nm, emission: 580 nm) on a ChemiDoc (BioRad, CA, USA). The protein content of the samples was visualized by staining the gel with SYPRO^™^ Ruby (Invitrogen, Carlsbad, CA, USA) according to the manufacturer’s instructions.

### RNA seq

7-day-old seedlings were transferred from plates to individual wells of 24-well plates containing 0.5 ml of liquid ½ MS medium containing 0.5% (w/v) sucrose. After 5 additional days in liquid culture, the medium was replaced with 1 ml of 500*μ* M dAdo in 2.5 mM MES buffer pH 5.6 or 2.5 mM MES buffer pH 5.6 alone as a negative control. Three replicates of four seedlings were harvested for each treatment at 0, 3, and 12 h after treatment (hpt), frozen in liquid nitrogen, and stored at -80° C until processing for RNA extraction. Total RNA was extracted using TRIZOL reagent as described above. RNA integrity was confirmed by gel electrophoresis, and quantity and purity were determined using a NanoDrop 2000. Stranded mRNA-seq libraries were prepared according to the manufacturer’s instructions (Vazyme Biotech Co., Nanjing, China). Qualified libraries were sequenced on a HiSeq 3000 system instrument in the Genomics & Transcriptomics Laboratory at Heinrich Heine University to generate >100 million reads with a read length of 150 bp from three biological replicates. Trimmomatic v. 0.36,(Bolger et al., 2014) was used for quality trimming and adapter clipping. Reads were then mapped to Arabidopsis TAIR10 CDS assembly and quantified using kallisto v. 0.46.2,(Bray et al., 2016), resulting in estimated counts and transcripts per million (TPM) values. The log2 fold difference in gene expression between conditions was estimated using the R packages tximport (Soneson et al., 2015) and DESeq2 (Love et al., 2014). Genes with statistical significance were selected (FDR-adjusted p-value < 0.05). Data have been deposited at NCBI under GEO accession number GSE209761.

### Collection of extracellular fluid

7-day-old Arabidopsis seedlings germinated on ½ MS agar containing 1% sucrose were transferred to 24 well plates. Each well contained 3 seedlings in 1.5 ml of 2.5 mM MES and the specific treatment. After transfer, the plants were returned to the growth chamber for three days. The seedlings were then removed and the liquid centrifuged at 4000 g for 15 minutes. The supernatant was freeze dried and sent for metabolite analysis.

### Metabolite analysis

5 µl of the apoplastic liquid was injected into an Acquity UPLC (Waters Inc.) equipped with a Nucleoshell RP18 column (Macherey & Nagel, 150mm × 2 mm × 2.1µm) using tributylammonium as the ion pairing agent. Solvent A: 10 mM tributylamine (aqueous) acidified with glacial acetic acid to pH 6.2; solvent B acetonitrile. Gradient: 0-2 min: 2% B, 2-18 min 2-36% B, 18-21 min 36-95% B, 21-22.5 min 95% B, 22.51-24 min 2% B. Column flow was 0.4 ml min^−1^ throughout. The column temperature was 40 °C. Scheduled metabolite detection based on multiple reaction monitoring (MRM) was performed in negative mode with electrospray ionization (ESI) on a QTrap 6500 (AB-Sciex GmbH, Darmstadt, Germany): Ion source gas 1: 60 psi, ion source gas 2: 70 psi, curtain gas: 35 psi, temperature: 450 °C, ion spray voltage floating: and -4500V). MRM transitions of 189 metabolites covering central carbon and energy metabolism were previously signal optimized and retention times determined (Table S10).

### Ca^2+^ influx quantification

Calcium influx assays were performed as previously described in (Wanke et al. 2020). Briefly, individual 7-day-old Arabidopsis seedlings were placed in white 96-well plates filled with 200 µl reconstitution buffer (2.5 mM MES pH 5.7 [Sigma-Aldrich, Taufkirchen, Germany], 10 mM CaCl_2_ [Roth, Karlsruhe, Germany]). Before incubation overnight in the dark, the solution was replaced with 133 µl reconstitution buffer containing 10 mM coelenterazine (Roth, Karlsruhe, Germany). The following day, chemiluminescence was measured using a TECAN SPARK 10M microplate reader. After baseline measurement, 67 µl of three-fold concentrated elicitor solutions (or Milli-Q water as a sham control) were added manually. Cytosolic calcium influx after addition of the trigger was measured continuously for 30 min. To determine the undischarged aequorin for treatment normalization, 100 μl 3 M CaCl_2_ (in 30% EtOH) was injected into each well, followed by constant measurement for 1 minute. All steps were performed with an integration time of 450 msec.

### Heterologous protein production in *Nicotiana benthamiana* and protein purification

For heterologous protein production in Nicotiana *benthamiana*, leaves of 4-week-old plants *were* infiltrated with *Agrobacterium tumefaciens* GV3101 strains. *A. tumefaciens* was grown in LB liquid medium with the appropriate antibiotics at 28°C and 180 rpm for 2 days until an OD_600_ of 1 was achieved. Cultures were centrifuged at 3500 rpm for 15 min at RT, the supernatant was discarded and resuspended in 1 ml infiltration buffer (10 mM MES pH 5.5, 10 mM MgCl_2_, 200 µM acetosyringone) and incubated for 1 h in the dark at 28°C, 180 rpm. All strains were diluted with the infiltration buffer to an OD_600_ of 1. Each strain was mixed with P19-expressing strains at a 1:1 ratio. 2-3 leaves per plant were infiltrated with a needleless syringe. After 4 dpi, leaves were separated from the plant and crushed in liquid nitrogen. Protein purification was performed according to (Werner *et al*., 2008) with minor modifications performed: A 15-ml tube was filled to the 2-ml mark with ground leaf material, and 2 ml of cold extraction buffer (100 mM Tris pH 8.0, 100 mM NaCl, 5 mM EDTA, 0.5% Triton X100, 10 mM DTT, 100 μg/ml avidin) was added. The powder was resuspended by vortexing and then centrifuged at 12000 rpm and 4°C for 10 minutes. The supernatant was transferred to a new tube and 100 µl of Strep-Tactin® Macroprep (50% slurry) was added and incubated for 60 min at 4°C in a rotating wheel. The mixture was centrifuged at 700xg at RT for 30 s and the supernatant was completely removed from the beads. The beads were washed once with 4 ml and two additional times with 2 ml of wash buffer (50 mM Tris pH 8.0, 100 mM NaCl, 0.5 mM EDTA, 0.005% Triton X-100, 2 mM DTT) by centrifugation for 30 s at 700xg and the supernatant was discarded. During the final wash, beads were transferred to a 1.5 ml tube with a low binding level. The beads were either boiled directly for 5 min at 95°C with 6x SDS loading dye to run on an SDS-PAGE, or the proteins were eluted from the beads by adding 100 µl of elution buffer (wash buffer + 2.5 - 10 mM biotin) and incubating at 25°C, >800 rpm for 5 min. Samples were centrifuged at 700xg for 20 s and elution was repeated. Elution fractions were pooled, SDS loading dye was added, and samples were run on SDS-PAGE followed by Western blot.

### Ion leakage measurements

The seedlings for measurement of ion leakage were prepared in the same manner as seedlings for PAM fluorometry measurements. Ion leakage was measured with a conductivity meter (LAQUAtwin EC-11; Horiba, Newhampton, UK).

### Seed germination test

Sterile Arabidopsis seedlings were transferred into 24-well plates containing 2 ml of 1/10 PNM medium. The medium contained either 500 µM dAdo or the same volume of 2.5 mM MES (pH 5.6). Ten seeds were placed in each well and grown under short-day conditions after 2 days of stratification. Seedling growth was monitored by PAM fluorometry.

### Root length measurements

To evaluate root length, scans of the square plates containing seedlings were analyzed using Fiji (ImageJ) and the length of the primary root was measured.

### Oxidative burst assay

Individual 7-day-old Arabidopsis seedlings were transferred to white 96-well plates containing 200 µl reconstitution buffer (2.5 mM MES pH 5.7 (Sigma-Aldrich, Taufkirchen, Germany), 10 mM CaCl_2_ (Roth, Karlsruhe, Germany) and incubated overnight in the growth chamber. The following day, the buffer was replaced with 133 µl reconstitution solution containing 15 µgml^−1^ horseradish peroxidase (Sigma-Aldrich, Taufkirchen, Germany) and 15 µM L-O12 (Wako Chemicals, Neuss, Germany). After 10 min incubation, 67 µL of three-fold concentrated elicitor solutions (or Milli-Q water as a mock control) were added manually to the wells. Measurements were started immediately and chemiluminescence was measured continuously with a TECAN SPARK 10M microplate reader (Tecan, Männedorf, Switzerland) at an integration time of 450 ms.

## Acknowledgements

We would like to thank Prof. Jijie Chai and Jane Parker for discussions and reading the manuscript prior to submission. We further want to thank Johana Misas Stadtel for her support during ABPP assays. ND & AZ would like to thank Lisa Madhi, Lisa Leson and Lucia Schmitz for their excellent support in the lab. ND was supported by the International Max Planck Research School (IMPRS) on “Understanding Complex Plant Traits using Computational and Evolutionary Approaches”, by the University of Cologne and by the SFB 1403 Project ID: 1403-414786233. AZ gratefully acknowledges support from the Cluster of Excellence on Plant Sciences (CEPLAS) funded by the German Research Foundation (DFG) under the Excellence Strategy - EXC 2048/1 - Project ID: 390686111 and SFB 1403 Project ID: 1403-414786233. We further would like to thank the U.S. Department of Energy Joint Genome Institute (https://ror.org/04xm1d337) and Yu Zhang, Sravanthi Tejomurthula, Daniel Peterson, Vivian Ng & Igor Grigoriev for producing sequencing data within the work proposal 10.46936/10.25585/60001292.

CPW gratefully acknowledges support from DFG grant WI3411/8-1. Some of the graphical representations were generated using the online tool BioRender.

## References

Antonioli, L., Pacher, P., Vizi, E.S., and Hasko, G. (2013). CD39 and CD73 in immunity and inflammation. Trends in Molecular Medicine 19, 355–367. 10.1016/j.molmed.2013.03.005.

Bolger, A.M., Lohse, M., and Usadel, B. (2014). Trimmomatic: a flexible trimmer for Illumina sequence data. Bioinformatics 30, 2114–2120. 10.1093/bioinformatics/btu170.

Bray, N.L., Pimentel, H., Melsted, P., and Pachter, L. (2016). Near-optimal probabilistic RNA-seq quantification. Nature biotechnology 34, 525–527. 10.1038/nbt.3519.

Brinkmann, V., Reichard, U., Goosmann, C., Fauler, B., Uhlemann, Y., Weiss, D.S., Weinrauch, Y., and Zychlinsky, A. (2004). Neutrophil extracellular traps kill bacteria. Science 303, 1532–1535. 10.1126/science.1092385.

Chambard, M., Plasson, C., Derambure, C., Coutant, S., Tournier, I., Lefranc, B., Leprince, J.M., Kiefer-Meyer, M.C., Driouich, A., Follet-Gueye, M.L., and Boulogne, I. (2021). New Insights into Plant Extracellular DNA. A Study in Soybean Root Extracellular Trap. Cells 10. ARTN 69 10.3390/cells10010069.

Choi, J., Tanaka, K., Cao, Y., Qi, Y., Qiu, J., Liang, Y., Lee, S.Y., and Stacey, G. (2014). Identification of a plant receptor for extracellular ATP. Science 343, 290–294. 10.1126/science.343.6168.290.

D’Haeze, W., De Rycke, R., Mathis, R., Goormachtig, S., Pagnotta, S., Verplancke, C., Capoen, W., and Holsters, M. (2003). Reactive oxygen species and ethylene play a positive role in lateral root base nodulation of a semiaquatic legume. Proceedings of the National Academy of Sciences 100, 11789–11794. 10.1073/pnas.1333899100.

Deshmukh, S., Huckelhoven, R., Schafer, P., Imani, J., Sharma, M., Weiss, M., Waller, F., and Kogel, K.H. (2006). The root endophytic fungus Piriformospora indica requires host cell death for proliferation during mutualistic symbiosis with barley. Proc Natl Acad Sci U S A 103, 18450-18457. 0605697103 [pii] 10.1073/pnas.0605697103.

Driouich, A., Smith, C., Ropitaux, M., Chambard, M., Boulogne, I., Bernard, S., Follet-Gueye, M.L., Vicre, M., and Moore, J. (2019). Root extracellular traps versus neutrophil extracellular traps in host defence, a case of functional convergence? Biol Rev Camb Philos Soc 94, 1685–1700. 10.1111/brv.12522.

Hawes, M., Allen, C., Turgeon, B.G., Curlango-Rivera, G., Tran, T.M., Huskey, D.A., and Xiong, Z.G. (2016). Root Border Cells and Their Role in Plant Defense. Annu Rev Phytopathol 54, 143–161. 10.1146/annurev-phyto-080615-100140.

Hilbert, M., Voll, L.M., Ding, Y., Hofmann, J., Sharma, M., and Zuccaro, A. (2012). Indole derivative production by the root endophyte Piriformospora indica is not required for growth promotion but for biotrophic colonization of barley roots. New Phytologist 196, 520–534. 10.1111/j.1469-8137.2012.04275.x.

Horsefield, S., Burdett, H., Zhang, X., Manik, M.K., Shi, Y., Chen, J., Qi, T., Gilley, J., Lai, J.S., Rank, M.X., et al. (2019). NAD(+) cleavage activity by animal and plant TIR domains in cell death pathways. Science 365, 793–799. 10.1126/science.aax1911.

Kilian, J., Whitehead, D., Horak, J., Wanke, D., Weinl, S., Batistic, O., D’Angelo, C., Bornberg-Bauer, E., Kudla, J., and Harter, K. (2007). The AtGenExpress global stress expression data set: protocols, evaluation and model data analysis of UV-B light, drought and cold stress responses. The Plant Journal 50, 347–363. https://doi.org/10.1111/j.1365-313X.2007.03052.x.

Kloppholz, S., Kuhn, H., and Requena, N. (2011). A Secreted Fungal Effector of Glomus intraradices Promotes Symbiotic Biotrophy. Curr Biol 21, 1204-1209. S0960-9822(11)00717-2 [pii] 10.1016/j.cub.2011.06.044.

Kolodziejek, I., Misas-Villamil, J.C., Kaschani, F., Clerc, J., Gu, C., Krahn, D., Niessen, S., Verdoes, M., Willems, L.I., Overkleeft, H.S., et al. (2010). Proteasome Activity Imaging and Profiling Characterizes Bacterial Effector Syringolin A Plant Physiology 155, 477–489. 10.1104/pp.110.163733.

Lahrmann, U., Ding, Y., Banhara, A., Rath, M., Hajirezaei, M.R., Doehlemann, S., von Wiren, N., Parniske, M., and Zuccaro, A. (2013). Host-related metabolic cues affect colonization strategies of a root endophyte. Proceedings of the National Academy of Sciences of the United States of America 110, 13965–13970. 10.1073/pnas.1301653110.

Lahrmann, U., Strehmel, N., Langen, G., Frerigmann, H., Leson, L., Ding, Y., Scheel, D., Herklotz, S., Hilbert, M., and Zuccaro, A. (2015). Mutualistic root endophytism is not associated with the reduction of saprotrophic traits and requires a noncompromised plant innate immunity. New Phytologist 207, 841–857. 10.1111/nph.13411.

Lahrmann, U., and Zuccaro, A. (2012). Opprimo ergo sum-Evasion and Suppression in the Root Endophytic Fungus Piriformospora indica. Molecular Plant-Microbe Interactions 25, 727–737. 10.1094/mpmi-11-11-0291.

Lapin, D., Bhandari, D.D., and Parker, J.E. (2020). Origins and Immunity Networking Functions of EDS1 Family Proteins. Annu Rev Phytopathol 58, 253–276. 10.1146/annurev-phyto-010820-012840.

Li, G., Liu, K., Baldwin, S.A., and Wang, D. (2003). Equilibrative nucleoside transporters of Arabidopsis thaliana. cDNA cloning, expression pattern, and analysis of transport activities. The Journal of biological chemistry 278, 35732–35742. 10.1074/jbc.M304768200.

Lo Presti, L., Lanver, D., Schweizer, G., Tanaka, S., Liang, L., Tollot, M., Zuccaro, A., Reissmann, S., Kahmann, R., and Merchant, S. (2015). Fungal Effectors and Plant Susceptibility. In Annual Review of Plant Biology, Vol 66, pp. 513–545. 10.1146/annurev-arplant-043014-114623.

Love, M.I., Huber, W., and Anders, S. (2014). Moderated estimation of fold change and dispersion for RNA-seq data with DESeq2. Genome biology 15, 550. 10.1186/s13059-014-0550-8.

Marquès-Bueno, M.M., Morao, A.K., Cayrel, A., Platre, M.P., Barberon, M., Caillieux, E., Colot, V., Jaillais, Y., Roudier, F., and Vert, G. (2016). A versatile Multisite Gateway-compatible promoter and transgenic line collection for cell type-specific functional genomics in Arabidopsis. The Plant Journal 85, 320–333. https://doi.org/10.1111/tpj.13099.

Mucha, J., Guzicka, M., Ratajczak, E., and Zadworny, M. (2014). Strategies utilized by trophically diverse fungal species for Pinus sylvestris root colonization. Tree Physiol 34, 73–86. 10.1093/treephys/tpt111.

Myrach, T., Zhu, A., and Witte, C.P. (2017). The assembly of the plant urease activation complex and the essential role of the urease accessory protein G (UreG) in delivery of nickel to urease. The Journal of biological chemistry 292, 14556–14565. 10.1074/jbc.M117.780403.

Ngou, B.P.M., Ahn, H.K., Ding, P., and Jones, J.D.G. (2021). Mutual potentiation of plant immunity by cell-surface and intracellular receptors. Nature 592, 110–115. 10.1038/s41586-021-03315-7.

Nizam, S., Qiang, X., Wawra, S., Nostadt, R., Getzke, F., Schwanke, F., Dreyer, I., Langen, G., and Zuccaro, A. (2019). Serendipita indica E5’NT modulates extracellular nucleotide levels in the plant apoplast and affects fungal colonization. EMBO Rep 20. 10.15252/embr.201847430.

Nostadt, R., Hilbert, M., Nizam, S., Rovenich, H., Wawra, S., Martin, J., Kupper, H., Mijovilovich, A., Ursinus, A., Langen, G., et al. (2020). A secreted fungal histidine-and alanine-rich protein regulates metal ion homeostasis and oxidative stress. New Phytol. 10.1111/nph.16606.

Oberwinkler, F., Riess, K., Bauer, R., Selosse, M.-A., Weiß, M., Garnica, S., and Zuccaro, A. (2013). Enigmatic Sebacinales. Mycological Progress 12, 1–27. 10.1007/s11557-012-0880-4.

Olvera-Carrillo, Y., Van Bel, M., Van Hautegem, T., Fendrych, M., Huysmans, M., Simaskova, M., van Durme, M., Buscaill, P., Rivas, S., Coll, N.S., et al. (2015). A Conserved Core of Programmed Cell Death Indicator Genes Discriminates Developmentally and Environmentally Induced Programmed Cell Death in Plants. Plant Physiol 169, 2684–2699. 10.1104/pp.15.00769.

Pham, A.Q., Cho, S.-H., Nguyen, C.T., and Stacey, G. (2020). Arabidopsis Lectin Receptor Kinase P2K2 Is a Second Plant Receptor for Extracellular ATP and Contributes to Innate Immunity1 [OPEN]. Plant Physiology 183, 1364–1375. 10.1104/pp.19.01265.

Plett, J.M., Daguerre, Y., Wittulsky, S., Vayssieres, A., Deveau, A., Melton, S.J., Kohler, A., Morrell-Falvey, J.L., Brun, A., Veneault-Fourrey, C., and Martin, F. (2014). Effector MiSSP7 of the mutualistic fungus Laccaria bicolor stabilizes the Populus JAZ6 protein and represses jasmonic acid (JA) responsive genes. Proc Natl Acad Sci U S A 111, 8299–8304. 10.1073/pnas.1322671111.

Qiang, X., Zechmann, B., Reitz, M.U., Kogel, K.H., and Schafer, P. (2012). The mutualistic fungus Piriformospora indica colonizes Arabidopsis roots by inducing an endoplasmic reticulum stress-triggered caspase-dependent cell death. Plant Cell 24, 794–809. 10.1105/tpc.111.093260.

Rafiqi, M., Jelonek, L., Akum, N.F., Zhang, F., and Kogel, K.H. (2013). Effector candidates in the secretome of Piriformospora indica, a ubiquitous plant-associated fungus. Front Plant Sci 4, 228. 10.3389/fpls.2013.00228.

Ragnelli, A.M., Aimola, P., Maione, M., Zarivi, O., Leonardi, M., and Pacioni, G. (2014). The cell death phenomenon during Tuber ectomycorrhiza morphogenesis. Plant Biosystems - An International Journal Dealing with all Aspects of Plant Biology 148, 473–482. 10.1080/11263504.2013.788575.

Rich-Griffin, C., Stechemesser, A., Finch, J., Lucas, E., Ott, S., and Schäfer, P. (2020). Single-Cell Transcriptomics: A High-Resolution Avenue for Plant Functional Genomics. Trends in plant science 25, 186–197. 10.1016/j.tplants.2019.10.008.

Saur, I.M.L., Panstruga, R., and Schulze-Lefert, P. (2021). NOD-like receptor-mediated plant immunity: from structure to cell death. Nat Rev Immunol 21, 305–318. 10.1038/s41577-020-00473-z.

Schneider, H.M., and Lynch, J.P. (2018). Functional implications of root cortical senescence for soil resource capture. Plant and Soil 423, 13–26. 10.1007/s11104-017-3533-1.

Soneson, C., Love, M.I., and Robinson, M.D. (2015). Differential analyses for RNA-seq: transcript-level estimates improve gene-level inferences. F1000Research 4, 1521. 10.12688/f1000research.7563.2.

Straube, H., Niehaus, M., Zwittian, S., Witte, C.P., and Herde, M. (2021). Enhanced nucleotide analysis enables the quantification of deoxynucleotides in plants and algae revealing connections between nucleoside and deoxynucleoside metabolism. Plant Cell 33, 270–289. 10.1093/plcell/koaa028.

Tedersoo, L., Bahram, M., Ryberg, M., Otsing, E., Koljalg, U., and Abarenkov, K. (2014). Global biogeography of the ectomycorrhizal /sebacina lineage (Fungi, Sebacinales) as revealed from comparative phylogenetics analyses. Mol Ecol. 10.1111/mec.12849.

Thammavongsa, V., Kern, J.W., Missiakas, D.M., and Schneewind, O. (2009). Staphylococcus aureus synthesizes adenosine to escape host immune responses. J Exp Med 206, 2417–2427. 10.1084/jem.20090097.

Thammavongsa, V., Missiakas, D.M., and Schneewind, O. (2013). Staphylococcus aureus degrades neutrophil extracellular traps to promote immune cell death. Science 342, 863–866. 10.1126/science.1242255.

Thürich, J., Meichsner, D., Furch, A.C.U., Pfalz, J., Krüger, T., Kniemeyer, O., Brakhage, A., and Oelmüller, R. (2018). Arabidopsis thaliana responds to colonisation of Piriformospora indica by secretion of symbiosis-specific proteins. PLOS ONE 13, e0209658. 10.1371/journal.pone.0209658.

Tran, T.M., MacIntyre, A., Hawes, M., and Allen, C. (2016). Escaping Underground Nets: Extracellular DNases Degrade Plant Extracellular Traps and Contribute to Virulence of the Plant Pathogenic Bacterium Ralstonia solanacearum. PLoS Pathog 12, e1005686. 10.1371/journal.ppat.1005686.

Vijayaraghavareddy, P., Adhinarayanreddy, V., Vemanna, R.S., Sreeman, S., and Makarla, U. (2017). Quantification of Membrane Damage/Cell Death Using Evan’s Blue Staining Technique. Bio-protocol 7, e2519. 10.21769/BioProtoc.2519.

Voß, S., Betz, R., Heidt, S., Corradi, N., and Requena, N. (2018). RiCRN1, a Crinkler Effector From the Arbuscular Mycorrhizal Fungus Rhizophagus irregularis, Functions in Arbuscule Development. Frontiers in Microbiology 9. 10.3389/fmicb.2018.02068.

Wan, L., Essuman, K., Anderson, R.G., Sasaki, Y., Monteiro, F., Chung, E.H., Osborne Nishimura, E., DiAntonio, A., Milbrandt, J., Dangl, J.L., and Nishimura, M.T. (2019). TIR domains of plant immune receptors are NAD(+)-cleaving enzymes that promote cell death. Science 365, 799–803. 10.1126/science.aax1771.

Wawra, S., Fesel, P., Widmer, H., Neumann, U., Lahrmann, U., Becker, S., Hehemann, J.H., Langen, G., and Zuccaro, A. (2019). FGB1 and WSC3 are in planta-induced beta-glucan-binding fungal lectins with different functions. New Phytol. 10.1111/nph.15711.

Wawra, S., Fesel, P., Widmer, H., Timm, M., Seibel, J., Leson, L., Kesseler, L., Nostadt, R., Hilbert, M., Langen, G., and Zuccaro, A. (2016). The fungal-specific beta-glucan-binding lectin FGB1 alters cell-wall composition and suppresses glucan-triggered immunity in plants. Nat Commun 7, 13188. 10.1038/ncomms13188.

Weiss, M., Waller, F., Zuccaro, A., and Selosse, M.A. (2016). Sebacinales - one thousand and one interactions with land plants. New Phytologist 211, 20–40. 10.1111/nph.13977.

Wen, F.S., White, G.J., VanEtten, H.D., Xiong, Z.G., and Hawes, M.C. (2009). Extracellular DNA Is Required for Root Tip Resistance to Fungal Infection. Plant Physiology 151, 820–829. 10.1104/pp.109.142067.

Werner, A.K., Sparkes, I.A., Romeis, T., and Witte, C.P. (2008). Identification, biochemical characterization, and subcellular localization of allantoate amidohydrolases from Arabidopsis and soybean. Plant Physiol 146, 418–430. 10.1104/pp.107.110809.

Winstel, V., Missiakas, D., and Schneewind, O. (2018). Staphylococcus aureus targets the purine salvage pathway to kill phagocytes. Proc Natl Acad Sci U S A 115, 6846–6851. 10.1073/pnas.1805622115.

Wu, Z., Li, M., Dong, O.X., Xia, S., Liang, W., Bao, Y., Wasteneys, G., and Li, X. (2019). Differential regulation of TNL-mediated immune signaling by redundant helper CNLs. New Phytol 222, 938–953. 10.1111/nph.15665.

Xiao, Y., Savchenko, T., Baidoo, E.E., Chehab, W.E., Hayden, D.M., Tolstikov, V., Corwin, J.A., Kliebenstein, D.J., Keasling, J.D., and Dehesh, K. (2012). Retrograde signaling by the plastidial metabolite MEcPP regulates expression of nuclear stress-response genes. Cell 149, 1525–1535. 10.1016/j.cell.2012.04.038.

Yu, D., Song, W., Tan, E.Y.J., Liu, L., Cao, Y., Jirschitzka, J., Li, E., Logemann, E., Xu, C., Huang, S., et al. (2021). TIR domains of plant immune receptors are 2′,3′-cAMP/cGMP synthetases mediating cell death. bioRxiv, 2021.2011.2009.467869. 10.1101/2021.11.09.467869.

Yuan, M., Jiang, Z., Bi, G., Nomura, K., Liu, M., Wang, Y., Cai, B., Zhou, J.M., He, S.Y., and Xin, X.F. (2021). Pattern-recognition receptors are required for NLR-mediated plant immunity. Nature 592, 105–109. 10.1038/s41586-021-03316-6.

Zipfel, C., and Oldroyd, G.E.D. (2017). Plant signalling in symbiosis and immunity. Nature 543, 328–336. 10.1038/nature22009.

Zuccaro, A., Lahrmann, U., Gueldener, U., Langen, G., Pfiffi, S., Biedenkopf, D., Wong, P., Samans, B., Grimm, C., Basiewicz, M., et al. (2011). Endophytic Life Strategies Decoded by Genome and Transcriptome Analyses of the Mutualistic Root Symbiont Piriformospora indica. Plos Pathogens 7. 10.1371/journal.ppat.1002290.

