## Supplementary Figures for "A nucleoside signal generated by a fungal endophyte regulates host cell death and promotes root colonization"

### *Si*NucA structure and sequence

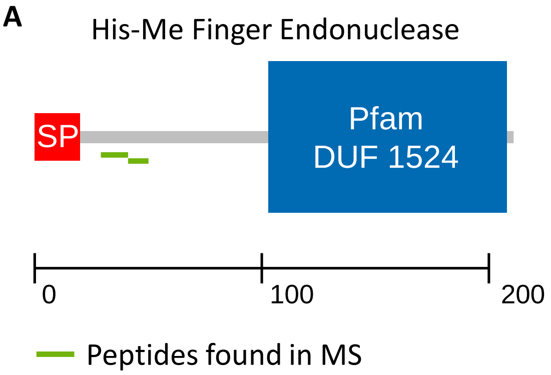


>PIIN02121_corrected_protein sequence
MIPTFSFAALILGMLATVSAAPLNLMESRALPTPVSVATAKTYLAALTIKAEANADTYDRDLFPHWITISGACNTRETVLKRDGTNVVTSSSCA
ATSGSWYSVYEGATQTSAGNIQIDHLVPLAEAWRSGASGWTTARRQAFANDLTRPQLIAVTGSVNQAKSDSDPAEWMPPLKSYYCTYVR
AWIQVKYYYSLSVDQAEYNALYKYLYAC

>PIIN02121_corrected_cDNA sequence
ATGATACCTACTTTCTCTTTTGCCGCCCTCATACTCGGCATGCTTGCAACCGTGAGCGCTGCTCCGCTCAACCTCATGGAGAGTCGTGC
ACTCCCGACCCCTGTTTCCGTGGCCACAGCAAAGACCTACCTCGCTGCTTTGACAATCAAGGCCGAGGCAAATGCGGACACTTATGAT
CGCGATTTATTCCCTCATTGGATCACCATTTCTGGCGCTTGTAACACGCGTGAGACGGTACTCAAGCGTGACGGAACGAATGTCGTGA
CGAGCAGCTCTTGCGCGGCTACCTCCGGCTCTTGGTATTCTGTCTATGAAGGTGCTACCCAGACATCTGCCGGCAATATTCAAATCGAC
CATCTTGTTCCATTAGCCGAGGCGTGGAGGTCTGGTGCATCTGGATGGACAACAGCTAGGAGACAAGCGTTTGCAAACGATCTCACTC
GTCCTCAACTCATCGCCGTAACCGGTAGCGTCAACCAGGCTAAGAGTGACAGTGATCCGGCTGAATGGATGCCACCCCTCAAGTCTTA
CTATTGCACTTATGTGCGCGCGTGGATTCAGGTCAAGTACTACTACTCGTTGTCGGTGGATCAGGCCGAGTACAATGCATTGTACAAG
TATTTGTACGCATGCTAG

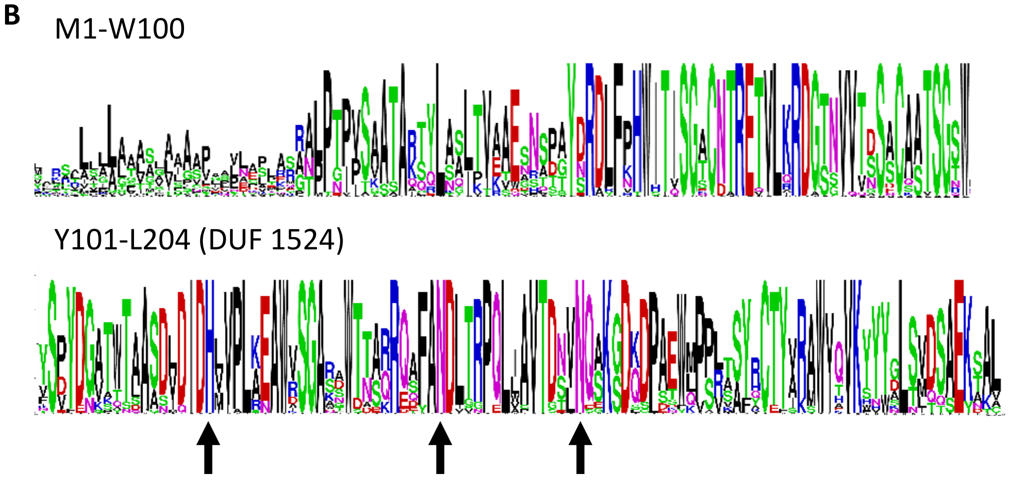


**Fig. S1: *Si*NucA structure and sequence**

(A) *Si*NucA (PIIN_02121) is 211 amino acids long, has a predicted signal peptide SP (first 20 amino acids) and the Pfam domain DUF 1524 belonging to the His-Me finger endonuclease superfamily. Two unique peptides (in green) were found in the apoplastic fluid (APF) of inoculated barley roots at 5 dpi by LC-MS/MS (Nizam et al., 2019). cDNA sequence was verified by rapid amplification of cDNA-ends with polymerase chain reaction (RACE-PCR), which revealed a 30 bp earlier start and a SNP at bp 33.
(B) Protein logo of multiple alignment of *Si*NucA and ten best protein BLAST hits for bacteria, Basidiomycetes and Ascomycetes each. Amino acids M1-W100 and Y101-L204 (DUF 1524 domain) shown. With the exception of the SP, the protein sequence is highly conserved. Arrows show the conserved HNN motif.

### *Si*NucA purification form culture filtrate

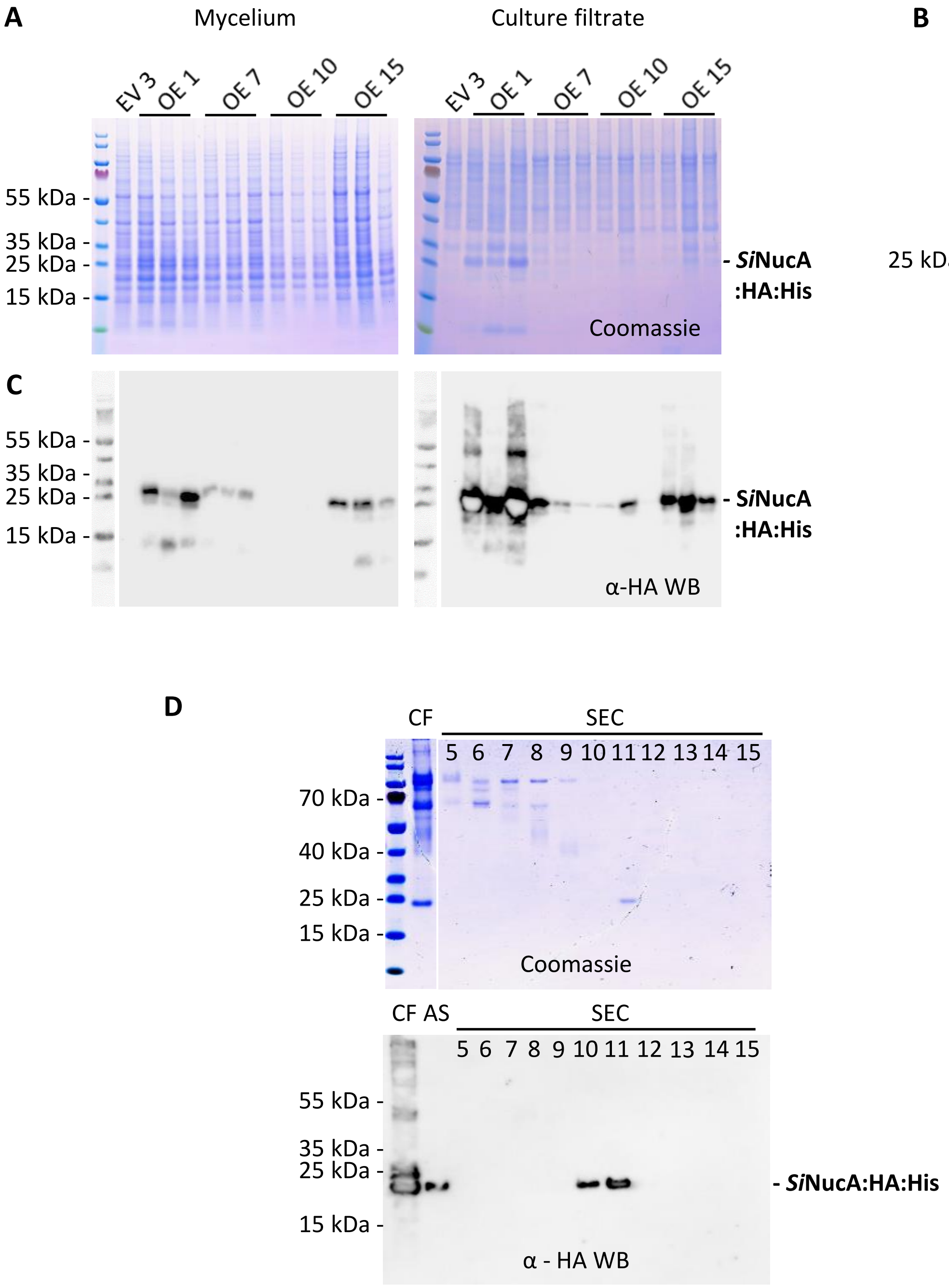


**Fig. S2: *Si*NucA:HA:His purification from the culture filtrate of *S. indica* overexpression strains**

(A) *S. indica Si*NucA:HA:His (OE 1, 7, 10, 15) and the empty vector strain 3 (EV 3) were grown for 5 days in CM medium followed by 2 days in MYP medium. Protein expression and secretion was analysed with mycelium (M) and culture filtrate (CF) on Coomassie-stained SDS-PAGEs and anti-HA Western blots (C).
(B) Control for intracellular protein contamination from M into CF with *S. indica* strain expressing cytosolic GFP processed in parallel.
(C) Anti-HA Western blots of *S. indica Si*NucA:HA:His strains from (A).
(D) *Si*NucA:HA:His protein enrichment from culture filtrate (CF) precipitated with 80% ammonium sulfate (AS) and separated by size exclusion chromatography (SEC, fractions 5- 15 shown). Fractions 10 and 11 were confirmed to contain *Si*NucA:HA:His by an anti-HA Western blot.

### Activity of purified *Si*NucA

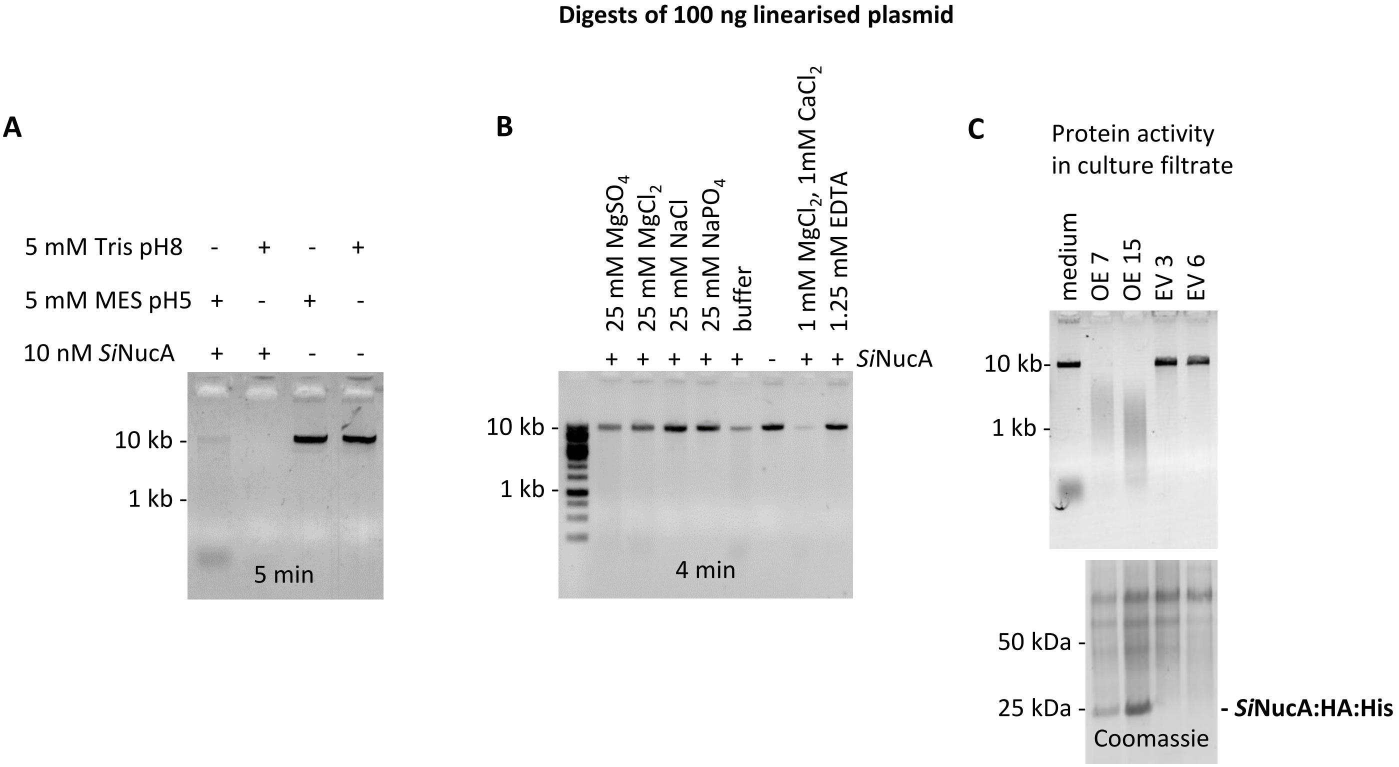


**Fig. S3: Activity of *Si*NucA purified from *S. indica* overexpression strains and culture filtrate.**

(A) 100 ng linearised plasmid added to 10 nM purified *Si*NucA in 5 mM Tris pH 8 or 5 mM MES pH 5 with 1mM $MgCl_2$, 1mM $CaCl_2$ and microelements incubated at RT for 5 min and loaded on an agarose gel.
(B) 100 ng linearised plasmid added to 10 nM purified *Si*NucA in buffer 5 mM Tris pH 8 supplemented with microelements. The influence of the addition of different salts and EDTA on protein activity was tested. The linearised plasmid in the different solutions with *Si*NucA was incubated for 4 min at RT and loaded on an agarose gel.
(C) Culture filtrate (CF) of *S. indica Si*NucA:HA:His overexpression strains OE 7 and OE 15 and empty vector strains EV 3 and EV 6 incubated with 100 ng linearised plasmid for 30 min and loaded on an agarose gel. 20 ml CF precipitated with trichloroacetic acid and loaded on SDS-PAGE as control for total protein amount in CF.

### Effect of *S. indica* colonization on *A. thaliana SiNucA* overexpression lines

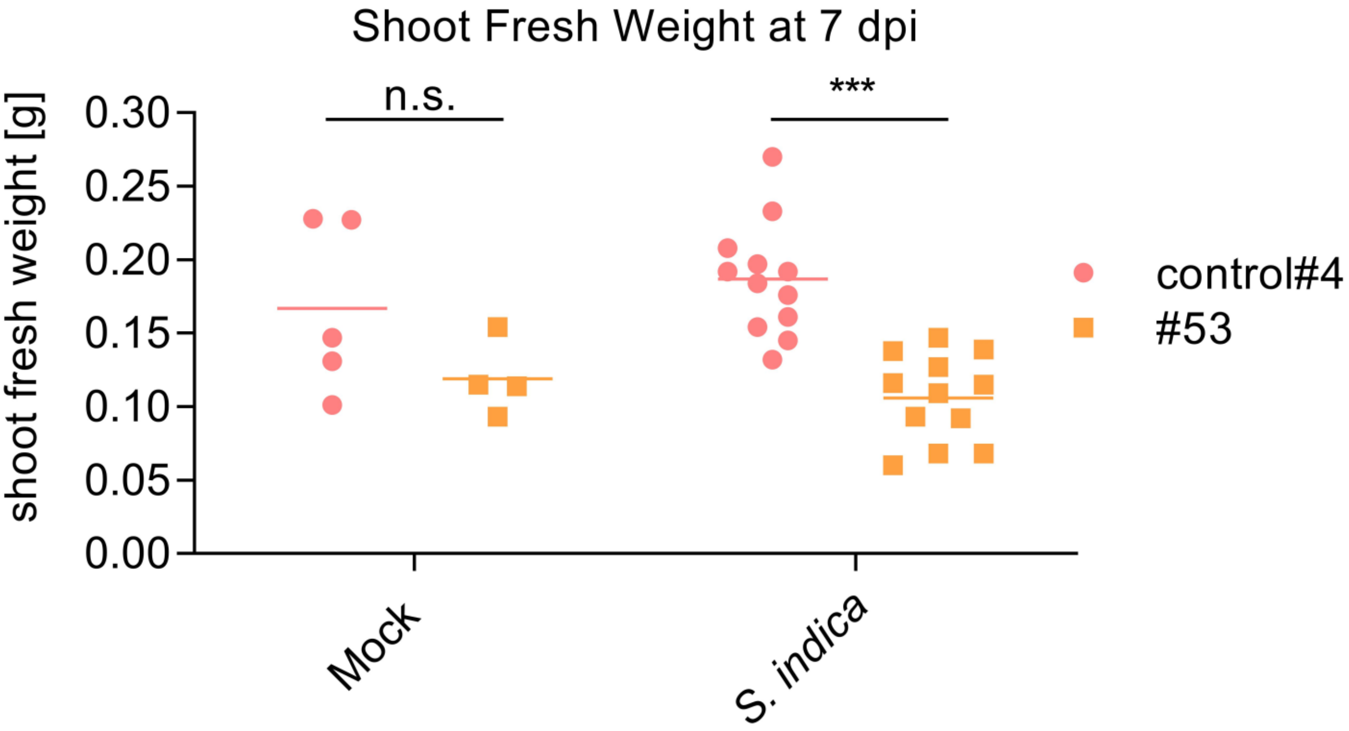


**Fig. S4: Effect of *S. indica* colonization on *SiNucA* overexpression lines**
Shoot fresh weight of *A. thaliana* seedlings overexpressing *SiNucA* (#53) and the corresponding control line (control#4). The data show the shoot fresh weight of *S. indica* and mock-inoculated seedlings at 7 dpi. Data points depict independent biological replicates. Asterisks represent significant difference between different genotypes analyzed by Student's t-test - $p < 0.005$ (***).

### Localization of *Si*NucA:mCherry in *A. thaliana* roots

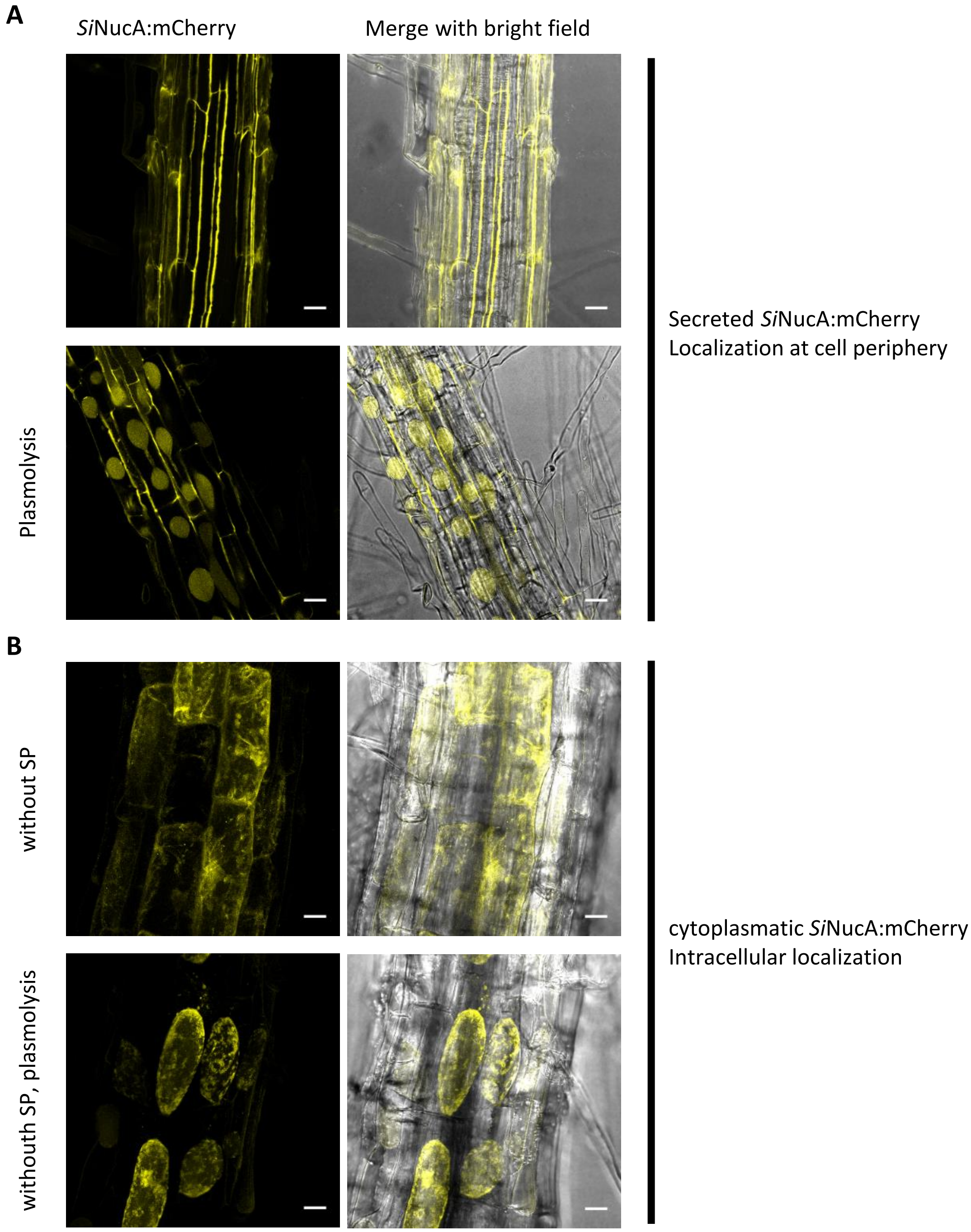


**Fig. S5: Localization of *Si*NucA in Arabidopsis heterologously expressing *Si*NucA:mCherry with and without signal peptide**

CLSM live cell imaging of Arabidopsis roots.
(A) Arabidopsis root expressing *Si*NucA:mCherry with and without plasmolysis.
(B) Arabidopsis root expressing *Si*NucA (w/o SP):mCherry with and without plasmolysis. Maximum projection of z-stacks.
Plasmolysis was achieved using 1 M sorbitol or NaCl. Bars = 20 µm. CLSM microscopy was repeated with 3 independent samples.

### Localization of *Si*NucA:mCherry in *A. thaliana* roots

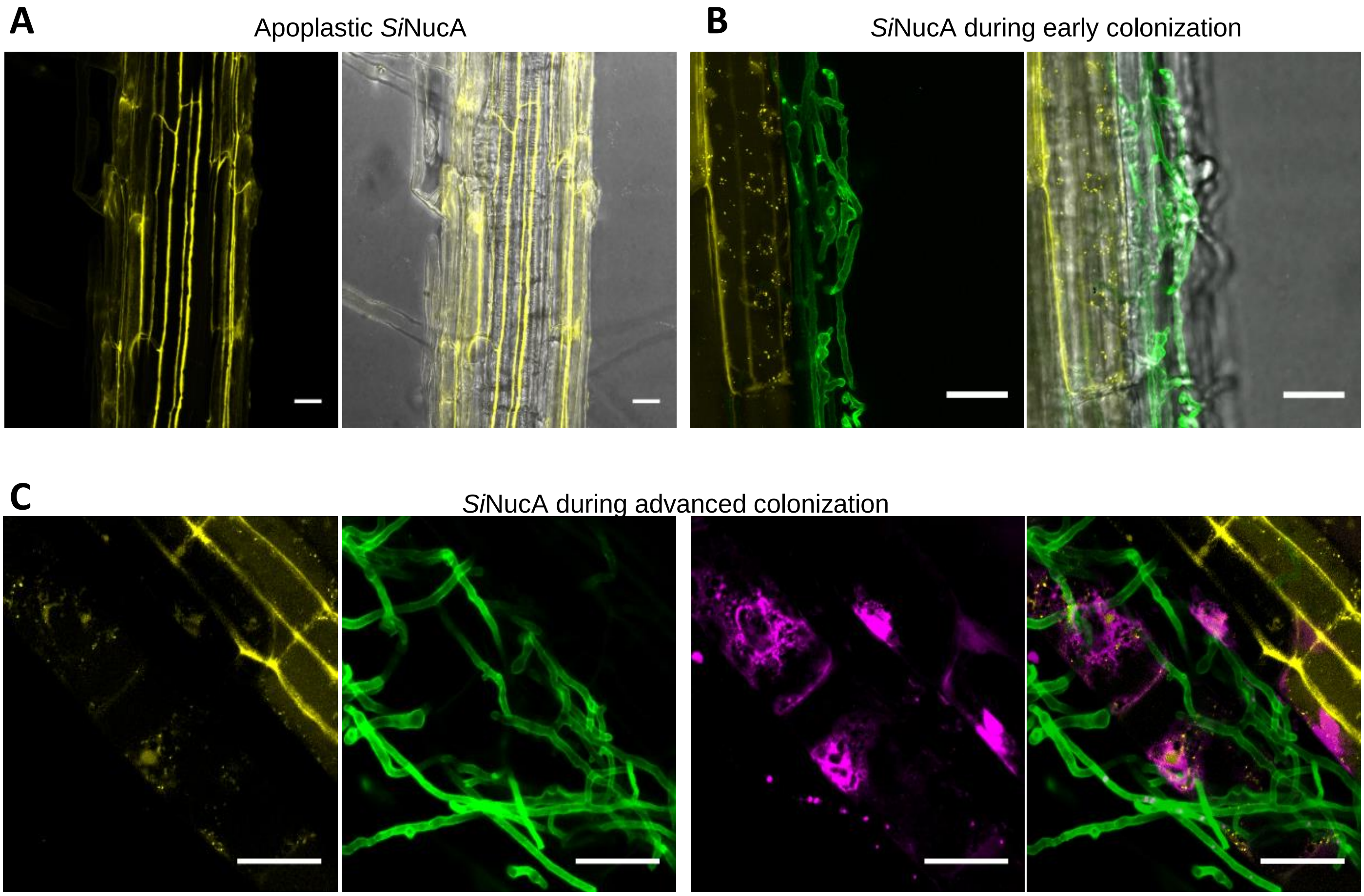


**Fig. S6: Time course of localization of *Si*NucA in Arabidopsis heterologously expressing *Si*NucA:mCherry.**

CLSM live cell images of Arabidopsis root expressing *Si*NucA:mCherry (yellow) inoculated with *S. indica*. Fungal cell wall of hyphae stained with WGA-AF 488 (green) and nuclei with DAPI (magenta).
(A) *Si*NucA:mCherry accumulates in the apoplast in non-inoculated roots.
(B) *Si*NUCA:mCherry accumulates around penetrating hyphae (intracellular hyphae in living cells not stainable with WGA-AF 488 in the biotrophic interaction phase).
(C) In a later phase, *Si*NUCA:mCherry is mainly present in nuclei of colonized cells.
Bars = 20 µm.

### Identification of products upon *Si*E5NT incubation

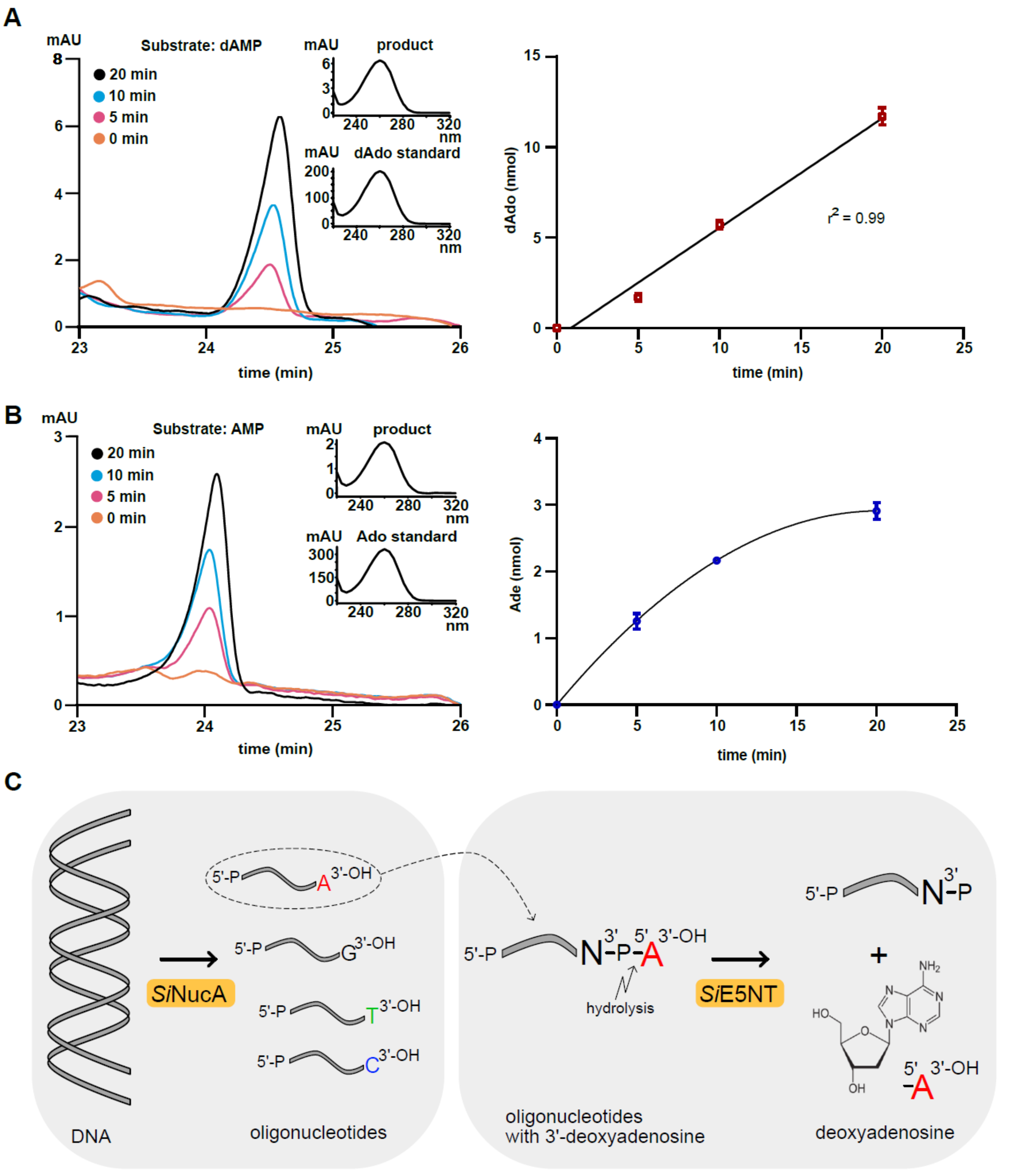


**Figure S7: Analysis of metabolites derived from incubation of *Si*E5NT with deoxyadenosine monophosphate (dAMP) or adenosine monophosphate (AMP) and model of the reaction mechanism.**

(A) Left panel, HPLC trace from spectrometric detection at 254 nm for deoxyadenosine (dAdo) derived from dAMP converted by *Si*E5NT after the indicated incubation times. 1 mM dAMP was incubated with 1.9 x $10^{-4}$ mg protein in 100 µl. The insert compares the spectra of standard and reaction product. Right panel, dAdo generated after 0, 5, 10 and 20 min of incubation (*n* = 3). The data was fitted by linear regression.
(B) Same as in (A) but using AMP as a substrate (*n* = 3).
(C) Model of the reaction mechanism of *Si*NucA and *Si*E5NT with DNA as substrate, showing preference for dAdo production.

### Expression of cell death marker genes after dAdo treatment

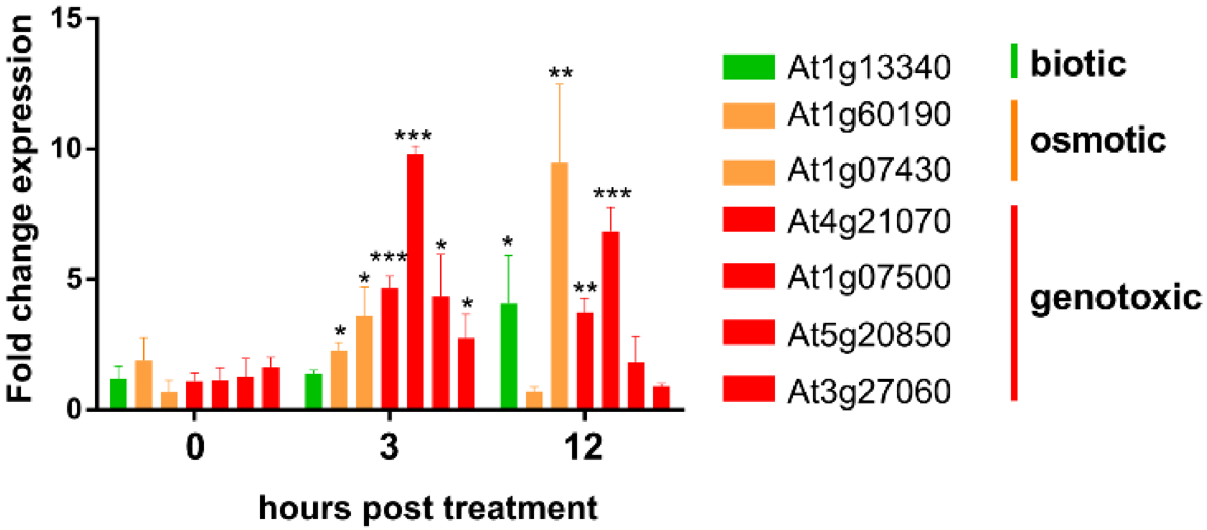


**Fig. S8: Relative expression of cell death marker genes in Arabidopsis seedlings after 500 µM dAdo treatment measured by qRT-PCR.**

Expression values relative to mock-treated seedlings were calculated using the ΔΔCt-method. Bars represent the average of three independent biological replicates while error bars represent the standard deviation. Asterisks represent significant difference to the mock-treated sample analyzed by Student's t-test. $p < 0.05$ (*), $p < 0.01$ (**), $p < 0.005$ (***).Marker genes were selected based on Olvera-Carrillo et al., 2015)

### dAdo-triggered cell death is concentration-dependent

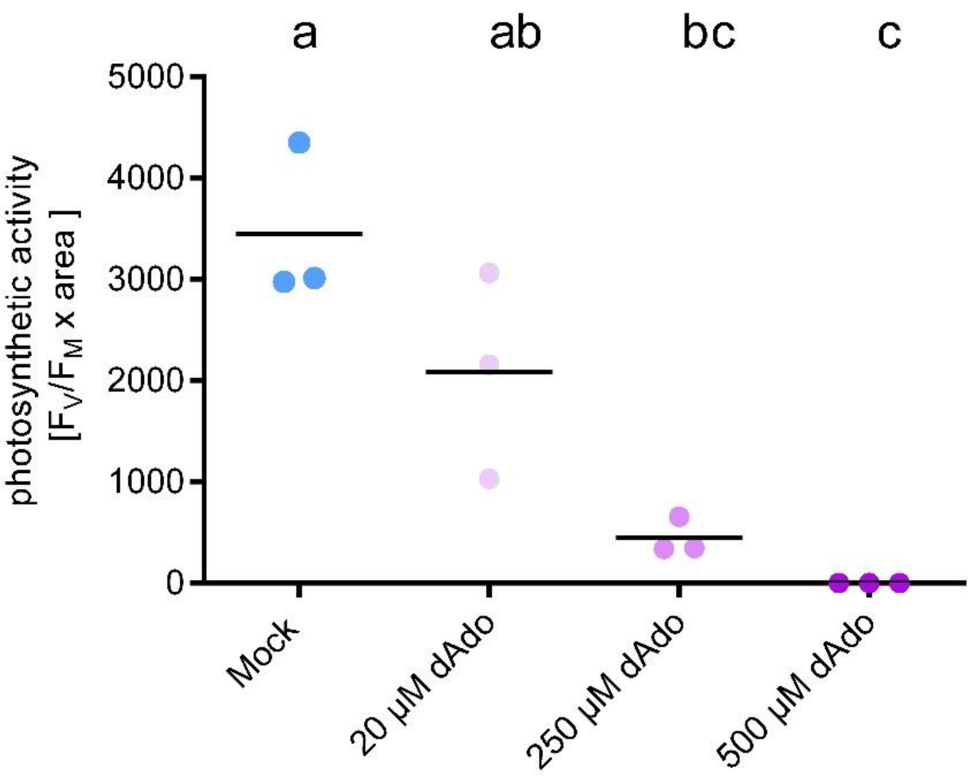


**Fig. S9: dAdo-triggered cell death is concentration-dependent.**

Photosynthetic activity ($F_V/F_M$ multiplied with corresponding photosynthetically active area) of 9-day-old Col-0 seedlings incubated with different concentrations of dAdo at 7 dpt. Dots represent 3 biologically independent replicates. Different letters indicate significant different groups determined with a one-way ANOVA with post-hoc Tukey HSD test ($p<0.05$).

### Recovery after dAdo treatment

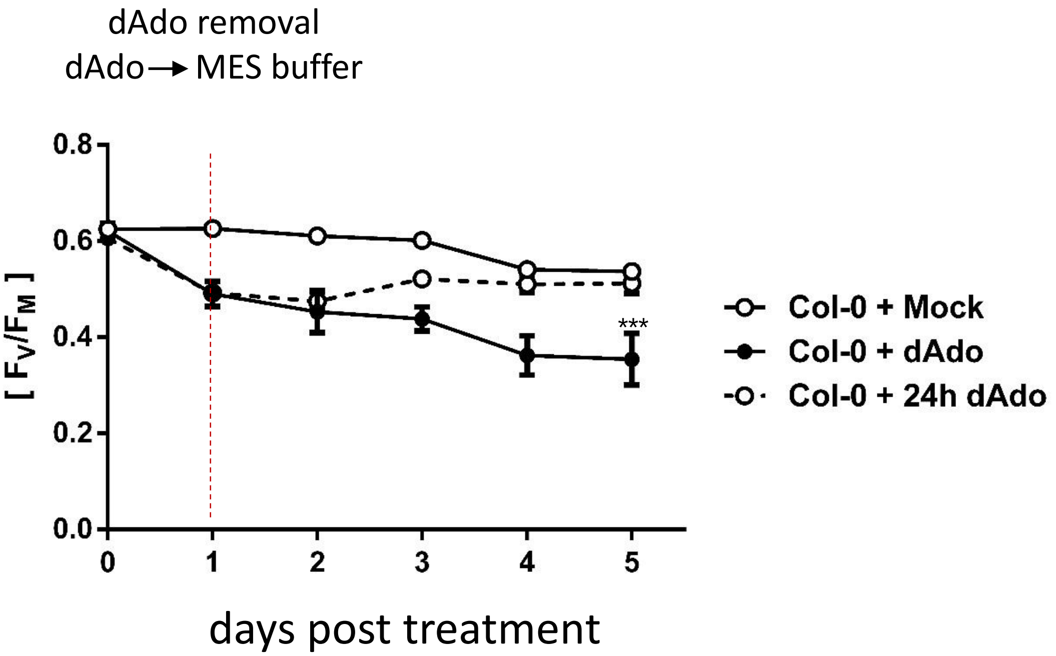


**Fig. S10: The dAdo-induced cell death process is reversible.**

Photosynthetic activity ($F_V/F_M$) of 7-day-old Col-0 seedlings incubated with 500 µM dAdo or 2.5 mM MES buffer (pH = 5.6). After 24 hours, the dAdo containing solution was replaced with MES buffer (Col-0 + 24 h dAdo). Dots represent 12 biological replicates, while error bars represent the SEM. Asterisks represent significant differences to the buffer-treated sample analyzed by Student's t-test. $p < 0.005$ (***).

*Si*E5NT expression in *N. benthamiana* leaves after agroinfiltration

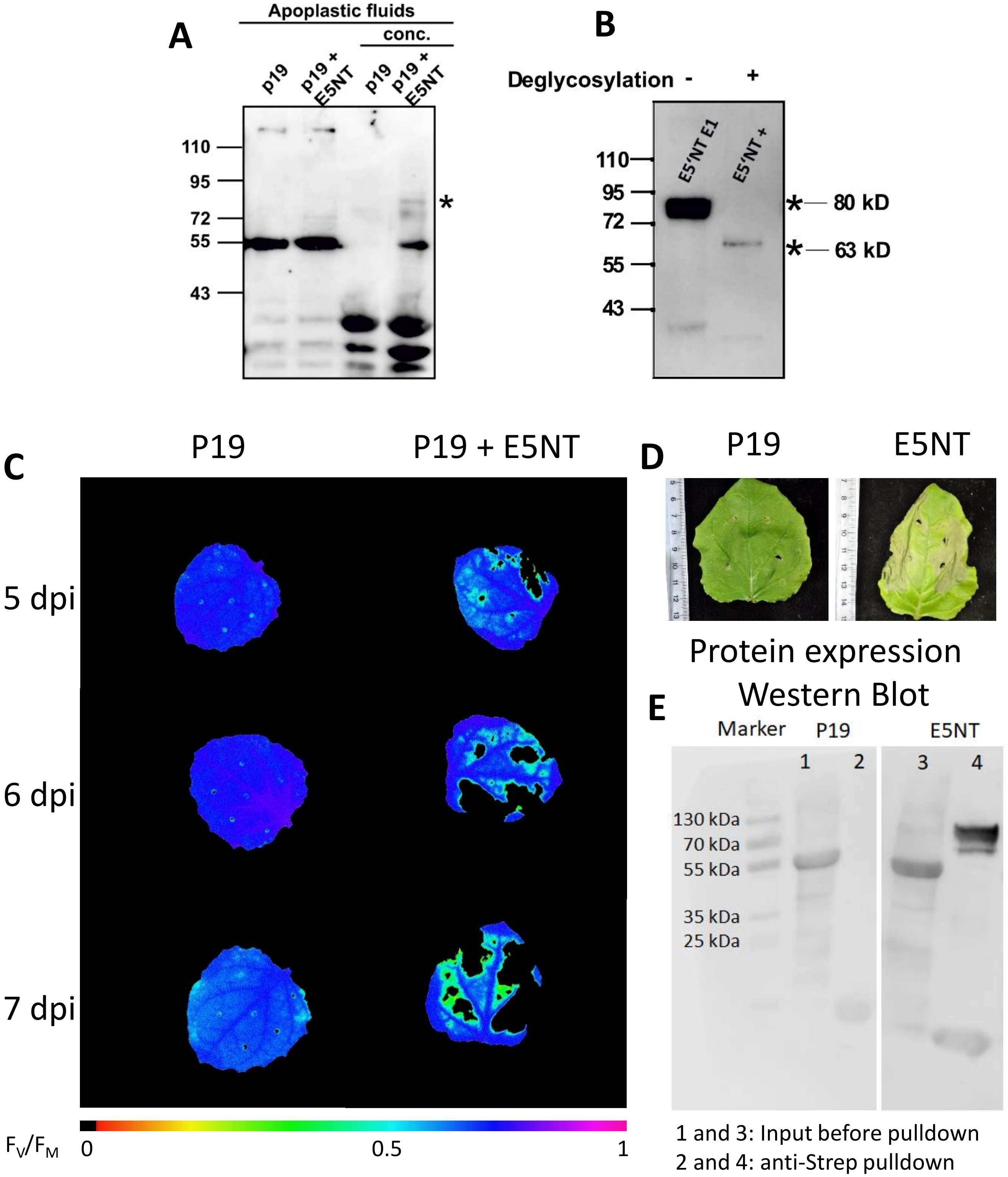

**Fig. S11: Transient *Si*E5NT expression and secretion in *N. benthamiana.***

(A) *Si*E5NT is detected in the apoplastic fluids of *Nicotiana benthamina* leaves after transcient overexpression using Agrobacterium. The leaves of *N. benthamiana* were infiltrated with Agrobacterium carrying either p19 or E5NT expression construct. To validate secretion of *Si*E5NT, the apoplastic fluids were collected from the leaves at 4 dpi. The proteins in the apoplastic fluids were concentrated with 100% acetone and were separated via SDS-PAGE. The StrepII-tagged-*Si*E5NT in the apoplastic fluids were detected via Western Blot using the StrepMAB antibody.

(B) Deglycosylation of *Si*E5NT overexpressed in the leaves of *N. benthamiana* leaves from (A). The purified *Si*E5NT was treated with protein deglycosylation mix and incubated 37 °C for 16 hours. The deglycosylated *Si*E5NT was separated using SDS-PAGE and blotted onto nitrocellulose membrane. The StrepII-tagged *Si*E5NT in the apoplastic fluids were detected using the StrepMAB antibody.

(C) Visualization of $F_V/F_M$ measured via PAM fluorometry. Blue indicates a high value while green decreased and yellow strongly decreased photosynthetic activity. Depicted are leaves of *N. benthamiana* heterologously expressing p19 or a combination of p19 and E5NT after Agrobacterium infiltration.

(D) Image of leaves of *N. benthamiana* heterologously expressing p19 or a combination of p19 and E5NT after Agrobacterium infiltration.

(E) Protein expression in *N. benthamiana* leaves. Proteins were extracted from infiltrated *N. benthamiana* leaves using Anti-Strep-beads for pulldown and analyzed by Western Blot using an anti-Strep antibody. Lane 1 and 3 represent raw extract prior to pulldown, whereas lane 2 and 4 represent pulled-down proteins. E5NT = 80 kDa.

### Heterologous *Si*E5NT expression in *N. benthamiana* leaves after agroinfiltration

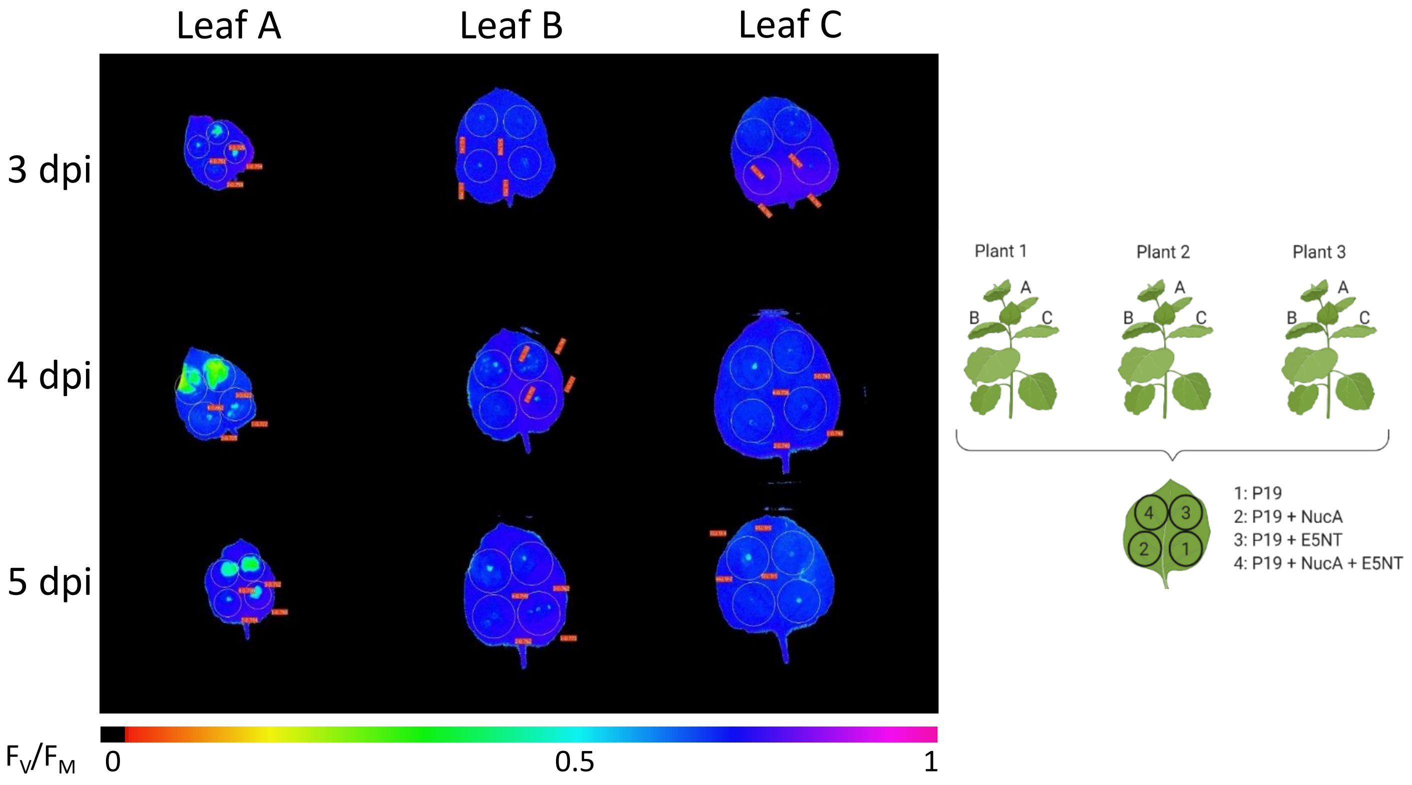


**Fig. S12: Heterologous expression of E5NT in *N. benthamiana* leaves induces cell death in younger leaves.**

Visualization of $F_V/F_M$ measured via PAM fluorometry. Blue indicates a high, while green indicates decreased and yellow strongly decreased photosynthetic activity. Depicted are leaves of *N. benthamiana* infiltrated with different mixtures of *Agrobacterium tumefaciens* strains expressing: 1 = p19, 2 = p19 + NucA, 3 = p19 + E5NT, 4 = p19 + NucA + E5NT. Photosynthetic activity was measured in three leaves of different ages (A to C = younger to older) originating from three different plants at each time point.

### dAdo-induced cell death in *N. benthamiana*

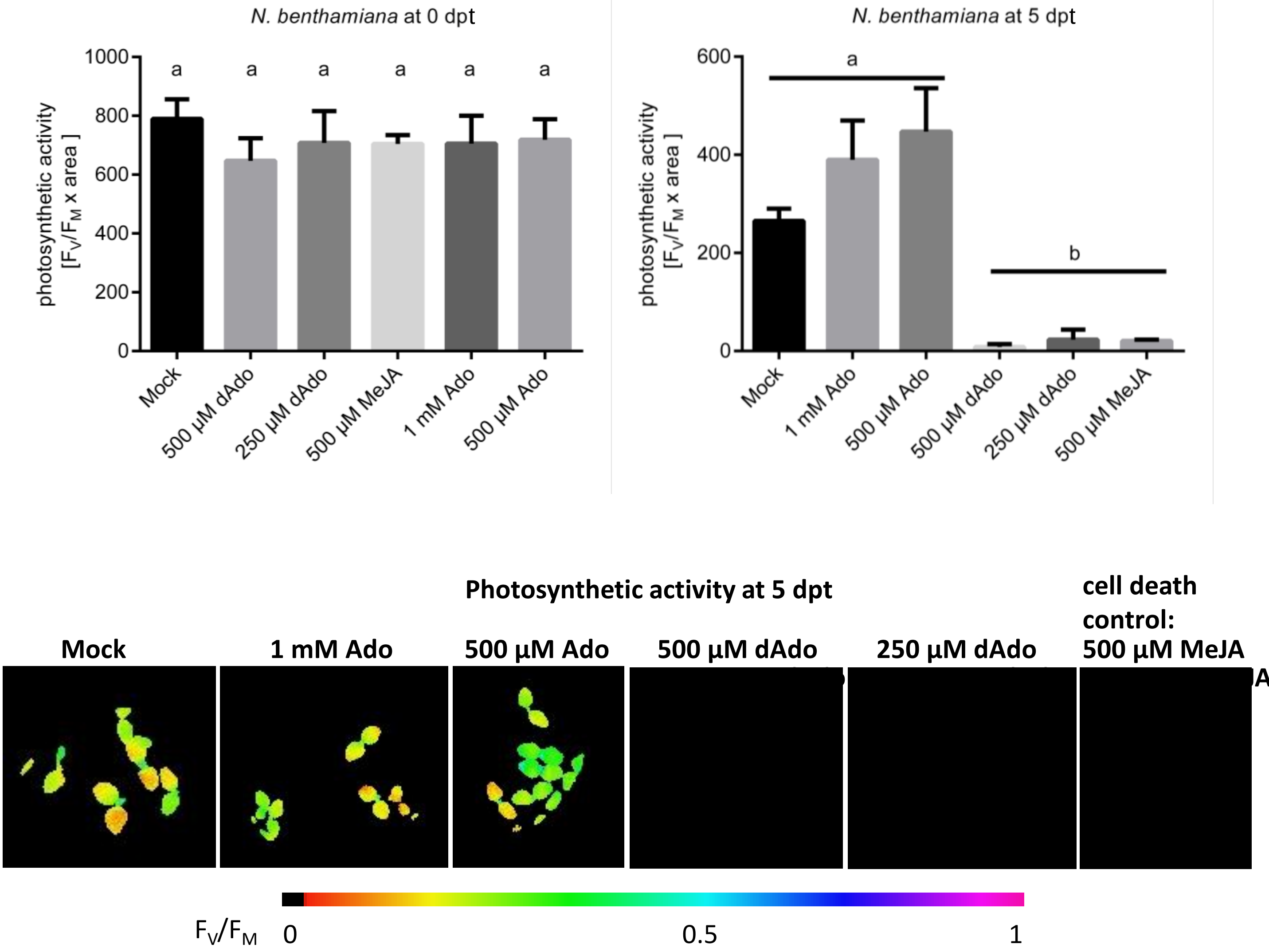


**Fig. S13: dAdo induces cell death in *N. benthamiana.***

Photosynthetic activity ($F_V/F_M$ x photosynthetic area) of 7-day-old *N. benthamina* seedlings incubated with dAdo, Ado, MeJA (cell death control) or Mock (2.5 mM MES, pH = 5.6) at 0 and 5 dpt. Bars represent the average of four independent biological replicates, while error bars represent the SEM. Letters indicate significant different groups determined with a one-way ANOVA with post-hoc Tukey HSD test ($p<0.05$).

### dAdo induces cell death in the liverwort *Marchantia polymorpha*

#### TAK1 (male):

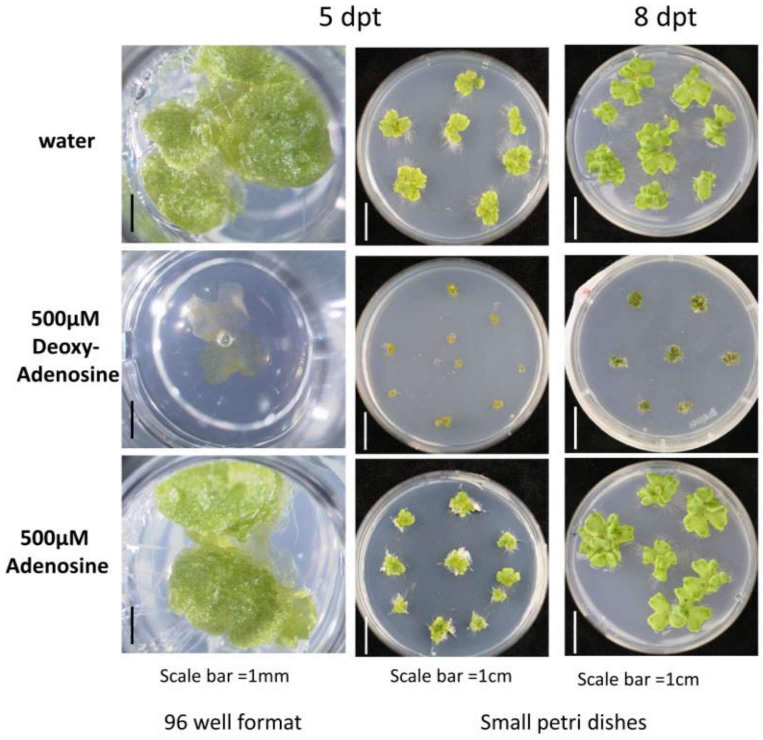


#### TAK2 (female):

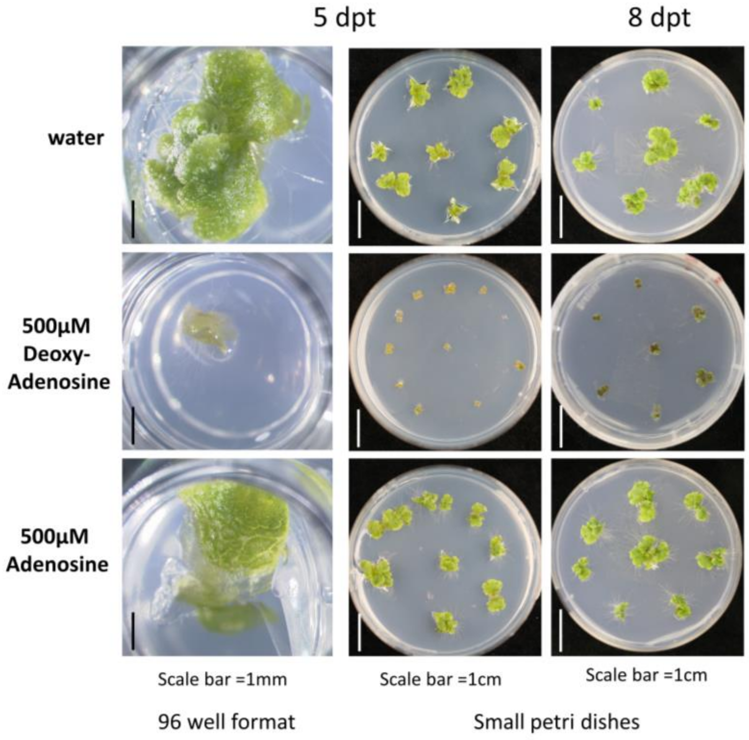


**Fig. S14: dAdo induces cell death in *Marchantia polymorpha*.**
Gemmae of *Marchantia polymorpha* Tak-1 and Tak-2 were cultured on half-strength B5 medium* under continuous light at 22°C for 5 days. (*1/2 Gamborg B5 salt mixture 1,5g/l, MES 0,5g/l, sucrose 10g/l, plant agar 10g/l, pH5.2 set with KOH; containing 500µM dAdo, 500µM Ado or water). The phenotype was monitored at 5 days post transfer (column 1 and 2) and at 8 days post transfer (column 3).

### MEcPP does not induce cell death

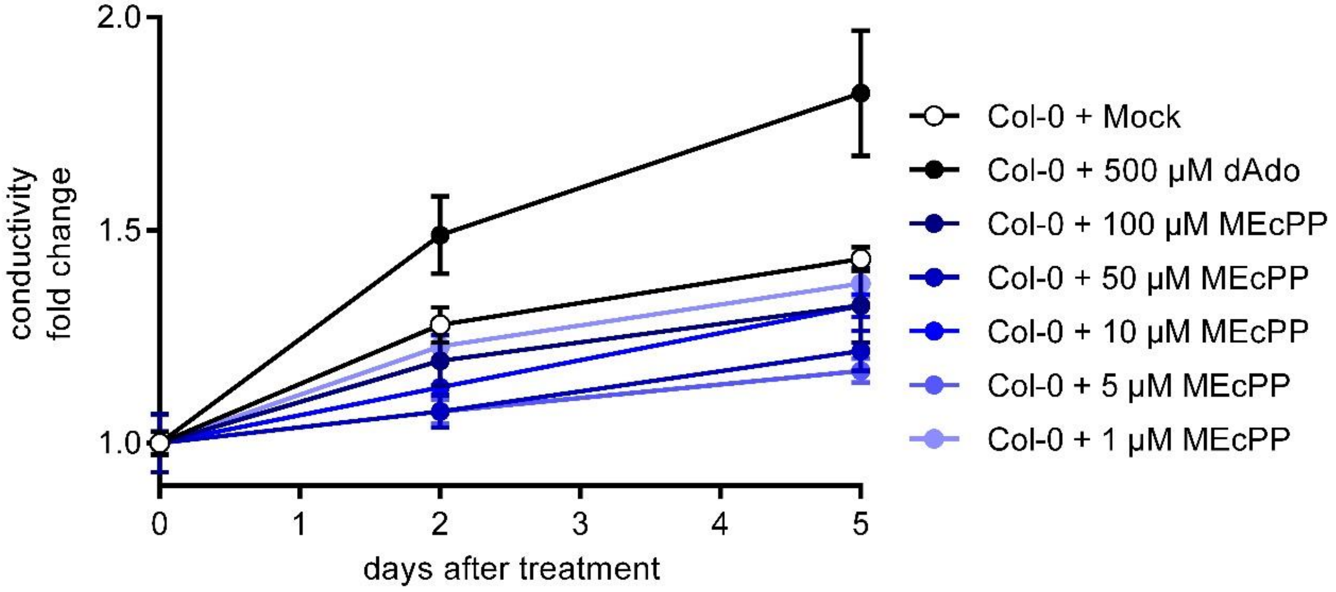


**Fig. S15: MEcPP does not induce cell death.**

Conductivity of MES buffer containing Col-0 seedlings (9-day-old) after Mock-, dAdo or MEcPP treatment. Data points depicts the mean, while error bars depict the standard error of the mean (SEM) obtained from 3 biological independent replicates.

### Purine derivates do not trigger a ROS burst

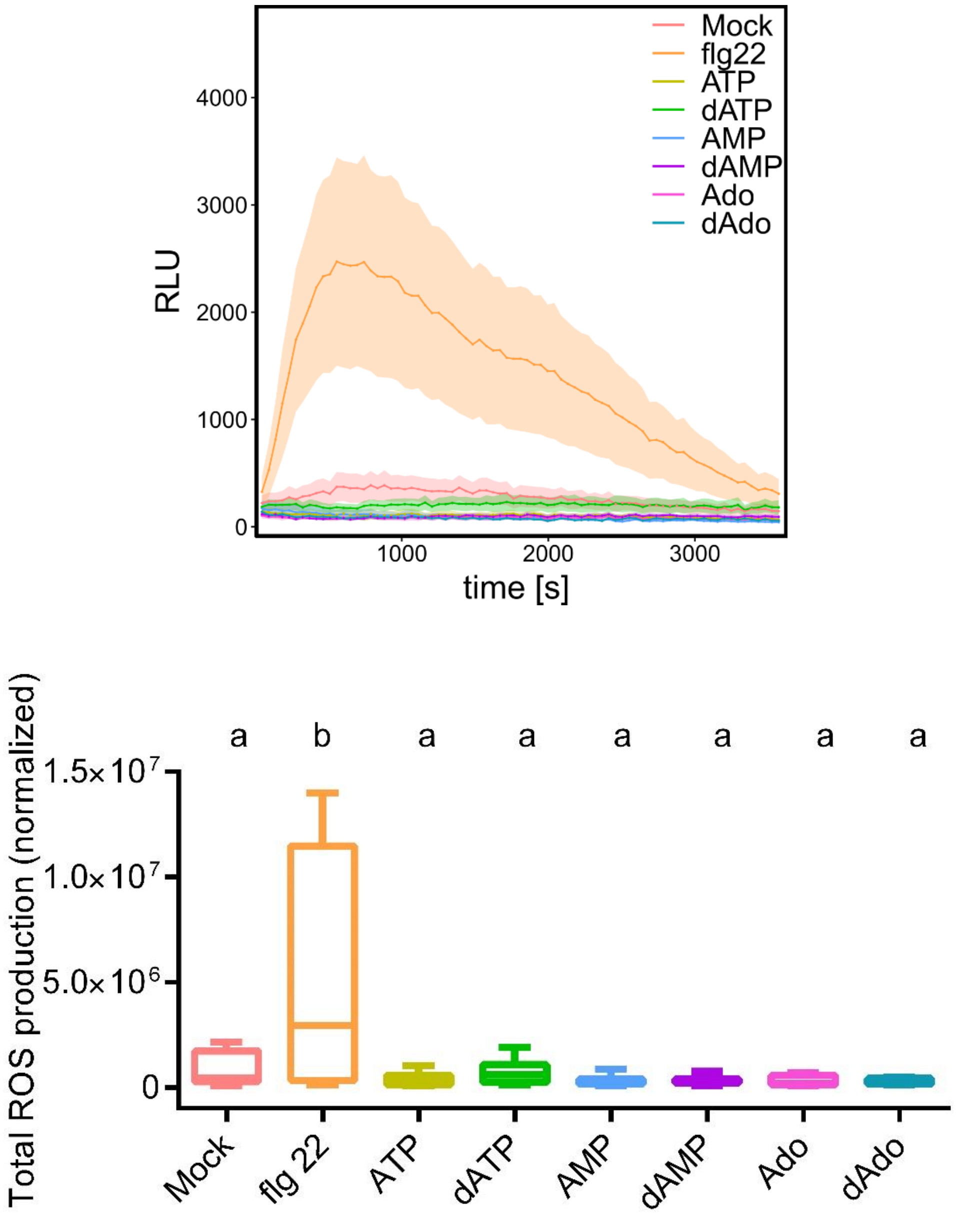


**Fig. S16: Purine derivatives do not trigger a ROS burst.**

A) Apoplastic ROS production after treatment of seven-day-old *A. thaliana* Col-0$^{AEQ}$ seedlings with 500 µM purine derivatives solved in 2.5 mM MES buffer (pH 5.7 buffer). Buffer and 33 µM flg22 were used as controls. ROS production was monitored via a luminol-based chemiluminescence assay. The curve represents eight biological replicates. The experiment was repeated three times with similar results.
B) Boxplots represent total ROS production over the measured time period. Values represent eight biological replicates, each containing one seedling. Different letters indicate significant different groups determined with a one-way ANOVA with post-hoc Tukey HSD test ($p<0.05$).

### *S. indica* growth on dAdo-containing medium

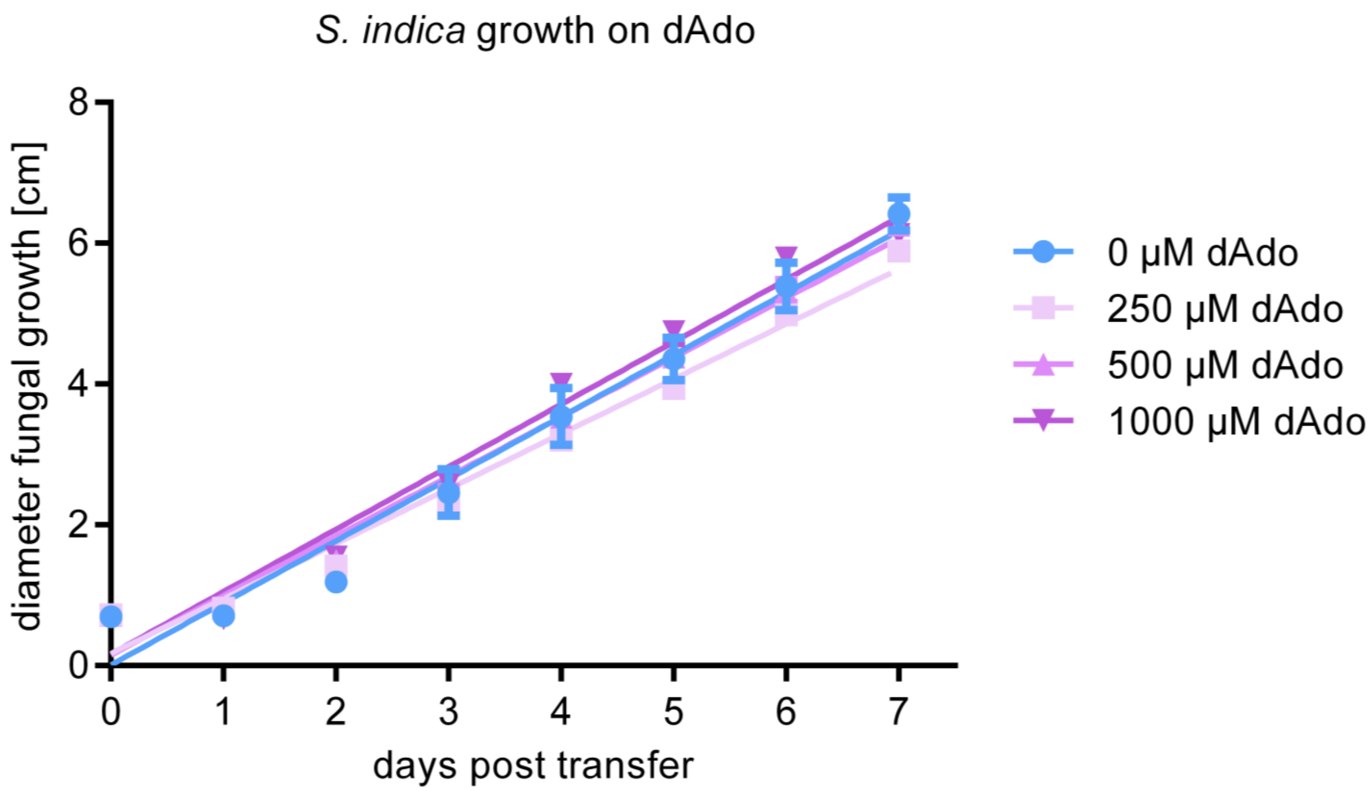


**Fig. S17: Growth of *S. indica* on CM medium containing different concentrations of dAdo.**

Dots represent the average diameter of 3 technical replicates while error bars represent the SD. Lines represent the linear regression describing the fungal growth calculated from the raw data. The experiment was repeated thrice showing comparable results (Data not shown). Growth was measured extending from the agar plug (diameter 0.7 cm).

### *ent3* but not *ent1 A. thaliana* knockout mutant shows reduced ion leakage after dAdo treatment

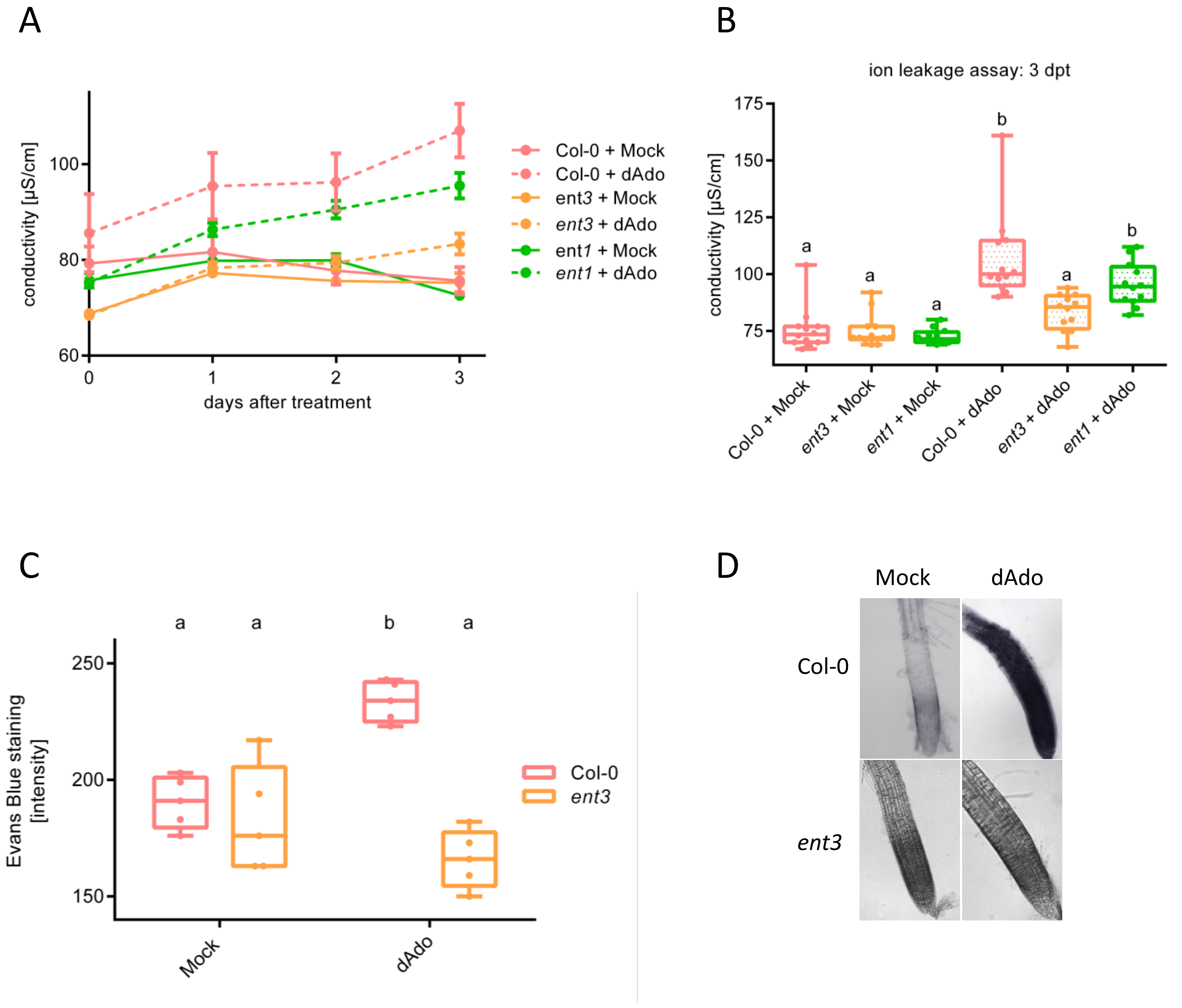


**Fig. S18: Reduced cell death in *ent3* after dAdo treatment.**

A) Conductivity of 2.5 mM MES buffer containing Arabidopsis seedlings (9-day-old) after Mock- or dAdo-treatment. Data points depicts the mean, while error bars depict the standard error of the mean (SEM) obtained from 12 biological replicates. Asterisks represent significant difference between dAdo-treated Col-0 and *ent3* analyzed by Student's t-test. The experiment was repeated three times with similar results.
B) Conductivity of samples from A at 3 dpt. Different letters indicate significant different groups determined with a one-way ANOVA with post-hoc Tukey HSD test ($p<0.05$). C) Quantification of root cell death in Arabidopsis roots at 4 days after dAdo-treatment. Cell death quantity was evaluated via Evans Blue staining and normalized to the mock-treated roots. Boxplots represent the data obtained from 5 biological replicates. Different letters indicate significant different groups determined with a one-way ANOVA with post-hoc Tukey HSD test ($p<0.05$). (D) Evans Blue stained roots tips from (C). The experiment was repeated three times with similar results.

### *ent3* shows a transient decrease in *S. indica* colonization

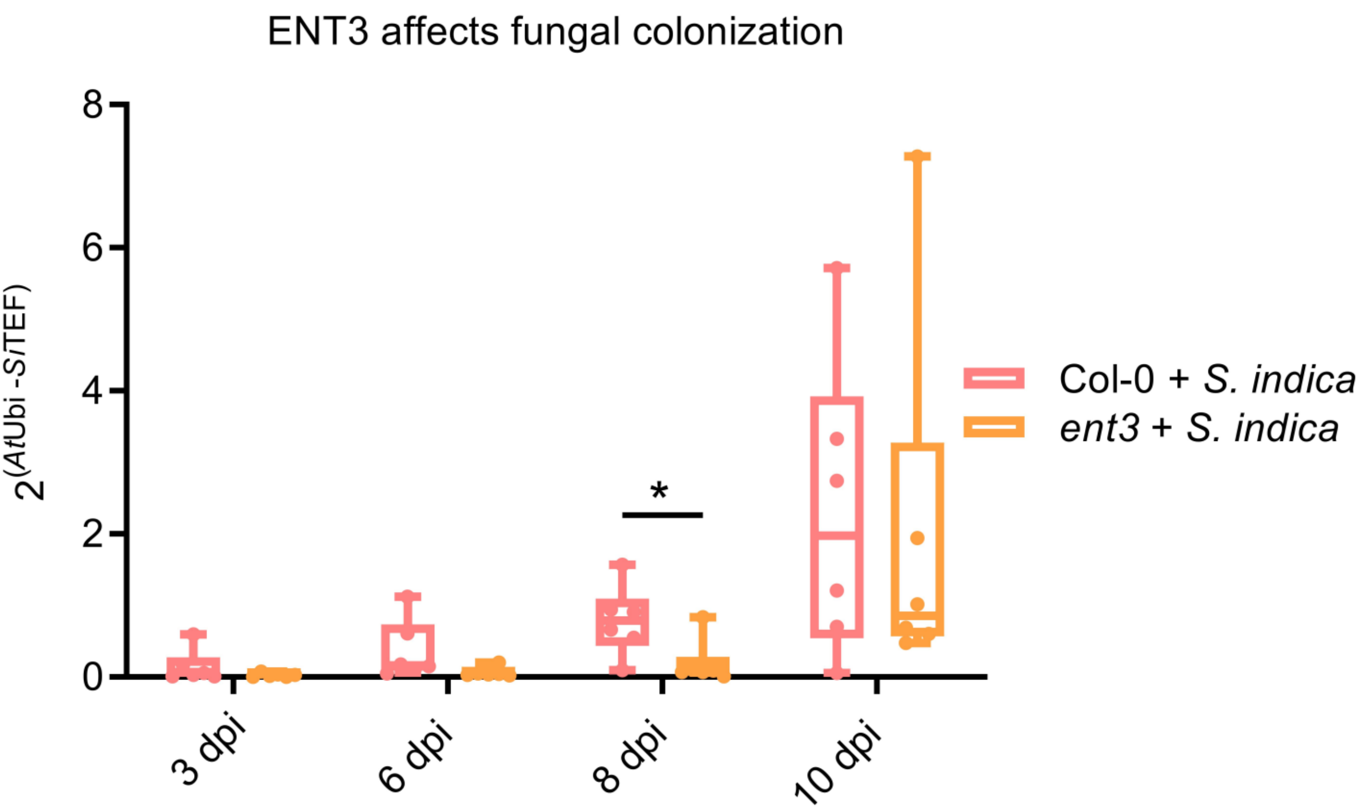


**Fig. S19: *ent3* shows a transient decrease in *S. indica* colonization**
*S. indica* abundance in 7-day-old Arabidopsis seedlings. The ratio of fungus (*Si*TEF) to plant (*At*Ubi) was calculated using cDNA as template and the $2^{-\Delta CT}$ method. Boxplots represent 6 independent biological replicates. Asterisks indicate significant difference from Col-0 samples (Student's t-test, $p < 0.05$ *).

### Screening of a *A. thaliana* T-DNA library

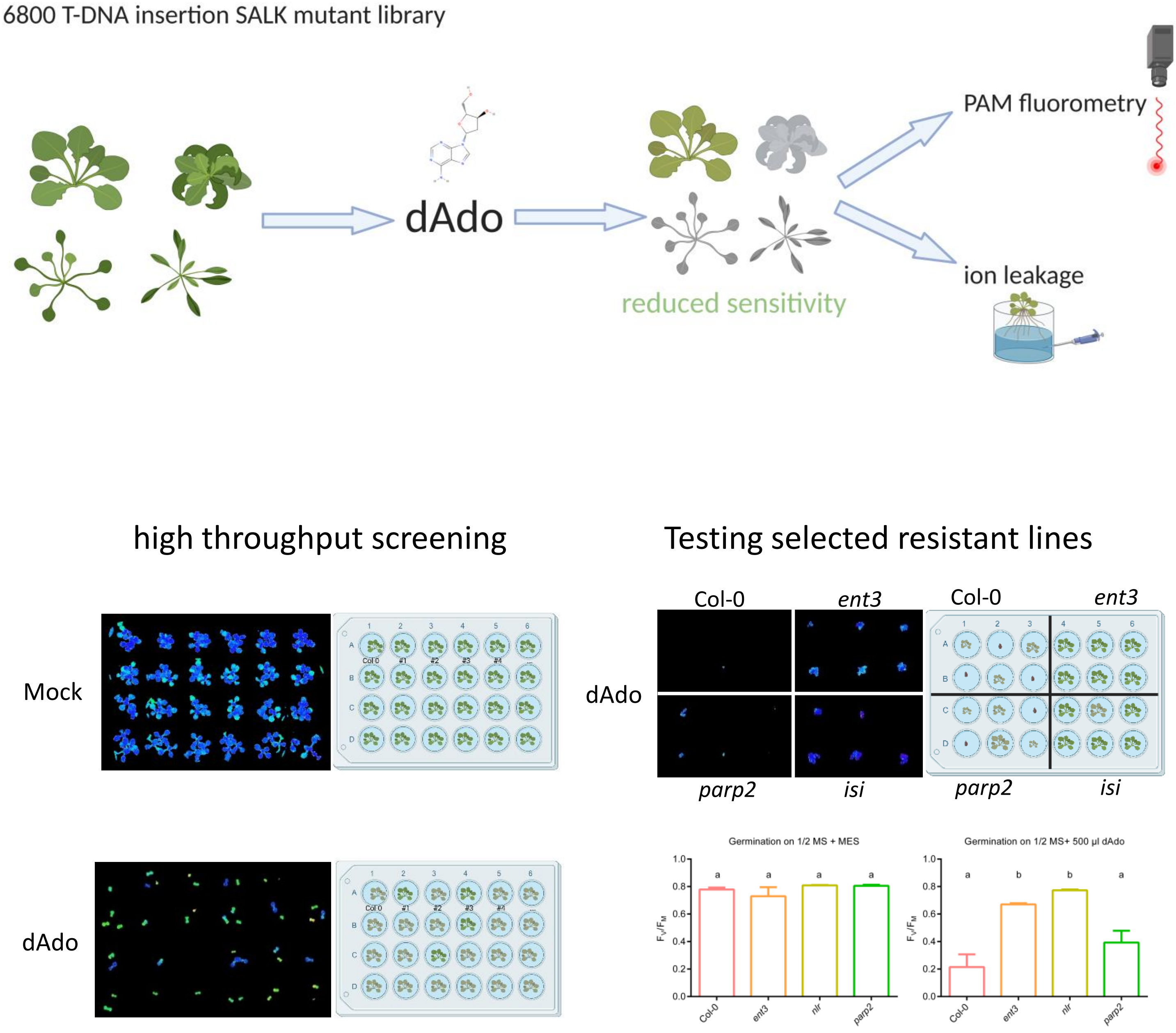


**Fig. S20: Screening of an *A. thaliana* library of T-DNA insertion mutant lines for alternation in dAdo-sensitivity.**

The model shows our screening process to identify mutants affected in dAdo-sensitivity. We screened over 6800 T-DNA insertion SALK lines for an altered response to dAdo. For high-throughput screening the mutant lines were analyzed via PAM fluorometry on 24 well plates, including Col-0 WT as control in each plate. Arabidopsis mutant lines with a resistant phenotype to dAdo were subsequently analyzed in ion leakage assays. The screening was repeated 3 times. Mutants that showed reproducible resistant phenotypes in the high-throughput screening were analyzed by germination assays on solid plant media with and without dAdo additionally tested as described in material and methods (see also main figures 5, 7 and 8).

### *isi* SALK –DNA insertion line

- **Gene:** AT5G45240
- **SALK line:** SALK_034517C

**A**

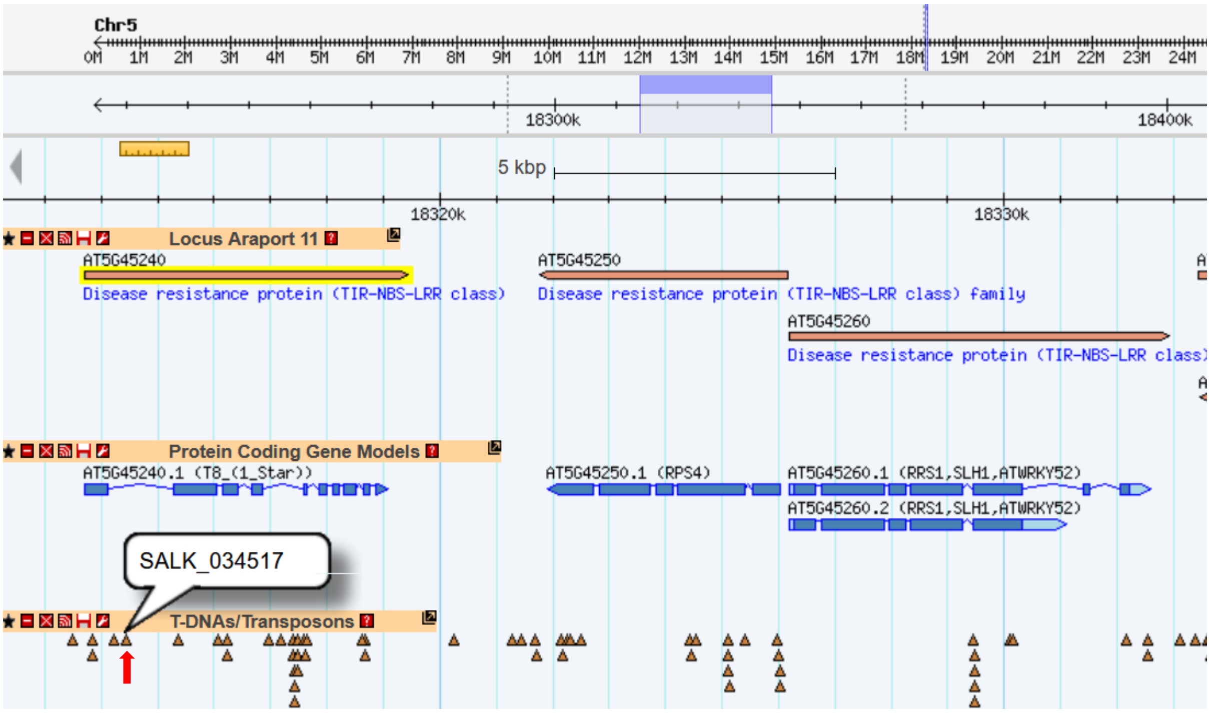


**B**

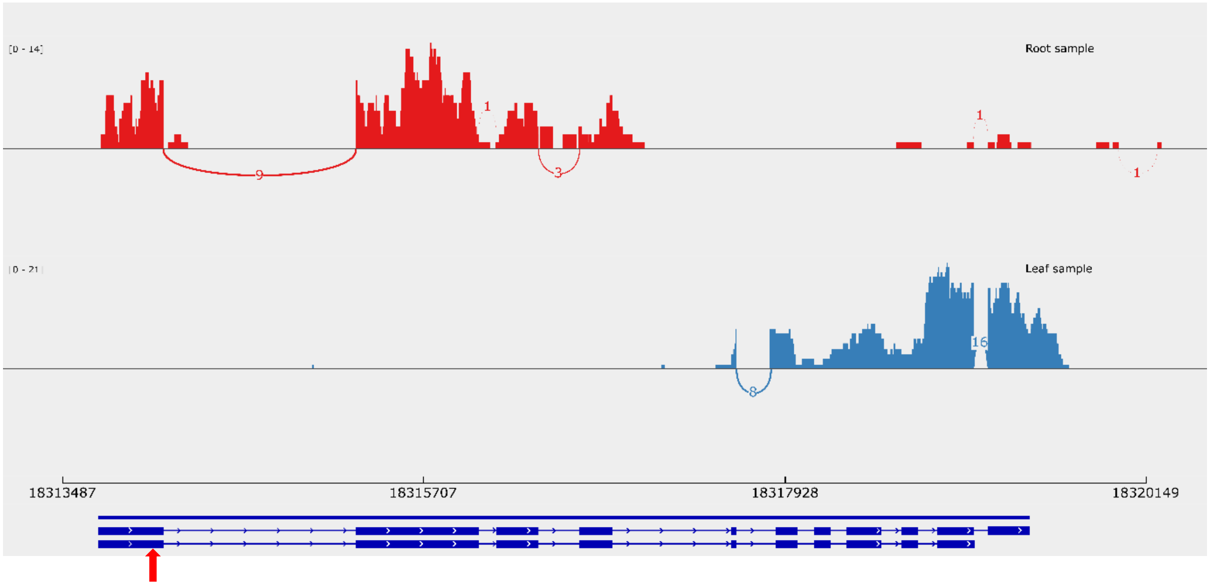


**Fig. S21: Gene locus of NLR gene and Exon Mapping.**
A) Figure and data were taken from https://gbrowse.arabidopsis.org of The Arabidopsis Information Resource (*TAIR*). The red arrow marks the T-DNA insertion site for SALK_034517C.
B) The sashimi plot shows the reads coverage of exons and the number of reads supporting splice junctions of the Arabidopsis AT5G45240 gene from Araport11 annotation in leaf and root samples.
The data shown were downloaded from the Sequence Read Archive (SRA) of NCBI database for Arabidopsis leaf tissue (PRJNA336058) and root tissue (PRJNA344473). Arabidopsis mock samples were analyzed for both datasets. The three biological replicates of each tissue were merged together.
Raw reads were trimmed with fastp and trimmed reads were aligned to *A. thaliana* reference genome TAIR9. The mapped reads were visualized with the Integrative Genomics Viewer (IGV).
The Sashimi plot was displayed in IGV by setting the minimum junction depth to 4.

### Expression of *A. thaliana* NLRs in different cell types

**A**

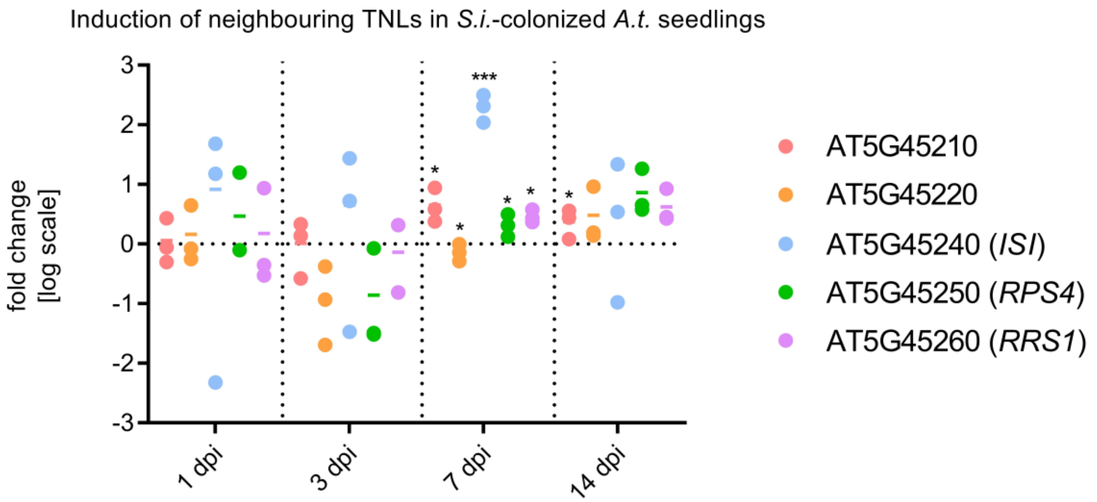


**B**

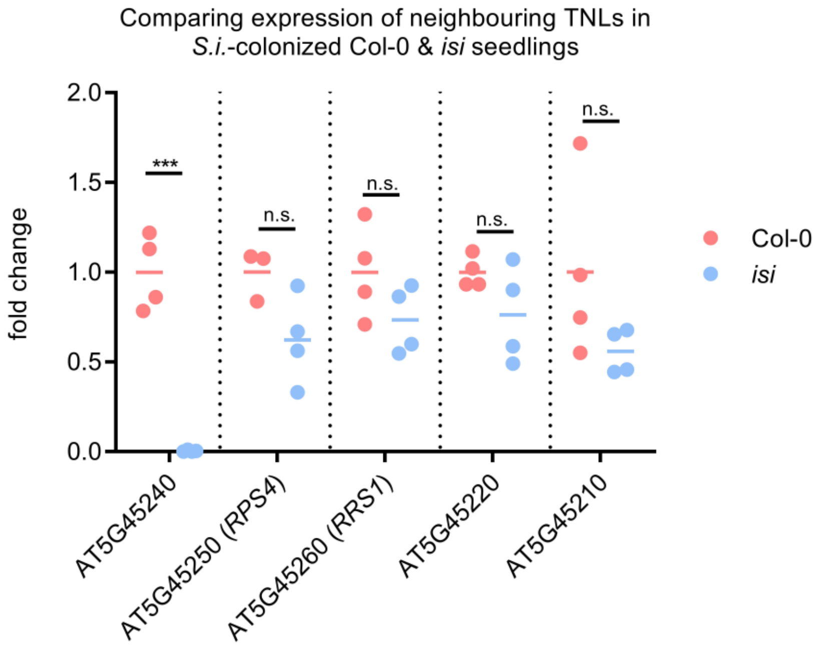


**Fig. S22: Gene locus of NLR gene and expression during *S. indica* colonization**

A) Relative expression of TNL genes of the *ISI* locus in Arabidopsis seedlings at different time points of *S. indica* colonization. Expression values relative to mock-treated seedlings at 1 dpi were calculated using the ΔΔCt-method. Dots represent three independent biological replicates while bars represent the average. The data is presented on a log2 scale. Asterisks represent significant difference to the Mock-treated sample analyzed by Student's t-test. $p < 0.05$ (*).

B) Relative expression of TNL genes of the NLR locus in Arabidopsis seedlings at 8 days after germination in *S. indica* presence on 1/10 PNM medium. Expression values relative to Col-0 seedlings were calculated using the ΔΔCt-method. Dots represent three independent biological replicates while bars represent the average. Asterisks represent significant difference to the mock-treated sample analyzed by Student's t-test. $p < 0.05$ (*).

### *isi* seedlings show reduced cell death after dAdo treatment

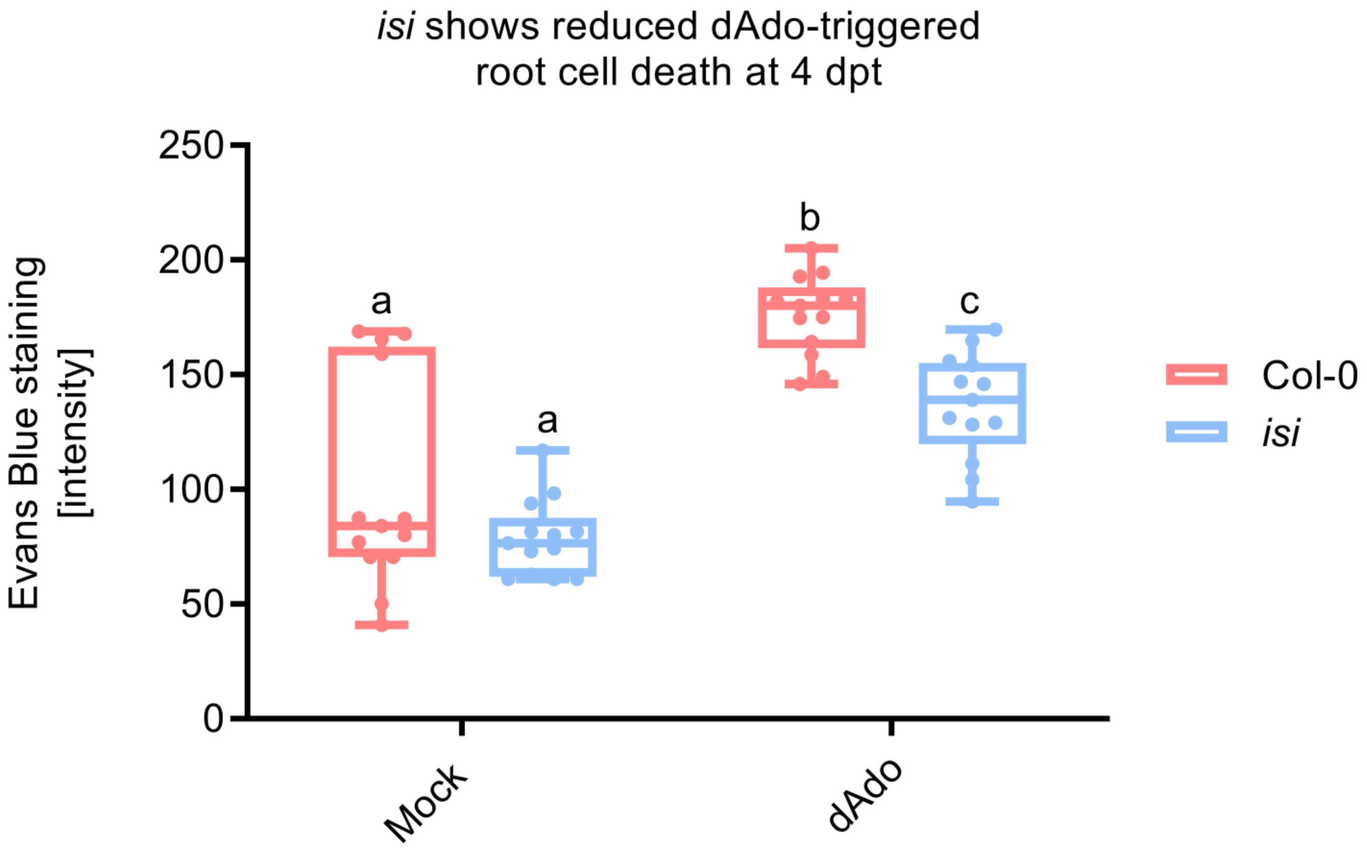


**Fig. S23: Evans Blue staining of dAdo-treated *A. thaliana isi* KO line**
Quantification of root cell death in Col-0 and *isi* root tips at 4 days after dAdo-treatment. Cell death was evaluated using Evans Blue staining. Boxplots represent the data obtained from 13 biological replicates. Different letters indicate significant differences determined with a two-way ANOVA with post-hoc Tukey HSD test ($p < 0.05$). The experiment was repeated independently 2 times with similar results.

### *nlr* but not *ent3* seedlings show loss of *S. indica*-induced growth promotion

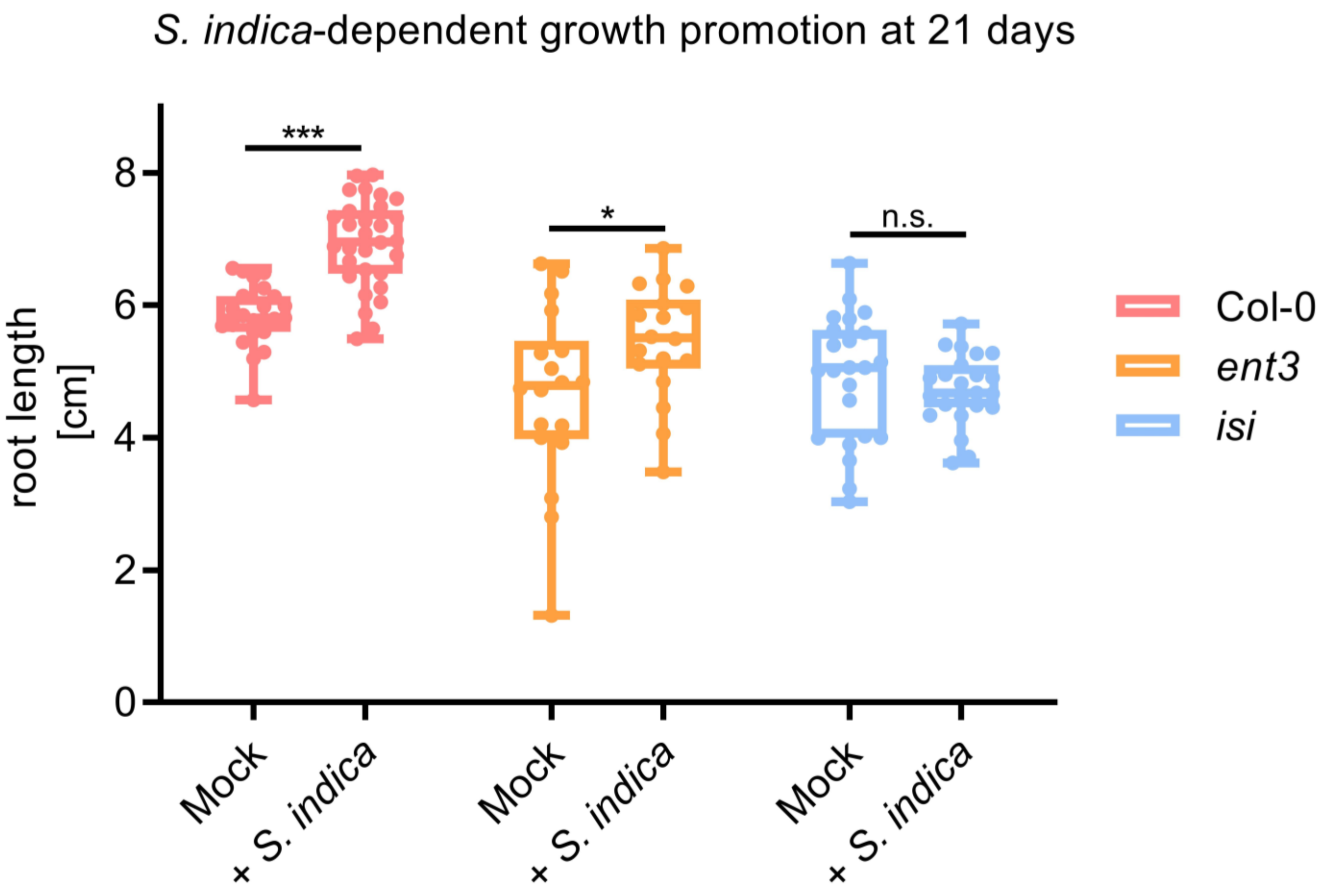


**Fig. S24: Root growth promotion in Arabidopsis seedlings**
**A)** Quantification of root length in Col-0, *ent3* and *isi* knockout lines at 21 days after germination. Seeds were treated with *S. indica* spores or a Mock control. Root length was evaluated from scans of the plates with Fiji (ImageJ). Boxplots represent the data obtained from 18-23 biological replicates. Asterisks represent significant difference to the Mock-treated sample analyzed by Student's t-test. $p < 0.05$ (*).

### Expression of NLR proteins during *S. indica* colonization

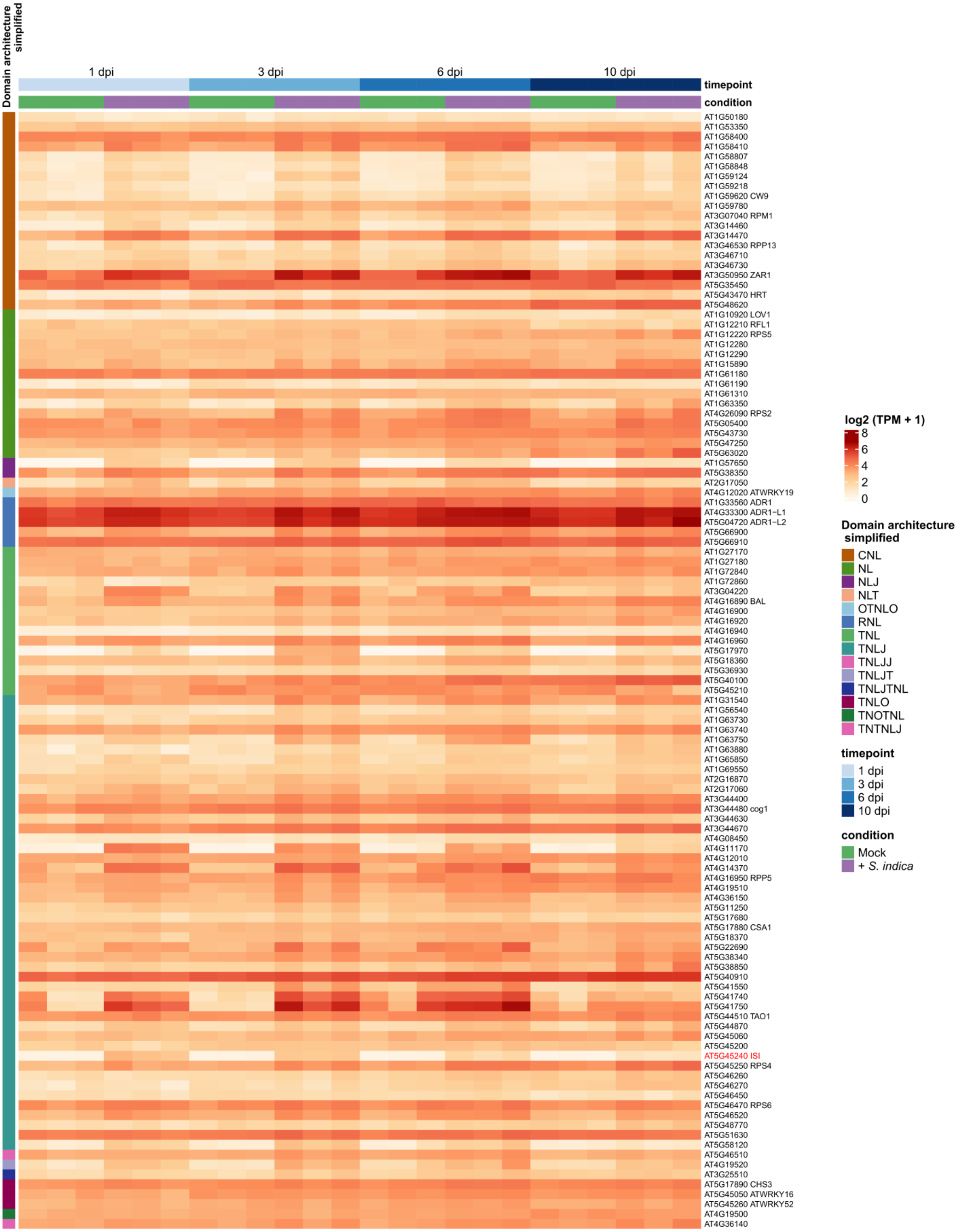

### Expression of NLR protein during *S. indica* colonization

**Fig. S25: Expression of different domain NLR proteins during *S. indica* colonization**

Gene locus of NLR gene and expression during *S. indica* colonization. The heatmap shows the expression values of *A. thaliana* NLR genes with at least NB-ARC and LRR (NL) domains in *A. thaliana* root samples. Specifically, *A. thaliana* plants were inoculated with Mock or *S. indica*. Roots samples were collected at 1 dpi, 3dpi, 6 dpi and 10 dpi. For each sample, stranded RNASeq libraries were generated and quantified by qPCR. RNA-seq libraries production and the sequencing were performed at the U.S. Department of Energy Joint Genome Institute under a project proposal (Proposal ID: 505829) (Zuccaro and Langen, 2020). The raw reads were filtered and trimmed using the JGI QC pipeline. Filtered reads from each library were aligned to the Arabidopsis TAIR9 reference genome using HISAT2 version 2.2.0. The gene counts were generated using featureCounts. NLR genes were identified in Arabidopsis TAIR10 proteins sequences using the software NLRtracker (Kourelis et al., 2021) which also provides the domain architecture of NLR proteins. Based on NLRtracker annotation, NLR genes with at least NB-ARC (N) and LRR (L) domains were selected. Next, genes with at least NL domains with an average TPM value > 1 TPM across all samples were selected. The log2 transformed TPM values of selected Arabidopsis NLR genes are shown in the heatmap generated using ComplexHeatmap package. In addition, the annotation of the protein domains annotation of selected NLR genes provided by NLRtracker are shown: TIR domain (T), NB-ARC domain (N), coiled-coil domain (CC), LRR domain (L), Other domain (O), C-JID domain (J), RPW8-type CC domain (R).

Project: Zuccaro, A., & Langen, G. (2020). Host-specific regulation of effector gene expression in mutualistic root endophytic fungi (Proposal ID: 505829). JGI Award DOI: 10.46936/10.25585/60001292.

### *S. indica* colonization of *A. thaliana* root tip

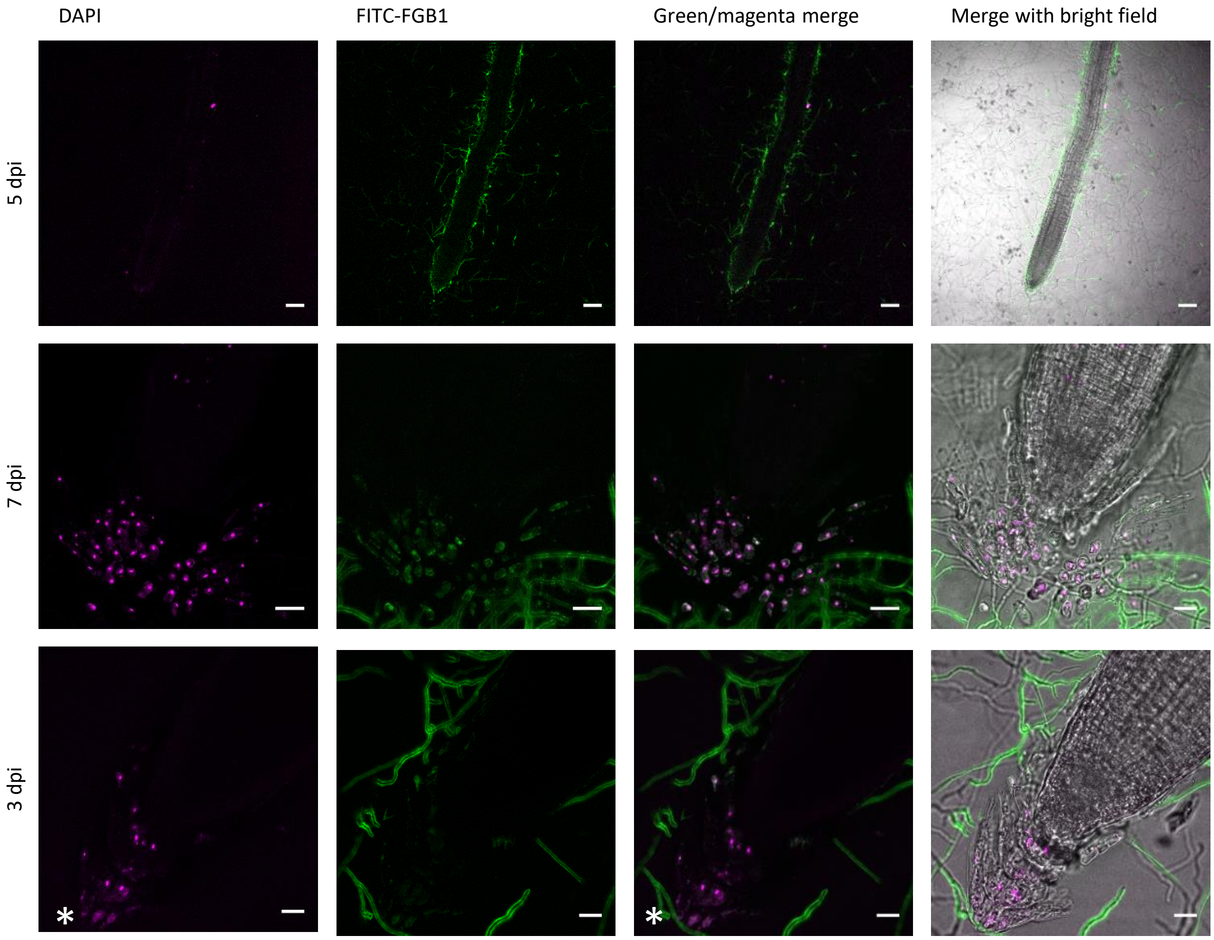


**Fig. S26: *S. indica* growth around *A. thaliana* root tip in Root Extracellular Traps (RET).**

CLSM live cell imaging of *S. Indica*-colonized *A. thaliana* roots. Nuclei stained with DAPI (magenta) and fungal cell wall and matrix with FITC-FGB1 (green). Roots were directly stained and imaged on plate to avoid removing the border-like cells (BLC) and RET. Nuclei of border-like cells (BLC) are visible in magenta. *S. indica* is growing around the root tip and among the BLC/RET of the root cap. Nuclei of the outer BLC layer are often blurred (asterisk). Bars = 100 µm (Row 1), Bars = 20 µm (Row 2-3)
